# Recombination between heterologous human acrocentric chromosomes

**DOI:** 10.1101/2022.08.15.504037

**Authors:** Andrea Guarracino, Silvia Buonaiuto, Leonardo Gomes de Lima, Tamara Potapova, Arang Rhie, Sergey Koren, Boris Rubinstein, Christian Fischer, Human Pangenome Reference Consortium, Jennifer L. Gerton, Adam M. Phillippy, Vincenza Colonna, Erik Garrison

## Abstract

The short arms of the human acrocentric chromosomes 13, 14, 15, 21, and 22 share large homologous regions, including the ribosomal DNA repeats and extended segmental duplications (Floutsakou et al. 2013; van Sluis et al. 2019). While the complete assembly of these regions in the Telomere-to-Telomere consortium’s CHM13 provided a model of their homology (Nurk et al. 2022), it remained unclear if these patterns were ancestral or maintained by ongoing recombination exchange. Here, we show that acrocentric chromosomes contain pseudo-homologous regions (PHRs) indicative of recombination between non-homologs. Considering an all-to-all comparison of the high-quality human pangenome from the Human Pangenome Reference Consortium (HPRC) (Liao et al. 2022), we find that contigs from all of the acrocentric short arms form a community similar to those formed by single chromosomes or the sex chromosome pair. A variation graph (Garrison et al. 2018) constructed from centromere-spanning acrocentric contigs indicates the presence of regions where most contigs appear nearly identical between heterologous CHM13 acrocentrics. Except on chromosome 15, we observe faster decay of linkage disequilibrium in the PHRs than in the corresponding short and long arms, indicating higher rates of recombination (N. Li and Stephens 2003; Huttley et al. 1999). The PHRs include sequences previously shown to lie at the breakpoint of Robertsonian translocations (Jarmuz-Szymczak et al. 2014), and we show that their arrangement is compatible with crossover in inverted duplications on chromosomes 13, 14, and 21. The ubiquity of signals of recombination between heterologous chromosomes seen in the HPRC draft pangenome’s acrocentric assemblies suggests that these shared sequences form the basis for recurrent Robertsonian translocations, providing sequence and population-based confirmation of hypotheses first developed cytogenetically fifty years ago (Hamerton et al. 1975).

## Introduction

Although the human reference genome is now 22 years old (International Human Genome Sequencing Consortium 2001), fundamental limitations of the bacterial artificial chromosome libraries on which it was built prevented its completion. Incomplete regions amount to 8% of the Genome Reference Consortium’s Human Build 38, and include heterochromatic regions in the centromeres and the short arms of the acrocentrics. Advances in long-read DNA sequencing recently enabled the creation of a complete reference assembly—the Telomere-to-Telomere consortium’s CHM13 (T2T-CHM13) (Nurk et al. 2022)—from a homozygous human cell line, providing a reference system for these regions for the first time and exposing them to potential study. In parallel, our ongoing work in the Human Pangenome Reference Consortium (HPRC) has yielded 94 haplotype-resolved assemblies for human cell lines (HPRCy1), based on the same Pacific Biosciences circular consensus (HiFi) sequencing that forms the foundation of T2T-CHM13 (Liao et al. 2022). These resources enable us to characterize patterns of variation in these previously-invisible regions for the first time. In this work, we study variation in the largest non-centromeric regions made visible in T2T-CHM13 and HPRCy1: those between the centromere and the ribosomal DNA (rDNA) on the short arms of the acrocentrics (SAACs), where the most common human translocations events occur, Robertsonian translocations (ROBs) (Mack and Swisshelm 2013).

Most human chromosomes (N=18) are metacentric, with the centromere found in a median position between short (“p”) and long (“q”) arms, while 5 are acrocentric, and feature one arm that is significantly shorter than the other. The short arms of human acrocentric chromosomes (13p, 14p, 15p, 21p, and 22p) host the nucleolus organizing regions (NORs), the genomic segments that harbor ribosomal DNA (rDNA) genes and that give rise to the interphase nucleoli (McClintock 1934; Spinner 2013; Lindström et al. 2018). Due to their repetitive nature, rDNA repeat arrays facilitate intramolecular recombination (Kobayashi 2011). rDNA repeats incur double-strand breaks at a high rate due to transcription-replication conflicts (Lindström et al. 2018). Moreover, rDNA from multiple acrocentric chromosomes can be co-located in nucleoli during interphase, and multiple acrocentric chromosomes often colocalize to a single nucleolus during pachytene (Holm and Rasmussen 1977), when chromosomes synapse and recombine. As it causes them to occupy the same constrained physical space, the positioning of rDNA-adjacent sequences at the nucleolar periphery could be a driver of genetic exchange between heterologous chromosomes (**Supplementary Note 1**). Based on estimates from (Holm and Rasmussen 1977) the probability of two NOR-adjacent regions being colocalized is 120,000 times larger than colocalization in a human spermatocyte nucleus. In line with this, distal and proximal sequences to rDNA repeat arrays are conserved among the acrocentric chromosomes, suggesting that recombination homogenizes them (Floutsakou et al. 2013; van Sluis et al. 2019). Experimental and sequence-based evidence indicates the presence of a common subfamily of alpha satellite DNA shared by 13/21 and 14/22 pairs of acrocentric chromosomes that provides evidence for an evolutionary process consistent with recombination between heterologous chromosomes (Choo et al. 1988). Furthermore, Robertsonian translocations, which occur in 1/800 births, are most common between chromosomes 13 and 14 (~75%), and 14 and 21 (~10%), begging the question of the underlying sequences and recombination processes that drive them (Jarmuz-Szymczak et al. 2014).

The T2T-CHM13 reference fully resolves the genomic structure of the short arms of acrocentric chromosomes for the first time, confirming their strong similarity and providing a complete view of the homologies in this single genome (Nurk et al. 2022). However, T2T-CHM13 does not provide information on how SAACs vary in the human population and additional genomes are needed to understand if T2T-CHM13’s representation is typical. Notably, alignments of HPRCy1 assemblies to T2T-CHM13 revealed individual contigs with optimal alignments to multiple CHM13 acrocentric chromosomes, suggesting possible translocations (Liao et al. 2022). This analysis was necessarily relative to only a single frame of reference used as target in alignment, leaving open questions as to the relationships between pairs of HPRCy1 haplotypes. A complete study of this region thus requires improvements in both sequence assembly and pangenome analysis to enable an unbiased assessment of its structure and variation in the population. Here, we combine T2T-CHM13 and HPRCy1 assemblies in a reference-free pangenome variation graph (PVG) model of the SAACs. Using this model and other symmetric analyses of T2T-CHM13 and the HPRCy1 assemblies, we establish the first coherent model of population-scale variation in the short arms of the human acrocentrics.

## Results

### Chromosome community detection

We sought to study the chromosome groupings implied by the homologies found in all 94 of the HPRCy1 pangenome’s assemblies. We used homology mapping to build a reference-free model of the HPRCy1 pangenome, represented as a mapping graph with nodes as contigs and edges as mappings between them. The graph was built using chains of 50 kbp seeds of 95% average nucleotide identity—features that we expect support homologous recombination (Peng et al. 2015)—with up to 93 alternative mappings allowed per contig (**Materials and Methods**). After applying this process to all 38,325 HPRCy1 contigs and narrowing our focus to only mappings involving contigs at least 1 Mbps long, we built a reduced mapping graph by selecting the best 3 mappings per contig segment and labeling each contig with its reference-relative assignment (**Figure 1A**). This simplified graph showed clusters that generally matched our expectations of higher similarity between certain chromosomes (**Figure 1B**) (Devilee et al. 1986; Jørgensen, Bostock, and Bak 1987; Greig, Warburton, and Willard 1993; Jørgensen et al. 1988). For a more quantitatively rigorous interpretation, we used a community detection algorithm (Traag, Waltman, and van Eck 2019) to divide the full mapping graph into 31 communities (**Supplementary Table 1**). These communities were consistent with our expectations based on mapping the contigs to reference chromosomes T2T-CHM13 and GRCh38 and known patterns of similarity between chromosomes. We found that the SAACs’ community contained the most distinct chromosomes and the most contigs (**Figure 1C-1D**). Many contigs from the pseudoautosomal regions (PARs) and X-transposed regions of chromosome X and all of those from chromosome Y (Helena Mangs and Morris 2007; Cotter, Brotman, and Wilson Sayres 2016) formed one community, while others from the short arm of chromosome X, including all of those from evolutionary strata 4 and 5 (Ross et al. 2005), formed another (**Supplementary Figure 1**). A few additional communities were identified that did not correspond to individual chromosomes, but typically represent single chromosome arms.

**Figure 1.**
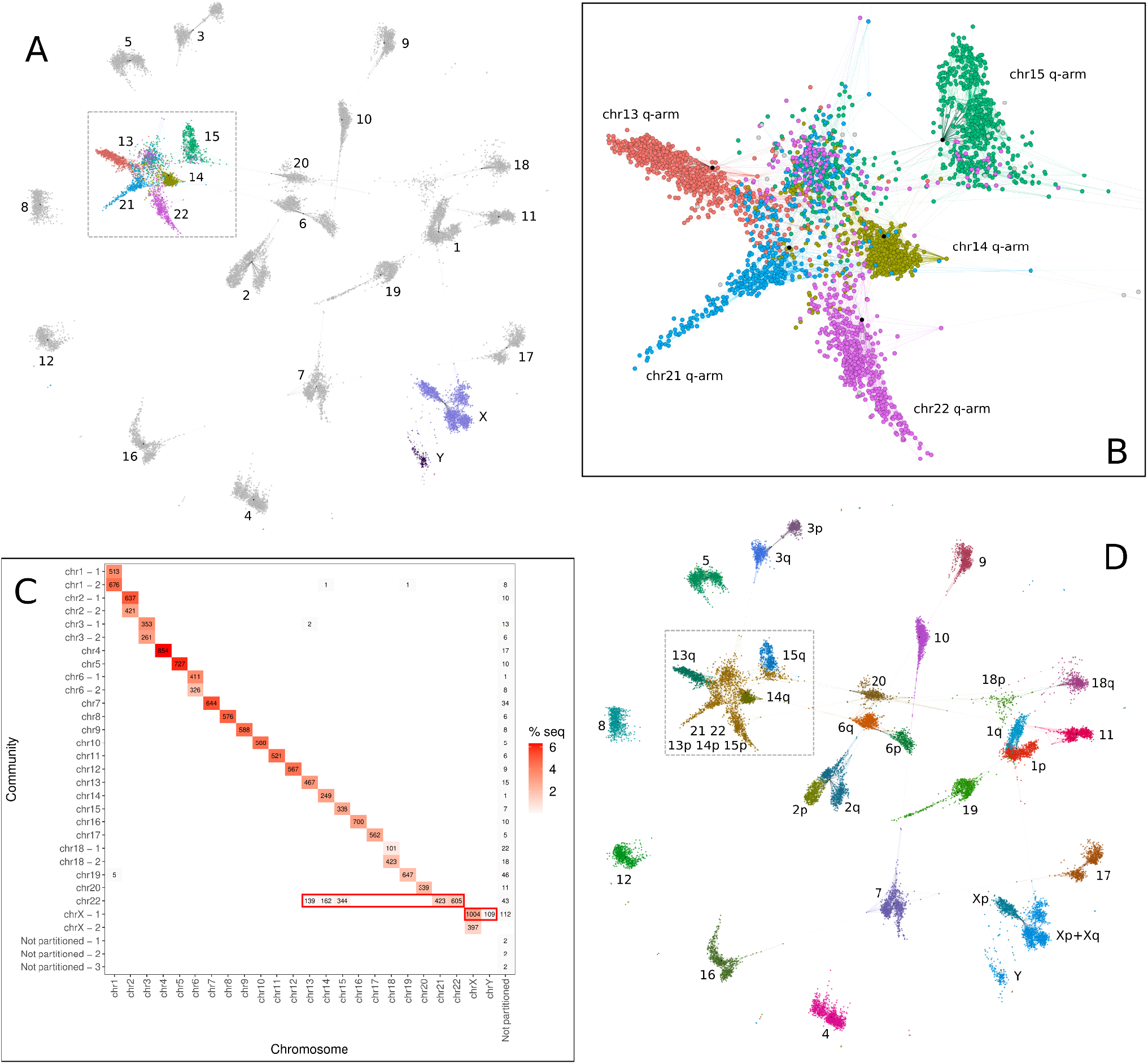
Community detection in the HPRCy1 pangenome. (**A**) The reduced all-to-all mapping graph of HPRCy1 versus itself, with contigs represented as nodes and mappings as edges, rendered in Gephi (Bastian, Heymann, and Jacomy 2009). Colors indicate the acrocentric or sex chromosome to which each contig was assigned by competitive mapping against T2T-CHM13 and GRCh38, with text labels indicating the chromosome for each visual cluster. (**B**) A close-up of the region indicated in A and D containing virtually all contigs matching acrocentric chromosomes. (**C**) Results of community assignment on the mapping graph. On the x-axis the chromosome to which contigs belong based on competitive mapping to T2T-CHM13 and GRCh38, while the y-axis indicates the community, which is named by the chromosome that contributes the largest number of contigs to it. In each square, there is the number of contigs that belong to a specific chromosome and community. For each chromosome-community combination, the red gradient indicates the percentage of out of the total assembly sequence present in the set. The sex and acrocentric chromosomes respectively participate in the only clusters (highlighted with the red rectangles) which mix many (N>100) contigs belonging to different chromosomes. (**D**) Coloring the reduced homology mapping graph in A with community assignments (colors do not correlate with panel A or B). The short arms of 13, 14, 15, and all of 21 and 22 are involved in one community, as are Y and most of X.

### An all-acrocentric pangenome graph

We constructed a pangenome graph from acrocentric contigs in the HPRCy1 draft pangenome to evaluate the hypothesis that heterologous SAACs recombine. We first collected long HPRCy1 contigs that span the acrocentric centromeres and can be assigned to specific acrocentric chromosomes (**Supplementary Figure 2**). We then used the PanGenome Graph Builder (PGGB) (Garrison et al. 2022) to construct a single pangenome variation graph (PVG) from these contigs (**Materials and Methods**). PVG nodes represent sequences and edges indicate when concatenations of the nodes they connect occur in the contigs represented by the graph (Paten et al. 2017). By relating pangenome sequences to the graph as paths of nodes (Garrison et al. 2018), PVGs support base-level analysis of variation and homology between genomes (Eizenga et al. 2020; Hickey et al. 2020; Sirén et al. 2021; Liao et al. 2022; Guarracino et al. 2022). PGGB’s symmetric all-to-all alignment (Marco-Sola et al. 2022) and graph induction (Garrison and Guarracino 2022) avoid sources of bias like reference choice and genome inclusion order that affect progressive PVG construction methods (H. Li, Feng, and Chu 2020; Armstrong et al. 2020). For cross-validation of our results, we additionally include two assemblies of HG002 in the PVG: HG002-HPRCy1 (Liao et al. 2022), obtained from HiFi reads, and HG002-Verkko, a T2T diploid assembly constructed from both HiFi and Oxford Nanopore Technologies (ONT) reads as described in (**Materials and Methods**) and (Rautiainen et al. 2022).

The resulting acrocentric PVG (acro-PVG) presents structures that echo those observed in T2T-CHM13 and the community structure of the homology mapping graph (**Figure 2A and Supplementary Files 1 and 2**). In more detail, the main connected component including all chromosomes presented a tangled region, anchored at the ribosomal DNA repeats, but also extending towards the centromere-proximal end of the short arms. The alpha satellite higher-order repeat (HOR) arrays in the centromeres of chromosomes 13/21 and 14/22 pairs shared high similarity within each pair (Greig, Warburton, and Willard 1993; Jørgensen et al. 1988), leading to collapsed motifs in the graph (**Figure 2B**). 13/21 and 14/22 diverge in centromere-proximal regions of the q-arms. Furthermore, a region in the pangenome graph centered on the GC-rich SST1 array was present in a single copy in chromosomes 13, 14, and 21, indicating a high degree of similarity of the samples in those regions (**Figure 2C** and **Supplementary Figure 3**). This is compatible with the frequent involvement of these regions of chromosomes 13, 14, and 21 in Robertsonian translocations (Sullivan et al. 1996; Cheng and Naluai-Cecchini 2004; Jarmuz-Szymczak et al. 2014). The SST1 elements in the segmentally duplicated region, also known as NBL2 (Nishiyama et al. 2005), are GC-rich 1.4-2.4 kb long sequences arranged in tandem clusters (Epstein et al. 1987), located throughout the genome including near the centromeres of the short arm of the acrocentric chromosomes 13, 14, 21 (Hoyt et al. 2022). The SST1 array size is variable in the human population (Tremblay et al. 2010) and its methylation status is clinically relevant to cancer (Samuelsson et al. 2017; González et al. 2021). SST1 repeats on chromosomes 13, 14, and 21 in T2T-CHM13 are highly similar to each other (Hoyt et al. 2022), consistent with homogenization via recombination. All the graph motifs described in the acro-PVG were also confirmed by building a pangenome graph without including the T2T-CHM13 and GRCh38 references (**Supplementary Figures 4 and 5**), indicating that the observed structure is independent of the reference assemblies.

**Figure 2.**
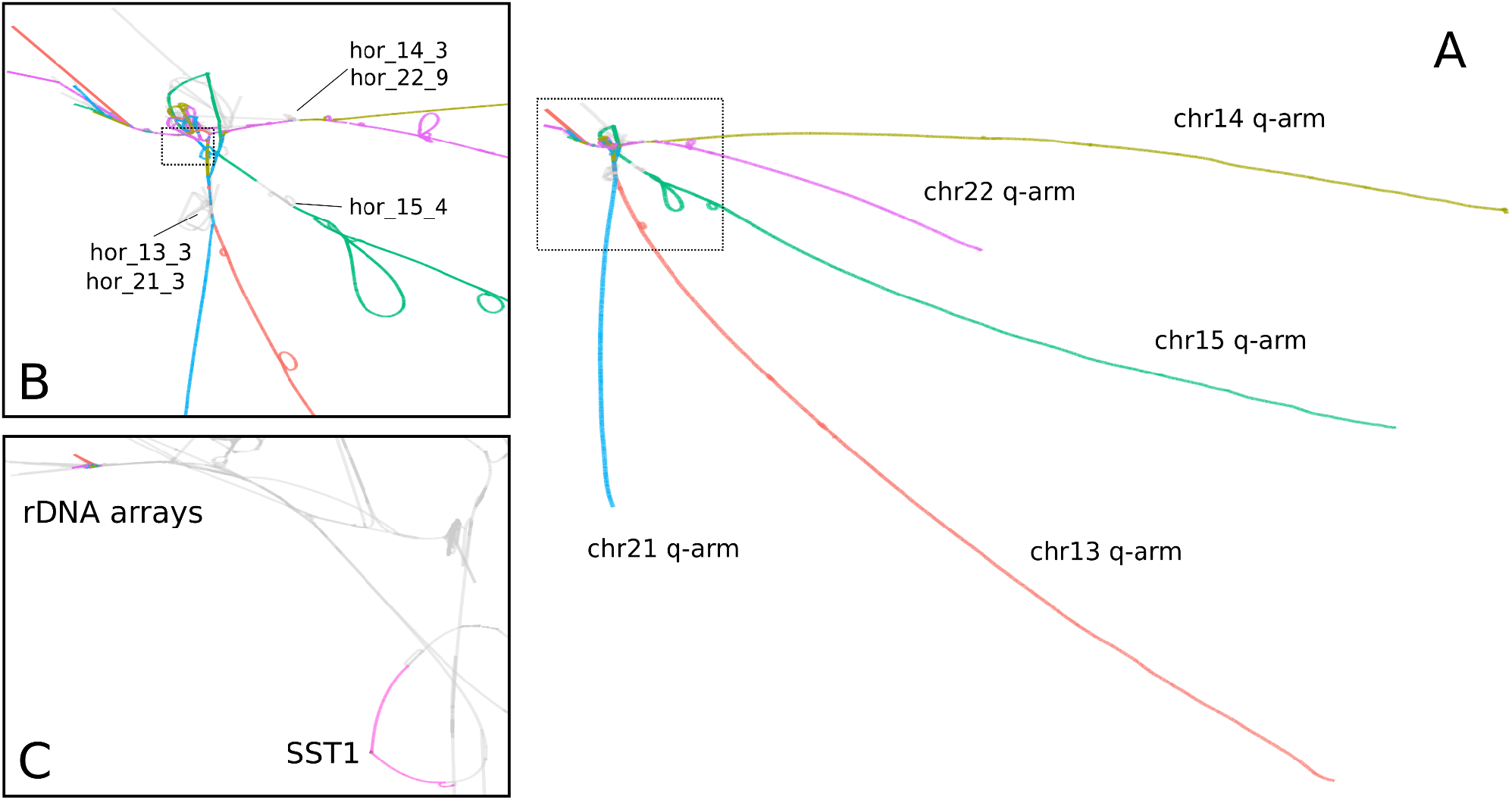
The HPRCy1 acrocentric PVG. We apply GFAESTUS (Fischer and Garrison 2022) to visualize the 2D layout generated by ODGI (Guarracino et al. 2022) in the PGGB pipeline (Garrison et al. 2022). This renders sequences and chains of small variants as linear structures, while repeats caused by segmental duplications, inversions, and other structural variants tend to form loops. (**A**) The major component of the graph, shown with nodes in T2T-CHM13 chromosomes labeled using colors also used in other figures. The acrocentric q-arms are almost completely separated, while the p-arms unite in a tangle adjacent to the rDNA array. (**B**) A close-up of the SAACs’ junction, showing the separation of chromosome 15’s centromeric high-order repeats (*hor_l5_4*) from the other chromosomes, while 13/21, and 14/22 share substantial homology in their arrays that causes them to collapse in the PVG. A few assemblies span the rDNA array into its distal junction, which presents as a single homologous region across all chromosomes (van Sluis et al. 2019), and then fray into diverse sequences visible as tips in the top left. (**C**) Zooming further, we focus on the segmentally duplicated core centered in the SST1 array and the rDNA arrays of the various chromosomes. In this panel, we label the rDNA arrays and SST1 subsequences in T2T-CHM13. The highlighted region around the SST1 array is in the same orientation on T2T-CHM13’s 13p11.2 and 21p11.2, and inverted on 14p11.2; these three regions show a pairwise identity above 99% (Nurk et al. 2022).

### Genomic information flow between heterologous SAACs

The acro-PVG provides a representation of the multiple alignment of SAACs found in the human population. In the acro-PVG, we observe many regions in the graph where multiple T2T-CHM13 chromosomes are aligned together. We expect these regions to potentially support homologous recombination, which largely depends on sequence homology and physical proximity (Kaniecki, De Tullio, and Greene 2018) both of which are common among heterologous SAACs (Henderson, Warburton, and Atwood 1973; Holm and Rasmussen 1977).

### HPRCy1 contigs are homology mosaics

We sought to test the hypothesis that homologous regions of the SAACs feature ongoing sequence exchange by looking for regions in the acrocentric PVG where individual contigs are best-described as a mosaic of diverse T2T-CHM13 acrocentric chromosomes. We derived a pairwise alignment from the acro-PVG through *untangling* (Guarracino et al. 2022), a process that projects the graph into an alignment between a set of query (HPRCy1-acro) and reference (T2T-CHM13) sequences, jointly considering all possible alignments represented by the pangenome graph. The untangling of the acro-PVG against multiple T2T-CHM13 chromosome reference sequences simultaneously shows the best match of segments within contigs to multiple reference chromosomes.

The hypothesis of recombination between heterologous acrocentric chromosomes implies that the HPRCy1-acro contigs untangled from the acro-PVG will be a mosaic of diverse acrocentric chromosomes in the regions undergoing homologous recombination. The same would not be true for flanking regions that should map to one specific chromosome.

We queried the PVG (Guarracino et al. 2022) to obtain a mapping from segments of all PVG paths onto T2T-CHM13. This segments the graph, and for each HPRCy1 contig (query) subpath through each graph segment, we find the most-similar reference segment (**Supplementary Figure 6**). To reduce the possibility of error, we focused the alignment projection only on the confidently-assembled regions of the HPRCy1-acro contigs (Liao et al. 2022) (**Materials and Methods**) and we filtered the mappings to retain only those at greater than 90% estimated identity, removing a total of 1.17 Gbps, or 2.52% of the total SAAC contig segments (**Supplementary Table 2 and Supplementary Figures 7-11**).

For a reference-relative interpretation of the results, we anchored the contigs to the single T2T-CHM13 reference chromosomes to which the q-arm maps (**Materials and Methods**), providing a reference-relative positioning of contigs in the PVG. We find that the q-arm of each contig maps to a single chromosome, while the p-arm is a mosaic of segments mapping to several acrocentric chromosomes (**Figure 3A-B)**. Results for all the acrocentric chromosomes are shown in **Supplementary Figures 13-17**.

**Figure 3.**
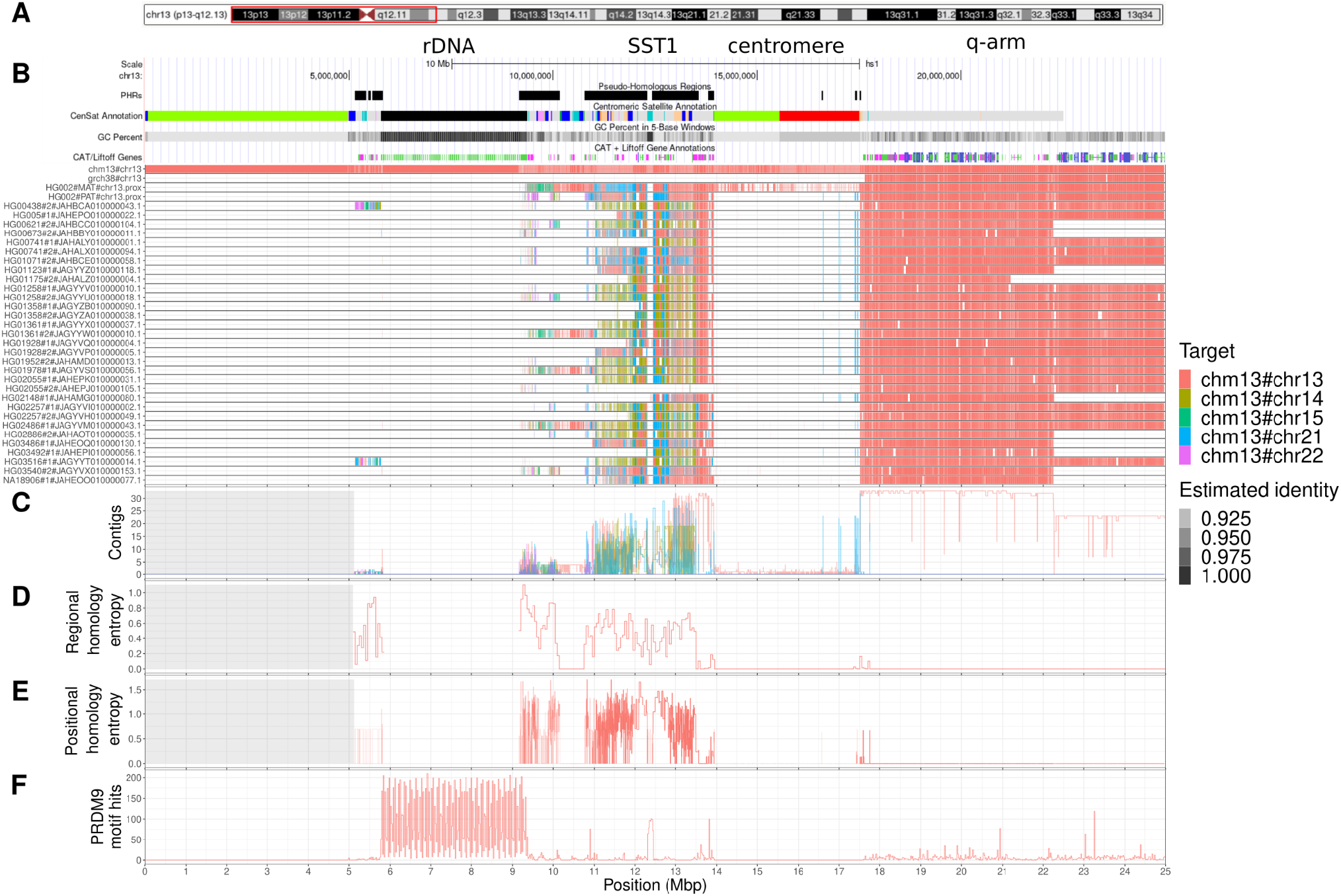
Characteristics of the pseudo-homologous regions of acrocentric chromosomes. **(A)** We focus on the first 25 Mbp of chromosome 13, shown here as a red box over T2T-CHM13 cytobands. **(B)** Pseudo-homologous regions (**PHRs**) are highlighted relative to T2T-CHM13 genome annotations for repeats, GC percentage, and genes. Above, we indicate regions of interest described in the main text: rDNA, SST1 array, centromere, and q-arm. Below, we show T2T-CHM13-relative homology mosaics for each chromosome 13 matched contig from HPRCy1-acro, with the most-similar reference chromosome at each region shown using the given colors (**Target**). **(C)** Aggregated untangle results in the SAACs. For each acrocentric chromosome, we show the count of its HPRCy1 q-arm-anchored contigs mapping itself and all other acrocentrics (**Contigs**), **(D)** as well as the regional (50kbp) untangle entropy metric (**Regional homology entropy**) computed over the contigs’ T2T-CHM13-relative untanglings. **(E)** By considering the multiple untangling of each HPRCy1-acro contig, we develop a point-wise metric that captures diversity in T2T-CHM13-relative homology patterns (**Positional homology entropy**), leading to our definition of the PHRs. **(F)** The patterns of homology mosaicism suggest ongoing recombination exchange in the SAACs. A scan over T2T-CHM13 reveals that both the rDNA and SST1 array units are enriched for PRDM9 binding motifs, and thus may host frequent double stranded breaks during meiosis. In (C-E) a gray background indicates regions with missing data due to the lack of non-T2T-CHM13 contigs. We provide the Centromeric Satellite Annotation (CenSat Annotation) track legend in **Supplementary Figure 12**.

We cross-validated homology mosaic patterns by comparing the reference-relative anchored untanglings of HG002-Verkko to HG002-HiFi, obtaining a 87.45% concordance rate on the SAACs, and a 99.93% concordance rate in the acrocentric q-arms (**Materials and Methods**). Although HG002-HiFi contains only one contig that would meet our HPRCy1 contiguity requirements, we observe broadly concordant patterns in the two assemblies (**Supplementary Figures 18-22**) and visually confirm patterns—such as those between 13p and 21p—which are seen in many HPRCy1 assemblies (**Supplementary Figure 18**).

### Homology mosaicism increases across acrocentric short arms

We counted the number of contigs that best-match each one of the T2T-CHM13 acrocentrics within the PVG (**Figure 3C**). On the q-arm, all contigs best-match their homologous T2T-CHM13 chromosome, agreeing with the observed structure of the PVG (**Figure 2A**). However, as we approach the centromere from the q-arm, we observe regions of homology between 13 and 21 (**Figure 3C and Supplementary Figures 13B and 16B**) and between 14 and 22 (**Supplementary Figures 14B and 17B**). In contrast, homology with other acrocentrics begins closer to the rDNA in chromosome 15 (**Supplementary Figure 15B**), corroborating the pattern observed in the PVG topology (**Figure 2B**).

Although the 13+21 and 14+22 HOR arrays are both collapsed in the PVG (**Figure 2B**), we observe sparse >90% identity mappings within the centromeres (**Figure 3B and Supplementary Figures 13A-17A**). This is consistent with other reports of high divergence within centromeric satellites (Logsdon et al. 2021). HPRCy1 contigs anchored on the q-arms of chromosomes 13, 14, and 21 share a segmental duplication, or homologous region, centered on the SST1 array (**Figure 3B**), in line with what is seen in the pangenome graph topology (**Figure 2C and Supplementary Figures 3 and 5**). Furthermore, as in T2T-CHM13, this region is in the same orientation on chromosomes 13 and 21, but inverted on chromosome 14 (**Supplementary Figures 23-25**). All chromosomes provide similarly good matches for contigs in the regions immediately proximal and distal to the rDNA. However, this is supported by relatively few q-arm-anchored contigs (N=9) which purport to cross the rDNA—loci which we do not expect to assemble correctly using current sequencing approaches (Nurk et al. 2022).

To assess the homology between the acrocentrics, we developed a metric that captures the degree of disorder in the untangling of HPRCy1-acro contigs over 50 kbp regions of T2T-CHM13 (**Materials and Methods**). This metric, *regional homology entropy*, is greater than 0 in regions where contigs match multiple T2T-CHM13 chromosomes—a pattern indicative of recombination. We find that regional homology entropy increases as we progress over each short arm and reaches a maximum immediately on the proximal flanks of the rDNA arrays (**Figure 3D**). We observed an equivalent increase of regional homology entropy in the PARs on chromosomes X and Y (**Supplementary Figure 26**), which are known to actively recombine.

### Delineating the acrocentric pseudo-homologous regions

Our analyses suggest that regions of near-identity between multiple T2T-CHM13 chromosomes are capable of supporting large-scale homologous recombination. To study the boundaries of these regions, we derived a *multiple* untangling, which orders, by identity, multiple T2T-CHM13 matches for every contig segment (**Supplementary Figures 27-31**). The order of T2T-CHM13 hits captures the leaf order of a HPRCy1 contig-rooted phylogeny (Kinene et al. 2016). Differences in chromosome-relative phylogenies across haplotypes indicate different evolutionary histories and imply ongoing recombination (Arenas 2013), which leads to checkerboard patterns in the multiple untangling plots (**Figure 4C**). To delineate regions where heterologous chromosomes are likely to recombine, we computed *positional homology entropy*—a measure of the diversity of reference-relative phylogenies—for each position in T2T-CHM13 (**Figure 3E and Supplementary Figure 32**). We consider regions with positional homology entropy greater than 0 over more than 30 kbps to be candidates for ongoing recombination (**Materials and Methods**).

**Figure 4.**
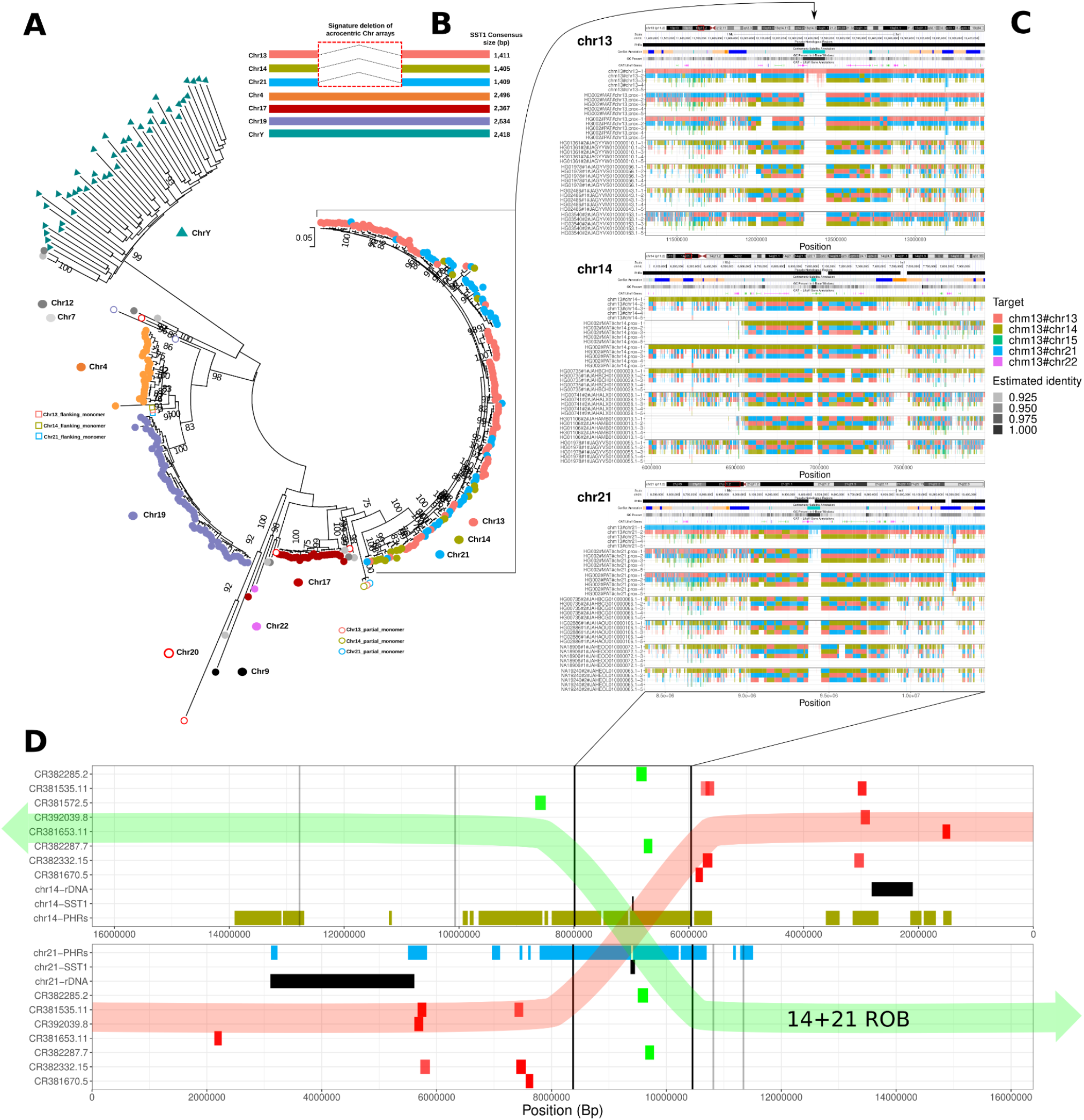
Pseudo-homologous regions of chromosomes 13, 14, and 21 centered on the SST1 array. **(A)** Maximum Likelihood phylogenetic analysis of SST1 full-length elements indicates a recent homogenization process of acrocentric arrays. Colored circles by chromosome labels indicate individual monomers retrieved from the T2T-CHM13 assembly. Colored triangles indicate SST1 full-length monomers retrieved from HG002 chrY assembly (Rhie et al. 2022). Partial or chimeric monomers flanking chr13/14/21 arrays (located ~250 kb from the main array) are labeled as open circles and squares, respectively, according to each chromosome color label. (**B**) Schematic representation of SST1 consensus alignments indicating a characteristic deletion present only in the SST1 unit from arrays on chr13/14/21. **(C)** Multiple untangling of T2T-CHM13, HG002-Verkko haplotypes, and HPRCy1-acro contigs versus T2T-CHM13. Three of the five acrocentric chromosomes are represented (from top: 13, 14, and 21), with contigs and chromosome assemblies below. Transparency shows the estimated identity of the mappings. We display all mappings above 90% estimated pairwise identity. To allow the display of simultaneous hits to all acrocentrics, each grouping shows the first 5 best alternative mappings. SST1 arrays described in (A) lie at the center of a PHR that displays checkerboard patterns indicative of recombination between heterologous acrocentrics (black arrows link the SST1 arrays in all panels). These patterns are less common on chromosomes 15 and 22 (**Supplementary Figures 29 and 31**). Panel (**D**) relates the PHRs on T2T-CHM13 (yellow and light blue) to BACs localized cytogenetically by (Jarmuz-Szymczak et al. 2014) to recurrent 14+21 Robertsonian translocation breakpoints. BACs shown in green are found in dicentric Robertsonian chromosomes, while those in red are not. We display chromosome 14 in an inverted orientation aligned to 21 at the breakpoint region suggested by these experiments. In a transparent overlay, we propose a retained dicentric chromosome (“14+21 ROB”, green) and lost (red) products of the studied recurrent translocations.

These pseudo-homologous regions (PHRs) total 18.329 Mbp in length (**Supplementary File 3**) and differs by chromosome: chr13 = 4.53 Mbp (**Figure 3B**), chr14 = 6.48 Mbp (**Supplementary Figure 14A**), chr15 = 719.25 kbp (**Supplementary Figure 15A**), chr21 = 3.79 Mbp (**Supplementary Figure 16A**), and chr22 = 2.81 Mbp (**Supplementary Figure 17A**). We term them PHRs by analogy to the PARs of sex chromosomes, because these homology domains could allow non-homologous chromosomes to pair like homologous chromosomes. Notably, the chromosomes involved in the most common ROBs (13+14, 14+21) have larger PHRs, which could promote the recombination events that lead to these translocations. Supporting this, bacterial artificial chromosome (BAC) clones surrounding common recurrent ROB breakpoints (Jarmuz-Szymczak et al. 2014) map to T2T-CHM13 PHRs (**Supplementary Figure 33**). A genome-wide phylogenetic analysis of SST1 array elements indicates the expected pattern of concerted evolution by chromosome, but the repeats from chromosomes 13, 14, and 21 display a unique pattern of concerted evolution between chromosomes (**Figure 4A**), and furthermore, share a ~1.0 kb deletion relative to all other SST1 repeats (**Figure 4B and Supplementary Figure 34**), suggesting inter-array recombination similar to the surrounding non-satellite sequences of the PHRs (**Figure 4C**). We confirmed that patterns observed by fluorescent in-situ hybridization (FISH) of these BACs are compatible with breakpoints occuring in the PHRs centered at the SST1 array (**Figure 4D**).

To provide a positive control, we applied the same method to the sex chromosome PVG to identify their PHRs (chrX = 2.75 Mbp, chrY 2.73 Mbp; **Supplementary File 4 and Supplementary Figure 35**). These regions precisely match the established boundaries for the PARs and contain sparse hits in the X-transposed regions (XTRs), which would be compatible with reports of X-Y interchange in the XTR (Veerappa, Padakannaya, and Ramachandra 2013). Biallelic SNP calls from a whole-genome HPRCy1 graph released in our companion work (Liao et al. 2022) show that variant density in the PARs and the acrocentric p-arms is markedly higher than elsewhere in these chromosomes (**Supplementary Figure 36**), which is consistent with increased rates of recombination in these regions (Cotter, Brotman, and Wilson Sayres 2016).

In humans and many other mammals, the sequence specificities of the DNA-binding zinc finger protein PRDM9 regulate the formation of double stranded breaks that drive meiotic homolog synapsis and recombination (Paigen and Petkov 2018; Zickler and Kleckner 2015). We scanned T2T-CHM13 acrocentrics for ChIP-seq derived PRDM9 motifs (Altemose et al. 2017) (**Supplementary Figure 37 and Supplementary File 5**), finding that both the rDNA and SST1 arrays are enriched for PRDM9 motifs relative to the surrounding sequence (**Figure 3F and Supplementary Figures 13E-17E**). In contrast, we find virtually no PRDM9 motifs in the centromeres, where meiotic recombination is harmful and suppressed by diverse mechanisms (Nambiar and Smith 2016).

### Linkage disequilibrium decay in pseudo-homologous regions

To quantify the magnitude of putative recombination in the PHRs, we calculated the rate of the linkage disequilibrium (LD) decay between SNPs detected in the acrocentric PVG (**Supplementary Figure 38**) (Paten et al. 2018). For each acrocentric, we plot the r^2^ allele correlation versus distance for three sets of pairs of variants separated by up to 4 kbp: variants on the q-arm, on the p-arm, and within the PHRs. The overall trend of LD decay on the q-arms is comparable to trends seen in other data sets that evaluate LD in humans (Beichman, Phung, and Lohmueller 2017; Bosch et al. 2009). On chromosome 13 the decay of LD in PHRs and p-arm is comparable and faster than the q-arm, while on chromosome 14, 15, and 22 LD decay in PHR is even faster on PHRs compared to the p-arm. The same trend does not apply to chromosome 21 (**Figure 5**).

**Figure 5.**
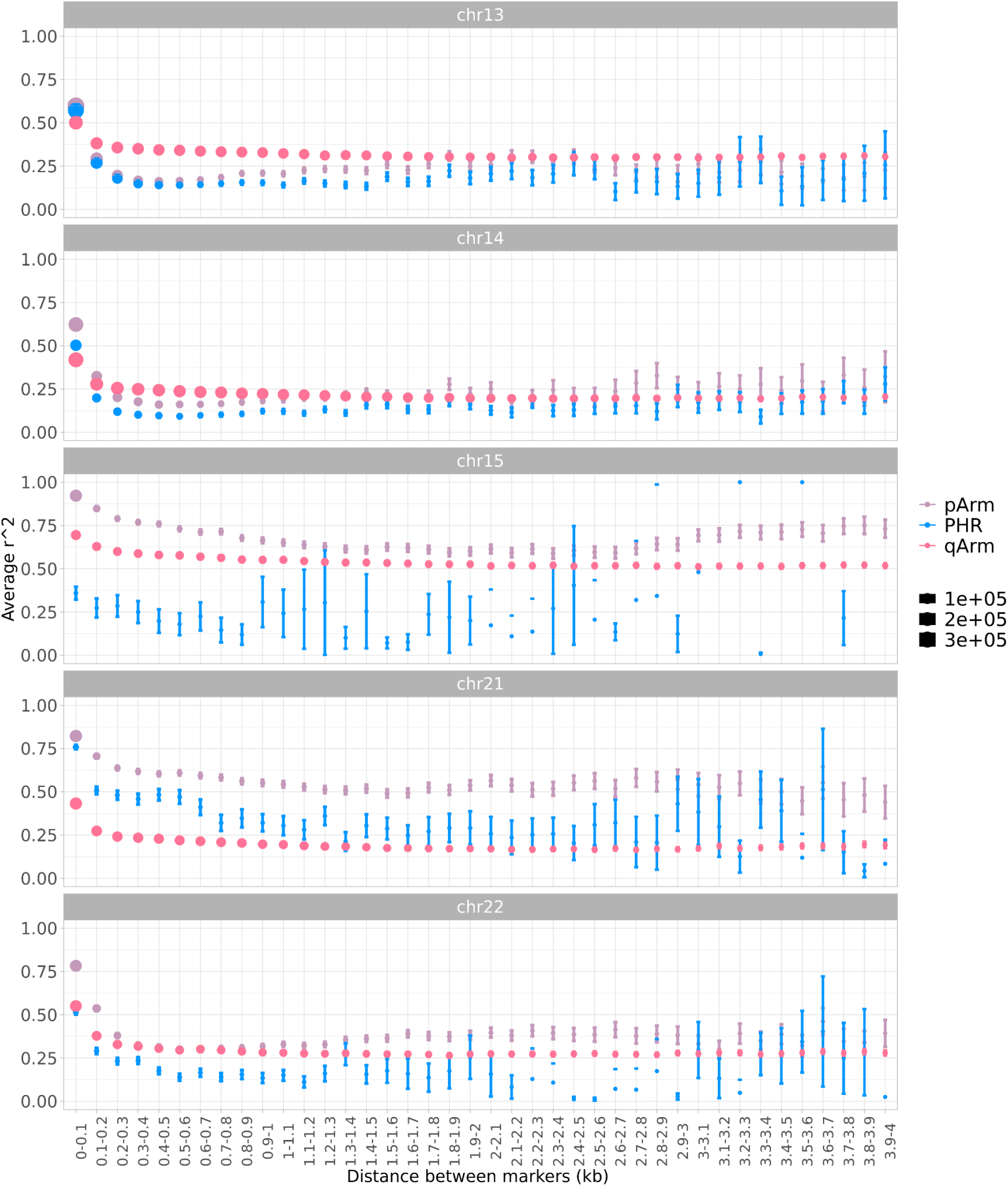
Linkage disequilibrium decay with distance between markers per acrocentric chromosome. Each LD decay plot shows the p-arm (purple), q-arm (pink), and PHR (blue) mean r^2^ and 95% confidence intervals in bins of the given inter-marker distance. Dot size is proportional to the number of pairwise comparisons within a bin. LD decay is faster in PHRs for chromosomes 13, 14, and 22. No significant LD decay is observed in PHRs for chromosome 15.

The fast decay in LD in the PHRs, compared to q-arms, supports the hypothesis of ongoing recombination exchange. In general on the p-arms we find a higher level of LD than on the q-arms, perhaps due to lower recombination in heterochromatic regions (Roberts 1965). However, for the majority of acrocentric chromosomes, we observe the opposite in the PHRs. This effect is clearest within the chromosomes that other analyses suggest to share the most homologous sequence: 13, 14, 21, and 22, while 15—which appears to be an outlier (Huttley et al. 1999) (as also observed in **Figures 1 and 2**)—contains shorter PHRs and we have less confidence in the LD decay trends (error bars on **Figure 5**, chromosome 15 PHRs). This pattern is consistent with increased recombination rate in the PHRs (N. Li and Stephens 2003; Huttley et al. 1999).

## Discussion

We develop multiple lines of evidence indicating active recombination between heterologous human acrocentric chromosomes. First, we find that a symmetric comparison of the sequences of a draft human pangenome contains multi-chromosome communities corresponding to both the sex chromosomes and acrocentric chromosomes (**Figure 1**). An acrocentric pangenome graph reveals base-level homologies that outline patterns of exchange between the heterologous chromosomes (**Figure 2**). The graph highlights regions featuring a diverse patchwork of best-match patterns involving non-homologous T2T-CHM13 chromosomes (**Figure 3**), and we cross-validate these findings with a T2T diploid assembly of a target sample. We develop an entropy metric sensitive to recombination between heterologs such as that seen between chromosomes X and Y, and apply it to delineate pseudo-homologous regions (PHRs) where heterologous SAACs recombine (**Figure 4 and Supplementary File 3**). Finally, we show that on chromosomes 13, 14, 21, and 22, the resulting 18 Mbp of sequence in their PHRs presents a higher rate of linkage disequilibrium decay than seen in sequences from non-PHR regions of the same chromosomes (**Figure 5**). These lines of evidence all suggest that heterologous SAACs recombine.

BACs used in a prior cytogenetic study of Robertsonian chromosomes map to the PHRs of chromosomes 14 and 21, with the recurrent breakpoint region found in a highly homologous region on 13p, 14p, and 21p centered on the PRDM9 motif-enriched SST1 array. This leads us to propose that PHRs are maintained by recombination between heterologous chromosomes, which occasionally results in a Robertsonian translocation (**Figure 6**). We posit that these homologous regions (**Figure 6A**) might share biological function as sequences proximal to the nucleolar organizing regions. Their proximity (**Figure 6B**) can facilitate inter-chromosomal recombination (**Figure 6C**)—which may be of both crossover or non-crossover type, and may occur during meiosis or mitosis. Due to an inversion of this region on 14p relative to 13p and 21p, crossover type recombination between pairs 13+14 and 14+21 leads to Robertsonian translocations (**Figure 6D**), which our study suggests are a pathological outcome of otherwise benign recombination between heterologous chromosomes.

**Figure 6.**
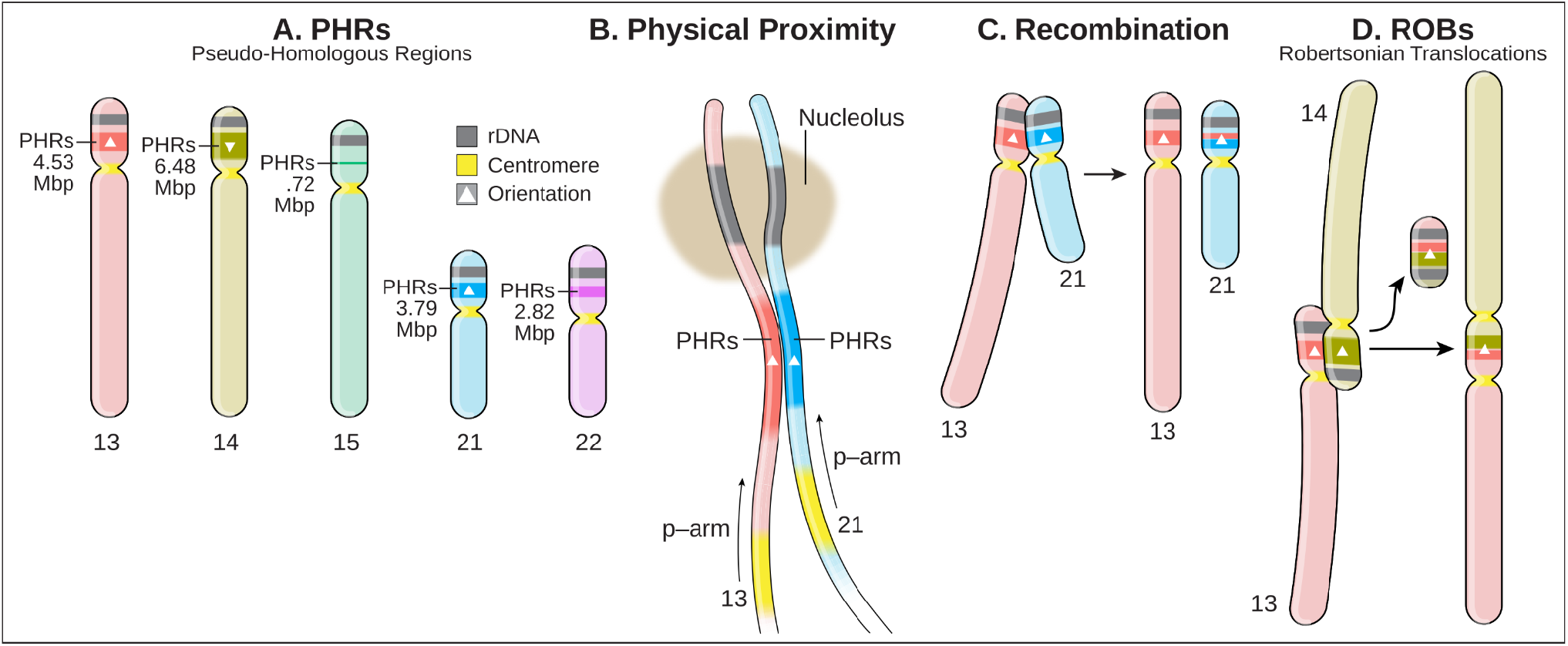
The pseudo-homologous regions of human acrocentric chromosomes. **(A)** PHRs are found on the rDNA-proximal regions of the short arms of acrocentric chromosomes 13, 14, 15, 21, and 22. **(B)** PHRs physically co-locate due to their proximity to the nucleolar organizing regions and rDNA, encouraging sequence exchange. **(C)** Patterns of sequence similarity observed in the PHRs indicate ongoing recombination exchange between heterologous chromosomes, in particular 13, 14, and 21, which may be mediated by both non-crossover recombination or crossover of the telomeric ends of heterologous chromosomes. **(D)** The PHR surrounding the SST1 arrays on 13, 14, and 21 is nearly identical on all three chromosomes, but typically inverted on 14 relative to 13 and 21 (triangles). Due to the inversion, crossover type recombination between PHRs in 14 and 13 or 21 will produce a Robertsonian translocation.

The HPRC pangenome provides base-level resolution of homology patterns across many SAAC haplotypes, letting us examine in detail the regions where ROBs occur. We observe that the GC-rich SST1 array lies at the center of a segmentally duplicated region on 13p, 14p, and 21p which shows a clear pattern of haplotype mixing between these chromosomes. This may also implicate this array as a nucleation point for recombination, as suggested by the observation (N=1/220) of heterologous 14p-21p synapse formation in pachytene oocytes (Cheng and Naluai-Cecchini 2004). We speculate these segmentally duplicated regions are where common Robertsonian translocations occur, a hypothesis supported by our reanalysis of prior cytogenetic mapping of the common ROB breakpoint for 14 and 22 (**Figure 4D**).

Although we find that assembly methods have difficulty in the SAACs (Liao et al. 2022), our validation based on ONT and HiFi data integration shows that the patterns that we observe in HiFi-only assemblies are consistent with ONT reads from the same sample. HG002-Verkko recapitulates key T2T-CHM13-relative untangling patterns also seen in HPRCy1 HiFi assemblies, such as the SST1-linked PHR at 13p11.2 and 21p11.2, rDNA-proximal mixing of all SAACs, and mixing of 22q11.21 and 14q11.2. Our analyses rest on the extensive assembly validation carried out by the HPRC, and observations used to establish signals for recombination are based only on assembly regions deemed to be reliable by mapping analyses (Liao et al. 2022). Our study confirms prior hypotheses based on decades of diverse inquiry, which provides additional assurance that patterns observed bioinformatically are biologically grounded. This body of evidence suggests that our definition of the PHRs is likely to evolve with improved resolution of rDNA arrays and distal regions of the SAACs, which remain among the most challenging regions of the human genome to assemble and lie beyond the scope of our presented work.

Our study is fundamentally population-based. We cannot directly observe recombination of SAACs in this context, leaving open questions about recombination mechanisms that may be difficult to resolve with sequence information alone (Ahuja et al. 2021). However, recombination, like mutation, is a rare event, making it easier to measure its distribution over chromosomes in a population genetic context as we have done here. This addresses key issues with many previous studies of recombination in the SAACs, which often feature small numbers of individuals (Guissani et al. 1996; Cheng and Naluai-Cecchini 2004) selected due to medically-relevant genomic states like trisomy and Robertsonian translocation (Bandyopadhyay et al. 2003). Our resolution of the acrocentric PHRs confirms reported homologies between the SAACs (van Sluis et al. 2019; Nurk et al. 2022) providing a reference for their structure that will be useful to future genomic and cytogenetic studies. In principle, recombination in the PHRs may be of either crossover or non-crossover type. Our data support both, but outside of recurrent Robertsonian translocations and our expectation that non-crossover recombination is significantly more common (~10:1) than crossover (Gay, Myers, and McVean 2007; Cole, Keeney, and Jasin 2010), we lack distinguishing evidence for either. To estimate relative rates of each type of event, we can use linkage disequilibrium patterns (Gay, Myers, and McVean 2007) to study the PHRs in large genomic cohorts (Taliun et al. 2021; Chen et al. 2022), which will require realigning cohort short read data to T2T-CHM13 or the HPRC pangenome. Future improvements to assembly of the SAACs and the planned increase in the number of individuals included in the HPRC should allow for confident estimates of the relative rates of recombination types.

The co-location of ribosomal DNA repeats from different acrocentric chromosomes in a nucleolus provides physical proximity that can facilitate recombination events, not only between rDNA repeats, but between the adjacent PHRs. Our analyses suggest that the rate of recombination between heterologous pairs of acrocentrics varies, which leads to characteristic patterns in the homology spaces that we have explored. Human cells generally have fewer nucleoli (1-5) than acrocentric chromosomes (10) (Soledad Berríos et al. 2004). One possibility is that groups of acrocentrics between which we observe stronger homology and recombination—such as 13, 14, and 21—may be more likely to co-localize to the same nucleolus, as observed in pachytene spermatocytes (Holm and Rasmussen 1977; S. Berríos and Fernández-Donoso 1990). Proximity, homology, recombination initiation sites, and sequence orientation are likely significant factors in the high rate of Robertsonian translocations between these chromosomes. The HPRC draft human pangenome has allowed us to approach genome evolution from a chromosome scale. By stepping away from a reference-centric model and directly comparing whole chromosome assemblies of the acrocentrics, we obtain sequence-resolved responses to long-standing questions first posed in early cytogenetic studies of human genomes.

## Materials and Methods

### Genome assemblies

We analyzed the 47 T2T phased diploid de novo assemblies (94 haplotypes in total) produced by the HPRC and described in (Liao et al. 2022). We included both T2T-CHM13 version 2 (Nurk et al. 2022) and GRCh38.

### Chromosome communities overview

#### The homology graph

We first used all-to-all mapping to build a reference-free model of homology relationships in the HPRCy1 pangenome. This models the full HPRCy1 as a mapping graph in which nodes are contigs and edges represent mappings between them. To build the HPRCy1 mapping graph, we generated homology mappings based on chains of 50 kbp seeds of 95% average nucleotide identity—which we expect to support homologous recombination (Peng et al. 2015)—allowing up to (N-1)=93 alternative mappings over any part of each contig. We first applied this process to map all 38,325 HPRCy1 contigs against all others, obtaining mappings for 38,036 of them covering the 99.9% of the total assembly sequence. This indicated that 38,036 / 38,325 or 99.2% of the HPRCy1 assembly contigs are homologous to at least one other contig. Complex tangles in the assembly graphs used to build the HPRCy1 generate short contigs and tend to result in higher rates of error (Liao et al. 2022). Thus, to simplify later analysis and focus on well-resolved regions of the assemblies, we narrowed our focus to consider only mappings involving the 16,118 contigs at least 1 Mbps long, covering the 98.72% of the total assembly sequence.

We then built a graph where nodes are contigs and edges represent the mappings between them—the “mapping graph”. Edges in this mapping graph have weights equal to the estimated sequence identity multiplied by the length of the mapping. To infer the chromosome represented by each contig, we mapped all contigs against both T2T-CHM13 and GRCh38 references and assigned them a chromosome identity based on this mapping. This mapping graph is very dense, with up to 93 mappings per contig, making it difficult to directly visualize with existing methods. To develop intuition about patterns in this graph, we instead viewed a reduced mapping graph built from the best 3 mappings per contig segment, labeling each contig with its reference-relative assignment (**Figure 1A**). The acrocentric cluster (**Figure 1B**) generally matches our prior expectations of higher similarity between chromosomes 13/21 and 14/22 (Devilee et al. 1986; Jørgensen, Bostock, and Bak 1987; Greig, Warburton, and Willard 1993; Jørgensen et al. 1988).

#### Community detection on the homology mapping graph

To quantify the significance of these patterns, we then applied a community detection algorithm to the full mapping graph. The algorithm assigns each contig to a community such that the total assignment maximizes modularity, which can be understood as the density of (weighted) links inside communities compared to links between communities (Traag, Waltman, and van Eck 2019). This process yielded 31 communities (**Supplementary Table 1**). We hypothesized that each cluster represented one chromosome or chromosome arm. Around half of the chromosomes (N=11) were each represented by a single community. Chromosomes 1, 2, 3, 6, and 18 were each represented in two communities corresponding to their short and long arms, likely due to frequent assembly breaks across their centromeres (**Figure 1C-1D**). Contigs from chromosome X and Y fell in the same community, although the short arm of chromosome X was represented in two communities (**Figure 1D**). The SAACs formed the community with the most distinct chromosomes and most contigs (1706 contigs containing 3.91% of HPRCy1 sequence), composed of contigs belonging to the short arms of all the five acrocentrics plus 21q and 22q (**Figure 1C-1D**). 13q, 14q, and 15q each had their own community. The inclusion of the q-arms of chromosome 21 and 22 in the community composed of p-arms contigs is likely related to their short lengths compared to chromosomes 13, 14, and 15. We obtained similar results when we increased the sensitivity of the mappings (**Supplementary Figure 39**).

In the homology mapping graph of the HPRCy1, only the acrocentric and sex chromosomes form combined communities containing multiple chromosomes. The sex chromosome community reflects the pseudoautosomal regions (PARs) on X and Y (Helena Mangs and Morris 2007), which are telomeric regions where these otherwise non-homologous chromosomes recombine as if they were homologs. We hypothesized that the acrocentric community might also reflect ongoing pseudo-homologous recombination

### Community detection workflow

We performed pairwise mapping for all contigs from the 47 T2T phased diploid de novo assemblies with the WFMASH sequence aligner (Guarracino et al. 2021) (commit ad8aeba). We set the following parameters:

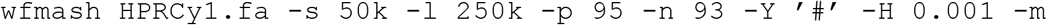

We used segment seed length of 50kbps (`-s`), requiring homologous regions at least ~250 kbp long (`-l`) and estimated nucleotide identity of at least ~95% (`-p`). Having 94 haplotypes in total, we kept up to 93 mappings for each contig (`-n`). Moreover, we skipped mappings when the query and target had the same prefix before the ‘#’ character (`-Y`), that is when involving the same haplotype. To properly map through repetitive regions, only the 0.001% of the most frequent kmers were ignored (`-H`). We skipped the base-level alignment (`-m`). We also generated pairwise mapping with the same parameters, but using a segment seed length of 10kbps and requiring homologous regions at least ~50kbp long.

From the resulting mappings, we excluded those involving contigs shorter than 1 Mbps to reduce the possibility of spurious matches. We then used the `paf2net.py` Python script (delivered in the PGGB repository) to build a graph representation of the result (a mapping graph), with nodes and edges representing contigs and mappings between them, respectively.

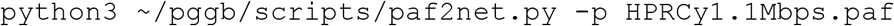

The script produces a file representing the edges, a file representing the edge weights, and a file to map graph nodes to sequence names. The weight of an edge is given by the product of the length and the nucleotide identity of the corresponding mapping (higher weights were associated with longer mappings at higher identities). Finally, we used the `net2communityes.py` Python script (delivered in the PGGB repository) to apply the Leiden algorithm (Traag, Waltman, and van Eck 2019), implemented in the igraph tools (“Igraph Citation Info” n.d.), to detect the underlying communities in the mapping graph.

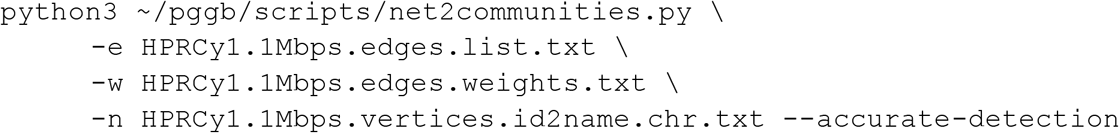

To identify which chromosomes were represented in each community, we partitioned all contigs by mapping them against both T2T-CHM13v1.1 and GRCh38 human reference genomes with WFMASH, this time requiring homologous regions at least 150 kbp long and nucleotide identity of at least 90%.

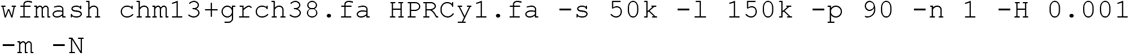

We disabled the contig splitting (`-N`) during mapping to obtain homologous regions covering the whole contigs. For the unmapped contigs, we repeated the mapping with the same parameters, but allowing the contig splitting (without specifying `-N`). We labeled contigs with ‘p’ or ‘q’ depending on whether they cover the short arm or the long arm of the chromosome they belonged to. Contigs fully spanning the centromeres were labeled with ‘pq’. We used such labels to identify the chromosome composition of the communities detected in the mapping graph obtained without reference sequences, and to annotate the nodes in the mapping graph.

To obtain a clean visualization of the homology relationships between the HPRC assemblies, we generated a simpler mapping graph by using the same parameters used for the main graph, but keeping up to 3 mappings for each contig and adding the T2T-CHM13 reference genome version 2, which includes also the complete HG002 chromosome Y (https://www.ncbi.nlm.nih.gov/assembly/GCF_009914755.1):

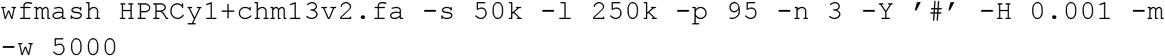

We set window size for sketching equal to 5000 (`-w`) to reduce the runtime by sampling fewer kmers. We used the `paf2net.py` Python script to build the mapping graph and then used Gephi (Bastian, Heymann, and Jacomy 2009) (version 0.9.4) to visualize it. We computed the mapping graph layout by running ‘Random Layout’ and then the ‘Yifan Hu’ algorithm.

### Pangenome graph building

For each of the 47 T2T phased diploid de novo assemblies, we mapped all contigs against the T2T-CHM13 human reference genome with the WFMASH sequence aligner (commit ad8aeba). For the HG002 sample, we included two assemblies: the HG002-HPRCy1 phased diploid de novo assembly (built with HiFi reads) and a phased diploid de novo assembly based on both HiFi and ONT reads, built with the Verkko assembler (Rautiainen et al. 2022). We set the following parameters:

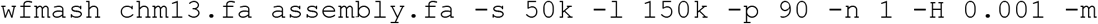

We used segment seed length of 50kbps (`-s`), requiring homologous regions at least ~150 kbp long (`-l`) and estimated nucleotide identity of at least ~90% (`-p`). We kept only 1 mapping (the best one) for each contig (`-n`). To properly map through repetitive regions, only the 0.001% of the most frequent kmers were ignored (`-H`). We skipped the base-level alignment (`-m`). For the HG002-HPRCy1 contigs, we disabled the contig splitting (`-N`).

Then, we identified contigs originating from acrocentric chromosomes and covering both the short and long arms of the chromosome they belonged to. We considered only contigs with mappings at least 1 kbp long on both arms and at least 1 Mbps away from the centromere. We call such contigs “p-q acrocentric contigs”. For HG002-HPRCy1, only contigs longer than or equal to 300 kbps were considered, regardless of covering both arms of the belonging chromosomes.

Finally, we built a pangenome graph with all the p-q acrocentric contigs and both T2T-CHM13 and GRCh38 human reference genomes by applying PGGB (Garrison et al. 2022) (commit a4a6668). We set the following parameters:

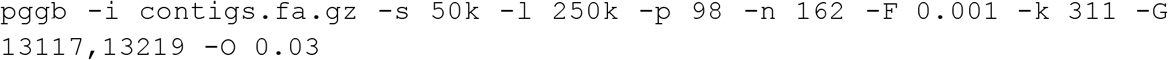

We used segment seed length of 50kbps (`-s`), requiring homologous regions at least ~250 kbp long (`-l`) and estimated nucleotide identity of at least ~98% (`-p`). Having 142 p-q acrocentric contigs in input (132 from HG002-HPRCy1 and 10 from HG002-Verkko) plus 10 acrocentric chromosomes from the T2T-CHM13 and GRCh38 reference genomes plus 49 HG002-HPRCy1 contigs representing other 10 acrocentric haplotypes (5 maternal and 5 paternal), we kept up to 162 mappings (142 +10 + 10) for each contig (`-n`). To properly map through repetitive regions, only the 0.001% of the most frequent kmers were ignored (`-F`). We filtered out alignment matches shorter than 311 bp to remove possible spurious relationships caused by short repeated homologies (`-k`). We set big target sequence lengths and a small sequence padding for two rounds of graph normalization (`-G` and `-O`). To visualize the acrocentric pangenome graph, we built the graph layout with ODGI LAYOUT (Guarracino et al. 2022) (commit e2de6cb) and visualized with GFAESTUS (Fischer and Garrison 2022) (commit 50fe37a).

### Pangenome graph untangling

We untangled the pangenome graph by applying ODGI UNTANGLE (commit e2de6cb). Practically, we projected the graph into an alignment between a set of query (HPRCy1 contigs) and reference (T2T-CHM13) sequences. We set the following parameter:

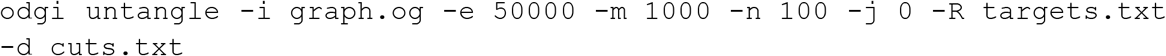

We segmented the graph into regular-sized regions of ~50kbps (`-e`), merging regions shorter than 1kbps (`-m`). We reported up to the 100th best target mapping for each query segment (`-n`), not applying any threshold for the jaccard similarity (`-j`). We used all paths in the graphs as queries and projected them against the 5 acrocentric chromosomes of the T2T-CHM13 genome (`-R`). Moreover, we emit the cut points used to segment the graph (`-d`).

For each query segment, if there were multiple best hits against different targets (i.e., hits with the same, highest jaccard similarity), we put as the first one the hit having as target the chromosome of origin of the query (obtained from the chromosome partitioning of the contigs).

We repeated the graph untangling another 5 times, but constrained the algorithm to use only one T2T-CHM13’s acrocentric chromosome as a target at a time (`-r`) and return the best-matching hit (`-n`).

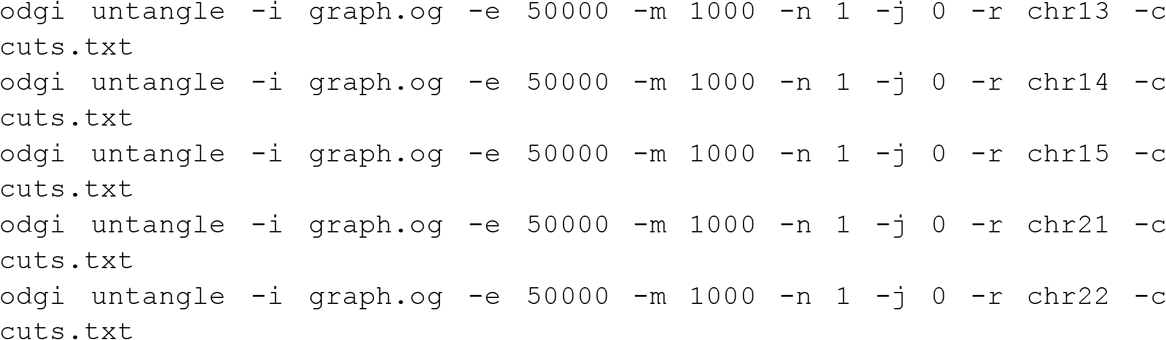

We used the cut points generated when using all T2T-CHM13’s acrocentric chromosomes as targets (`-c`). In this way, all untangling runs (six in total) used the same cut points for the segment boundaries.

Finally, we “grounded” the untangled output generated with all acrocentric chromosomes as targets: in more detail, each untangled query segment was placed against a particular acrocentric chromosome (not only the best-matching one) by using the untangled outputs constrained to a single target. We split the result by acrocentric chromosome and kept only queries untangling both p- and q-arms of the targets. Furthermore, we removed query segments overlapping regions flagged in the assemblies as unreliable (i.e., having coverage issues) by FLAGGER (Liao et al. 2022). FLAGGER is a HiFi read-based pipeline that detects different types of mis-assemblies within a phased diploid assembly by identifying read-mapping coverage inconsistencies across the maternal and paternal haplotypes. To focus on the more similar query-target hits, we used the jaccard metric to estimate the sequence identity by applying the corrected formula reported by (Belbasi et al. 2022), and retained only results at greater than 90% estimated identity. To analyze the orientation status of HPRCy1 contigs in the segmental duplication centered on the SST1 array, we generated a new pangenome graph with ODGI FLIP (commit 0b21b35). In more detail, we first flipped paths around if they tend to be in the reverse complement orientation relative to the pangenome graph. This leads to having an uniform orientation for the HPRCy1 contigs, all in forward orientation with respect to the graph. Then, we untangled the flipped graph in the same way as described above. We displayed the untangling results for each acrocentric chromosome with the R development environment (version 3.6.3), equipped with the following packages: tidyverse (version 1.3.0), RColorBrewer (version 1.1.2), ggplot2 (version 3.3.3) and ggrepel (version 0.9.1).

### Information flow aggregation and diversity entropy

#### Aggregating best-hit untangle results

For each group of HPRCy1-acro contigs anchored to a T2T-CHM13 acrocentric q-arm, we counted the number of contigs having as best-hit each one of the acrocentrics. In particular, for each base position of the T2T-CHM13 acrocentric chromosome of each group, we quantified how many times each of the acrocentrics appeared as best-hit in the pangenome graph untangling. We considered only best-hits with an estimated identity of at least 90%.

#### Regional homology entropy

To quantify the degree of disorder in the untangling result, we calculated the diversity entropy across the different acrocentrics that were present as best-hit. In more detail, we projected each HPRCy1 acrocentric p-q contig against the T2T-CHM13 acrocentric to which it is anchored via the q-arm and associated each reference base position to the corresponding acrocentric best-hit. We considered only best-hits with an estimated identity of at least 90%. Then, we computed the Shannon Diversity Index (SDI) in windows 50 kbp long over the contigs. We used −1 as missing SDI value in the regions where the contigs do not match any targets. For each group of contigs, we aggregated the SDI results by computing their average (ignoring the missing SDI values) for each reference base position. We call this metric “positional homology entropy” and it serves to show regions where contigs can be described as mosaics of different reference chromosomes. However, it cannot distinguish regions where there are different orders of reference chromosome similarity—that might be indicative of recombination exchange—from those where there is regional diversity in each contig’s relationship to T2T-CHM13. The latter case could occur if T2T-CHM13 itself contains rare recombinations between acrocentrics, or where ancient homology might result in “noise” in untangling alignments as contigs pick from two equally-good alternative mappings. To avoid these pitfalls and establish a more stringent graph-space recombination metric, we then extended the untangling diversity metric to operate on multiple mappings.

#### Positional homology entropy and pseudo-homologous regions derivation

To take into account the other hits in addition to the first one, including their order, we generalized the diversity entropy metric to work over orders of the top 5 untantling hits and consider all contigs jointly. For each reference segment, we collected the corresponding best 5 untantling hits for each of the HPRCy1 acrocentric p-q contig; this is possible because the reference segments are stable across all contigs. We considered only best-hits with an estimated identity of at least 90%. To avoid driving the untangle entropy by intra-chromosomal similarity caused by segmental duplications modeled in the structure of the PVG (as seen in loops on 13q, 15q, and 22q, **Figure 2**), we ignored consecutive duplicate target hits—in other words, we took the ordered set of unique reference targets. When multiple contig segments were grounded against the same reference segment, we considered the first contig segment having the best grounding, that is having the highest estimated identity when placed against the current reference segment. Then, we ranked the 5 best-hits by estimated similarity. Finally, for each reference segment, we computed the SDI across all available 5 best-hits orders. We used −1 as missing SDI value in the reference regions without any contig matches. We kept in the output also the information about how many HPRCy1 acrocentric p-q contigs contributed to the entropy computation in each reference segment. This yielded the “positional homology entropy”.

To obtain the pseudo-homologous regions (PHRs), we aggregated the final results by considering regions with positional homology entropy greater than 0 and supported by at least 1 contig, merging with BEDtools (Quinlan and Hall 2010) those that were less than 30 kbps away, and removing merged regions shorter than 30 kbps.

#### Display and annotation of untangle mosaics

We displayed the aggregated results for each acrocentric chromosome. We used genome annotations for the first 25 Mbp of each acrocentric chromosome, using T2T-CHM13v2.0 UCSC trackhub (https://genome.ucsc.edu/cgi-bin/hgTracks?db=hub_3671779_hs1). We made the figures with the scripts available at https://github.com/pangenome/chromosome_communities/tree/main/scripts. To plot the figures, we used R (version 3.6.3), equipped with the following packages: tidyverse (version 1.3.0), RColorBrewer (version 1.1.2), ggplot2 (version 3.3.3) and ggrepel (version 0.9.1). Finally, we used Inkscape (https://inkscape.org/) to compose main-text figures based on the results, and provide supplementary figures directly produced by these methods.

#### HPRCy1 SNP density plots

We displayed biallelic SNP density in the full HPRCy1 draft pangenome built with PGGB (Liao et al. 2022) versus both GRCh38 and T2T-CHM13. To do so, we extracted biallelic SNPs from the released VCF files versus each chromosome, for both references (<u>get_bisnp.sh</u>). Because T2T-CHM13 version 1.1, which was used in HPRCy1, does not have a Y chromosome, we used GRCh38’s, which includes masked PAR1 and PAR2 regions. We displayed biallelic SNP density in bins of 100kbp, using R (version 4.1.1) and tidyverse (version 1.3.1) package (<u>plot_bisnp dens.R</u>).

#### Robertsonian translocation breakpoints

We mapped BAC clones’ from (Jarmuz-Szymczak et al. 2014) against the T2T-CHM13 human reference genome with the WFMASH sequence aligner (commit ad8aeba). We kept only mappings covering acrocentric chromosomes and with an estimated identity of at least 90%. To plot the figures, we used R (version 3.6.3), equipped with the following packages: tidyverse (version 1.3.0), RColorBrewer (version 1.1.2), ggplot2 (version 3.3.3) and ggrepel (version 0.9.1). We colored BAC clones’ mappings according to (Jarmuz-Szymczak et al. 2014).

#### Maximum Likelihood phylogenetic analysis

We conducted the phylogenetic analysis by using the Maximum Likelihood method based on the best-fit substitution model (Kimura 2-parameter +G, parameter = 5.5047) inferred by Jmodeltest2 (Darriba et al. 2012) with 1,000 bootstrap replicates. Bootstrap values higher than 75 are indicated at the base of each node.

#### Recombination hotspots analysis

We obtained the human PRDM9 binding motifs (17 in total) from (Altemose et al. 2017) and used FIMO (Grant, Bailey, and Noble 2011) to scan their occurrences in T2T-CHM13v2.0 human reference genome:

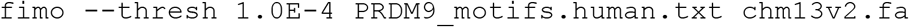

Each motif is associated with a measure of how likely it represents a true binding target for PRDM9. We retained for downstream analyses only motifs for which such a measure is at least 70% (14 of 17). For each motif, we counted the number of occurrences present in windows 20 kbp long across each T2T-CHM13v2.0 chromosome by using BEDtools (Quinlan and Hall 2010).

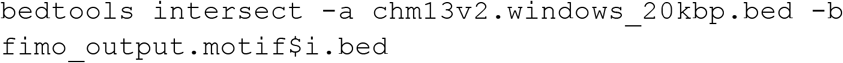

To plot the figures, we used R (version 3.6.3), equipped with the following packages: tidyverse (version 1.3.0), RColorBrewer (version 1.1.2), ggplot2 (version 3.3.3) and ggrepel (version 0.9.1).

### Linkage disequilibrium analysis

We identified variants embedded in the pangenome graph by using VG DECONSTRUCT (Garrison et al. 2018):

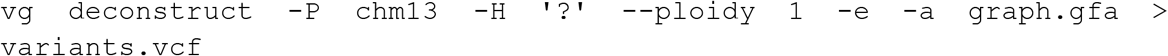

We called variants with respect to the T2T-CHM13 reference genome (`-P`), reporting variants for each HPRCy1 acrocentric p-q contig (`-H` and `--ploidy`). We considered only traversals that correspond to paths (i.e., contigs) in the graph (`-e`) and also reported nested variation (`-a`). From the variant set, we considered only single nucleotide variants. We estimated linkage disequilibrium between pairs of markers within 70kb by using PLINK v1.9 (Purcell et al. 2007) upon specification of haploid sets and retaining all values of r2 > 0 (<u>plot_ld_1.R</u>). Finally, we generated binned LD decay plots with confidence intervals using R (version 3.6.3), focusing on pairs less than 4 kbp apart

### Validating observed patterns of sequence exchange

#### HG002-Verkko assembler

We applied an earlier version of Verkko (beta 1, commit vd3f0b941b5facf5807c303b0c0171202d83b7c74) to build a diploid assembly graph for the HG002 cell line using the HiFi (105x) and ONT (85x) reads as described in (Rautiainen et al. 2022). The resulting assembly graph resolves the proximal junction in single contigs for each haplotype up to multi-mega bases, while the distal junctions (DJ) remain to be resolved. We used homopolymer compressed markers from the parental Illumina reads to assign unitigs to maternal, paternal haplotype or ambiguous when not enough markers supported either haplotype. For estimating the number of times a unitig has to be visited, we aligned HiFi and ONT reads to the assembly graph using GraphAligner with the following parameters: --seeds-mxm-length 30 --seeds-mem-count 10000 -b 15 --multimap-score-fraction 0.99 --precise-clipping 0.85 --min-alignment-score 5000 --clip-ambiguous-ends 100 --overlap-incompatible-cutoff 0.15 --max-trace-count 5 --hpc-collapse-reads --discard-cigar (Rautiainen and Marschall 2020). Four DJ junctions were connected to the rDNA arrays with ambiguous nodes connecting the maternal and paternal nodes, supporting they belong to the same chromosome. Two DJ unitigs, one maternal and paternal, were disconnected from each other but connected to the rDNA arrays, which were assigned to the same chromosome. Using the marker and ONT alignments, we identified paths in the graph and assigned them according to the most supported haplotype. If only ambiguous nodes were present between the haplotype assigned unitigs, with no ONT reads to resolve the path, nodes were randomly assigned to one haplotype to build the contig. After all paths were identified, we produced the consensus using `verkko --assembly <path-to-original-assembly> --paths <path-to-paths>`. The entire procedure to produce parental markers, tagging unitigs according to its haplotype on the assembly graph and finding the path using ONT reads is now available in the latest Verkko (v1) in a more automated way.

#### Validating observed patterns of sequence exchange

To provide cross-validation of the HiFi contig assemblies and our analysis of them, we compared the untangling of two assemblies of the same sample (HG002). One was made with the HPRCy1 pipeline, while the other used the Verkko diploid T2T assembler (Rautiainen et al. 2022). Verkko employs ONT to untangle ambiguous regions in a HiFi-based assembly graph, automating techniques first developed in production of T2T-CHM13. Verkko’s assembly aggregates information from ONT, thus providing a single integrated target for cross-validation of our analysis using an alternative sequencing and assembly approach.

We validated the results of the pangenome untangling by comparing the best-hits of the two HG002 assemblies, the one built with HiFi reads and the other based on both HiFi and ONT reads, built with the Verkko assembler (Rautiainen et al. 2022). For each base position of each T2T-CHM13 acrocentric chromosome, we compared the untangling best-hit of the HG002-HPRCy1 contigs with the best-hit supported by HG002-Verkko contigs. We considered only best-hits with an estimated identity of at least 90%. We defined reference regions as concordant when both HG002 assemblies supported the same T2T-CHM13 acrocentric as best-hit. We treated the two haplotypes (maternal and parental) separately.

We observe a high degree of concordance between the two methods at a level of the chromosome homology mosaicism plots. The best-hit untangling shows similar patterns in the HG002-Verkko assembly as those seen in HG002-HPRCy1 (**Supplementary Figures 18-22**). However, some SAAC haplotypes appear to be poorly-assembled in HG002-HPRCy1. On the q-arms we measured 99.93% concordance between HG002-HPRCy1 and HG002-Verkko untangling results, but only 87.45% concordance on the p-arms (**Supplementary Table 3**). This lower level is consistent with greater difficulty in assembling the SAACs due to their multiple duplicated sequences (including the PHRs), satellite arrays, and the rDNA. We found the discordance was driven by a single chromosome haplotype: while most p-arms achieve around 90% concordance, HG002-HPRCy1 14p-maternal exhibits a high degree of discordance in the assemblies (66.19% concordance) (**Supplementary Figure 19**). Although this N=1 validation focuses on only 10 haplotypes, it incorporates many independent reads provided by deep HiFi (105x) and ONT (85x) data used in HG002-Verkko (Rautiainen et al. 2022). We thus have compared the concordance between structures observed in single molecule reads across the SAACs and the HG002-HPRCy1 assembly that represents the HiFi-only assembly process that produced our pangenome.

However, this analysis should be seen as presenting a lower bound on our process accuracy. We are considering *all* HG002-HPRCy1 contigs that map to the acrocentrics—not only those that would meet our centromere-crossing requirement, and HG002-HPRCy1 is itself more fragmented than the other assemblies which we have selected for the acro-PVG (Liao et al. 2022). The fragmented nature of its contigs (only one from chromosome 22 meets our p-q mapping requirement) may introduce additional disagreements with HG002-Verkko. The overall result indicates that most patterns observed in HiFi-only assemblies are likely to be supported by an automated near-T2T assembly of the same sample.

## Supporting information

a BED file of the acrocentric pseudo-homologous regions (PHRs) on T2T-CHM13

## Acknowledgements

Our work depends on the HPRC draft human pangenome resource established in our companion paper (Liao et al. 2022), and we thank production and assembly groups for their efforts in establishing this resource. This work used the computational resources of the UTHSC Octopus cluster and NIH HPC Biowulf cluster. We acknowledge support in maintaining these systems that was critical to our analyses. We thank Mark Miller of the Stowers Institute for Medical Research for the development of a graphical synopsis of our study (**Figure 6**). We gratefully acknowledge the support and guidance of Prof. Robert Williams of the University of Tennessee Health Science Center and Prof. Nicole Soranzo of the Human Technopole in the design and discussion of our work.

## Funding

This work was supported, in part, by National Institutes of Health/NIDA U01DA047638 (E.G.), National Institutes of Health/NIGMS R01GM123489 (E.G.), NSF PPoSS Award #2118709 (E.G., C.F.), the Tennessee Governor’s Chairs program (C.F., E.G.), National Institutes of Health/NCI R01CA266339 (T.P., L.G.L., J.L.G.), and the Intramural Research Program of the National Human Genome Research Institute, National Institutes of Health (A.R., S.K., A.M.P.). We acknowledge support from Human Technopole (A.G.), Consiglio Nazionale delle Ricerche, Italy (S.B., V.C.), and Stowers Institute for Medical Research (T.P., L.G.L., B.R., J.L.G).

## Supplementary Materials

### Online Materials

Code and links to materials and methods used to perform all the analyses and produce all the figures can be found at the following repository: https://github.com/pangenome/chromosome_communities.

**Combined figures and supplementary material**

We provide links to materials including pangenome graph builds and layouts, and all source files for figures and tables in the manuscript and supplement.

**Main Figures**

https://drive.google.com/drive/folders/1hOK8A-YGhkkJv5IU6hqDH2a23EFc7z8-

**Supplementary Figures**

https://drive.google.com/drive/folders/102iFYLEnqD5FraJN3h771qFw8HSu1vyU

**Files** https://drive.google.com/drive/folders/1FFb-uF60qXg5kBeambBYvQVDUYc3AshW

**Tables** https://drive.google.com/drive/folders/1wOTyxKdv3PjwarFvHLtq-1YJMnXVqGZG

**Notes** https://drive.google.com/drive/folders/1qpmn2L5lAJilMRP4sWAmQDOX5yVHp3_P

### Figure captions

**Supplementary Figure 1.** (**A**) Visualization with Saffire (https://mrvollger.github.io/SafFire/) of the alignment between T2T-CHM13 X and Y reveals that strata 5 and 4 feature low identity (~90%), numerous inversions, and some rearrangements; (**B**) X chromosome ideogram according to (Ross et al. 2005). On the bottom, its evolutionary domains: the X-added region (XAR), the X-conserved region (XCR; dotted region in proximal Xp does not appear to be part of the XCR), the pseudoautosomal region PAR1, and evolutionary strata S5–S1. (**C**) The reduced all-to-all mapping graph of HPRCy1 versus itself, with contigs represented as nodes and mappings as edges, rendered in Gephi (Bastian, Heymann, and Jacomy 2009). In red contigs covering the evolutionary strata 5 and 4 on chromosome X; (**D**) Coloring the reduced homology mapping graph in C with community assignments. Panels **C** and **D** use the same layout as **Figure 1** but focus only on the X and Y region of the visualization.

**Supplementary Figure 2.** An overview of our approach to build a PVG for HPRCy1 contigs that can be anchored to a specific acrocentric q-arm. (**A**) As input, we take the entire HPRCy1 and map it to T2T-CHM13. (**B**) This yields mappings to acrocentric chromosomes, which we filter to select contigs that map across the centromeres (red cytobands) between non-centromeric regions (over-labeled green). We include two HG002 assemblies based on standard HiFi (from HPRCy1) and on both HiFi and ONT data (from Verkko). (**C**) We then apply PGGB to build a PVG from the HPRCy1-acro collection. PGGB first obtains an all-to-all alignment of the input (C.a.), which is converted to a variation graph with SEQWISH (Garrison and Guarracino 2022) (C.b.), then normalized with sorting and multiple sequence alignment steps in SMOOTHXG (C.c-f). (**D**) The resulting PVG expresses genomes as paths, or walks, through a common sequence graph. This model thus contains all input sequences and their relative alignments to all others—in the example we see a CTGG/AAGTA block substitution between genomes 1 and 2.

**Supplementary Figure 3.** Regions of a PVG built from centromere-spanning HPRCy1 contigs plus T2T-CHM13 and GRCh38 references. We apply GFAESTUS to visualize the 2D layout generated by ODGI in the PGGB pipeline. We focus on the segmentally duplicated core centered in the SST1 array (labeled).

**Supplementary Figure 4**. A PVG built from centromere-spanning HPRCy1 contigs, without embedding the T2T-CHM13 and GRCh38 references. We apply ODGI to visualize the 2D layout generated in the PGGB pipeline. This renders sequences and chains of small variants as linear structures, while repeats caused by segmental duplications, inversions, and other structural variants tend to form loops and tangles. The acrocentric q-arms are almost completely separated, while the p-arms unite in a structure adjacent to the rDNA array.

**Supplementary Figure 5.** Regions of a PVG built from centromere-spanning HPRCy1 contigs without the T2T-CHM13 and GRCh38 references. We apply GFAESTUS to visualize the 2D layout generated by ODGI in the PGGB pipeline. We focus on the segmentally duplicated core centered in the SST1 array (labeled).

**Supplementary Figure 6.** Scheme of the graph untangling. We applied ODGI UNTANGLE to obtain a mapping from segments of all PVG paths onto T2T-CHM13. The segmentation cuts the graph into regular-sized regions whose boundaries occur at structural variant breakpoints. For each query subpath through a graph segment, we use a Jaccard metric over the sequence space of the subpaths to find the best-matching reference segment.

**Supplementary Figures 7-11.** Untangling of HPRCy1-acro’s sequences belonging to chromosome 13, 14, 15, 21, 22 versus T2T-CHM13. We display all mappings above 90% estimated pairwise identity, without removing those covering regions classified as unreliable. Above each contig, we display the unreliable regions in black.

**Supplementary Figure 12.** Centromeric Satellite Annotation (CenSat Annotation) track legend.

**Supplementary Figures 13-17. (A)** We focus on the first 25 Mbp of chromosome 13, 14, 15, 21, 22, shown here as a red box over T2T-CHM13 cytobands. Pseudo-homologous regions (**PHRs**), where diverse sets of acrocentric chromosomes recombine, are highlighted relative to T2T-CHM13 genome annotations for repeats, GC percentage, and genes. Above, we indicate regions of interest described in the main text: rDNA, SST1 array, centromere, and q-arm. Below, we show T2T-CHM13-relative homology mosaics for each chromosome 13 matched contig from HPRCy1-acro, with the most-similar reference chromosome at each region shown using the given colors (**Target**). **(B)** Aggregated untangle results in the SAACs. For each acrocentric chromosome, we show the count of its HPRCy1 q-arm-anchored contigs mapping itself and all other acrocentrics (**Contigs**), **(C)** as well as the regional (50kbp) untangle entropy metric (**Regional homology entropy**) computed over the contigs’ T2T-CHM13-relative untanglings. **(D)** By considering the multiple untangling of each HPRCy1-acro contig, we develop a point-wise metric that captures diversity in T2T-CHM13-relative homology patterns (**Positional homology entropy**), leading to our definition of the PHRs. **(E)** The patterns of homology mosaicism suggest ongoing recombination exchange in the SAACs. A scan over T2T-CHM13 reveals that the rDNA units are enriched for PRDM9 binding motifs, and thus may host frequent double stranded breaks during meiosis. In (B-D) a gray background indicates regions with missing data due to the lack of non-T2T-CHM13 contigs. We provide the Centromeric Satellite Annotation (CenSat Annotation) track legend in **Supplementary Figure 12**.

**Supplementary Figures 18-22.** For each base position of T2T-CHM13 chromosome 13, 14, 15, 21, 22, we compared the untangling best-hit of the HG002-HPRCy1 contigs with the best-hit supported by HG002-Verkko’s contigs. We considered only best-hits with an estimated identity of at least 90% that cover regions labeled as reliable. We reported the number of different targets for each reference position. On the bottom, the untangling of HG002’s contigs from HG002-Verkko assembly as those seen in HG002-HPRCy1 assemblies, for chromosome 13, 14, 15, 21, 22 versus T2T-CHM13. Transparency shows the estimated identity of the mappings. We display all mappings above 90% estimated pairwise identity. Checkerboard patterns observed in several regions of the SAACs correspond to contexts that may permit ongoing recombination.

**Supplementary Figures 23.** Multiple untangling of T2T-CHM13, GRCh38, HG002-Verkko haplotypes, and HPRCy1-acro contigs versus T2T-CHM13. Chromosome 13 results are represented, in the chr13:11,301,367-13,440,010 region (censat_13_27 coordinates ± 1Mbp). Transparency shows the different orientations of the mappings. We display all mappings above 90% estimated pairwise identity. To analyze simultaneous hits to all acrocentrics, each grouping shows the first 3 best alternative mappings. The figure shows that chromosome 13’s contigs map in forward orientation on T2T-CHM13 chromosome 13 and 21 (orange and cyan rectangles), while their mappings are inverted on chromosomes 14 (transparent gold rectangles).

**Supplementary Figures 24.** Multiple untangling of T2T-CHM13, GRCh38, HG002-Verkko haplotypes, and HPRCy1-acro contigs versus T2T-CHM13. Chromosome 14 results are represented, in the chr14:5,960,008-7,988,409 region (censat_14_39 coordinates ± 1Mbp). Transparency shows the different orientations of the mappings. We display all mappings above 90% estimated pairwise identity. To analyze simultaneous hits to all acrocentrics, each grouping shows the first 3 best alternative mappings. The figure shows that chromosome 14’s contigs map in forward orientation on T2T-CHM13 chromosome 14 (gold rectangles), while their mappings are inverted on chromosomes 13 and 21 (transparent orange and cyan rectangles), with the sole exception of HG01071#2#JAHBCE010000082.1, where the trend is reversed.

**Supplementary Figures 25.** Multiple untangling of T2T-CHM13, GRCh38, HG002-Verkko haplotypes, and HPRCy1-acro contigs versus T2T-CHM13. Chromosome 21 results are represented, in the chr21:8,375,567-10,453,313 region (censat_21_45 coordinates ± 1Mbp). Transparency shows the different orientations of the mappings. We display all mappings above 90% estimated pairwise identity. To analyze simultaneous hits to all acrocentrics, each grouping shows the first 3 best alternative mappings. The figure shows that chromosome 21’s contigs map in forward orientation on T2T-CHM13 chromosome 21 and 13 (cyan and orange rectangles), while their mappings are inverted on chromosomes 14 (transparent gold rectangles).

**Supplementary Figure 26.** For HPRCy1 contigs from chromosome X and Y, we show the tracks for the pseudoautosomal regions (PARs) and the X-transposed region (XTRs) with respect to T2T-CHM13 (on the top) as well as the untangle entropy metric (Regional homology metric, on the bottom) computed over the contigs’ T2T-CHM13-relative untanglings.

**Supplementary Figures 27-31.** Multiple untangling of HPRCy1-acro’s sequences belonging to chromosome 13, 14, 15, 21, 22 versus T2T-CHM13. Transparency shows the estimated identity of the mappings. We display all mappings above 90% estimated pairwise identity that cover regions labeled as reliable. To allow the display of simultaneous hits to all acrocentrics, each grouping shows the first 5 best alternative mappings. Checkerboard patterns observed in several regions of the SAACs correspond to contexts that may permit ongoing recombination.

**Supplementary Figure 32.** Multi-hit untangling diversity entropy for HPRCy1-acro’s sequences belonging to chromosome 13, 14, 15, 21, 22 versus T2T-CHM13. We considered all mappings above 90% estimated pairwise identity that cover regions labeled as reliable.

**Supplementary Figure 33.** For each T2T-CHM13 acrocentric chromosome, we show the tracks for the pseudo-homologous regions (PHRs), the SST1 array, the rDNA arrays, and for the regions where the bacterial artificial chromosome (BAC) clones from (Jarmuz-Szymczak et al. 2014) map on those chromosomes. Most of the mappings cover the PHRs. The mappings with higher estimated identity (> 99%) are on chr21 and chr14. We colored BAC clones’ mappings according to (Jarmuz-Szymczak et al. 2014).

**Supplementary Figure 34.** Comparison of chr13 and chr17 SST1 consensus sequences by alignment dotplot indicate the unique deletion shared exclusively by the acrocentric chromosomes (dotted red rectangle).

**Supplementary Figures 35.** For HPRCy1 contigs from chromosome X and Y, we show the tracks for the pseudoautosomal regions (PARs) and the X-transposed region (XTRs) with respect to T2T-CHM13 as well as the multi untangle entropy metric (Positional homology entropy, on the bottom) computed over the contigs’ T2T-CHM13-relative untanglings.

**Supplementary Figure 36.** Biallelic SNP density in the full HPRCy1 draft pangenome. Colors indicate the count of SNPs in each bin at given variant nesting levels (Paten et al. 2018), with gray showing the base level and a red to violet spectrum for increasing level.

**Supplementary Figure 37.** For each T2T-CHM13 acrocentric chromosome, we show the number of human PRDM9 binding motif hits present in windows 20 kbps long.

**Supplementary Figure 38.** *Left*) Histograms showing missing genotypes per site for variants called with respect to the T2T-CHM13 acrocentrics. *Middle*) Histograms showing the allele frequency spectrum for variants called with respect to the T2T-CHM13 acrocentrics. MAF = Minor Allele Frequency. *Right-Top*) Segment size distributions of the pseudo-homologous regions with respect to the T2T-CHM13 acrocentrics. *Right-Bottom*) Distributions of the distance between consecutive markers with respect to the T2T-CHM13 acrocentrics.

**Supplementary Figure 39**. Results of community assignment on the mapping graph when using chains of 10 kbp seeds of 95% average nucleotide identity. On the x-axis the chromosome to which contigs belong based on competitive mapping to T2T-CHM13 and GRCh38, while the y-axis indicates the community, which is named by the chromosome partition that contributes the largest number of contigs to it.

### File descriptions

**Supplementary File 1.** Acrocentric pangenome variation graph in GFAv1 file format (https://github.com/GFA-spec/GFA-spec).

**Supplementary File 2.** Graph layout in 2 dimensions. The first column is an incremental number, the second and third columns are the X and Y coordinates and the fourth column identifies the graph connected component.

**Supplementary File 3.** Pseudo-homologous regions identified in the acrocentric chromosomes of T2T-CHM13. The first three columns represent the T2T-CHM13 chromosome, the starting position, and the ending position of each pseudo-homologous region. Column 4 is the average Shannon Diversity Index. Column 5 is the average number of contigs.

**Supplementary File 4.** Pseudo-homologous regions identified in the sex chromosomes of T2T-CHM13. The first three columns represent the T2T-CHM13 chromosome, the starting position, and the ending position of each pseudo-homologous region. Column 4 is the average Shannon Diversity Index. Column 5 is the average number of contigs.

**Supplementary File 5.** PRDM9 binding motif hits in T2T-CHM13. Column 1 and 2 identify the motif. Column 3, 4, 5 indicate the sequence on which the motif has been found, with the start and end coordinates. Column 6 reports the strand of the hit on the sequence. Column 7 is the score of the motif hit. Column 8 and 9 report, respectively, the p-value and the adjusted p-value. Column 10 is the matched sequence.

### Table captions

**Supplementary Table 1.** Results of community assignment on the mapping graph. The ‘community.of’ column reports the community names assigned by the chromosomal partition that contributes the most contigs. Columns ‘chr1’ to ‘chrY’ report the number of contigs assigned to each chromosome based on the competitive mapping to the chromosomes of T2T-CHM13 and GRCh38. The ‘non.partitioned’ column reports the number of contigs not mappable to T2T-CHM13 and GRCh38.

**Supplementary Table 2.** For each HPRCy1-acro contig (‘contig’ column), we report the amount of projected alignments (‘untangled.size’ column), the amount of reliable projected alignments (‘reliable.untangled.size’ column), and the fraction of unreliable alignments that we ignored in our analysis (‘fraction.removed’ column). Regions are labeled as reliable or unreliable by FLAGGER (Liao et al. 2022).

**Supplementary Table 3.** For each T2T-CHM13 chromosome (‘ground.target’ column), we report the number of bases where HG002-HiFi and HG00-Verkko assemblies are concordant and discordant (‘num.bp.ok’ and ‘num.bp.not.ok’ columns), and the percentage of concordant bases (‘percentage.bp.ok’ column). Concordant bases are those where the untangling best-hits from both assemblies is on the same T2T-CHM13 chromosome. Values are reported for both maternal and paternal haplotypes (‘haplotype’ column) and stratified by chromosomal arm or whole region (‘region’ column).

### Note captions

**Supplementary Note 1.** Physical proximity modeling of acrocentric short arms.

Human Pangenome Reference Consortium

Haley J. Abel^1^, Lucinda L Antonacci-Fulton^2^, Mobin Asri^3^, Gunjan Baid^4^, Carl A. Baker^5^, Anastasiya Belyaeva^4^, Konstantinos Billis^6^, Guillaume Bourque^7,8,9^, Silvia Buonaiuto^10^, Andrew Carroll^4^, Mark JP Chaisson^11^, Pi-Chuan Chang^4^, Xian H. Chang^3^, Haoyu Cheng^12,13^, Justin Chu^12^, Sarah Cody^2^, Vincenza Colonna^10,14^, Daniel E. Cook^4^, Omar E. Cornejo^15^, Mark Diekhans^3^, Daniel Doerr^16^, Peter Ebert^16^, Jana Ebler^16^, Evan E. Eichler^5,17^, Jordan M. Eizenga^3^, Susan Fairley^6^, Olivier Fedrigo^18^, Adam L. Felsenfeld^19^, Xiaowen Feng^12,13^, Christian Fischer^14^, Paul Flicek^6^, Giulio Formenti^18^, Adam Frankish^6^, Robert S. Fulton^2^, Yan Gao^20^, Shilpa Garg^21^, Erik Garrison^14^, Carlos Garcia Giron^6^, Richard E. Green^22,23^, Cristian Groza^24^, Andrea Guarracino^25^, Leanne Haggerty^6^, Ira Hall^26,27^, William T Harvey^5^, Marina Haukness^3^, David Haussler^3,17^, Simon Heumos^28,29^, Glenn Hickey^3^, Kendra Hoekzema^5^, Thibaut Hourlier^6^, Kerstin Howe^30^, Miten Jain^31^, Erich D. Jarvis^32,17^, Hanlee P. Ji^33^, Alexey Kolesnikov^4^, Jan O. Korbel^34^, Sergey Koren^41^, Jennifer Kordosky^5^, HoJoon Lee^33^, Alexandra P. Lewis^5^, Heng Li^12,13^, Wen-Wei Liao^2,35,26^, Shuangjia Lu^26^, Tsung-Yu Lu^11^, Julian K. Lucas^3^, Hugo Magalhães^16^, Santiago Marco-Sola^36,37^, Pierre Marijon^16^, Charles Markello^3^, Tobias Marschall^16^, Fergal J. Martin^6^, Ann McCartney^41^, Jennifer McDaniel^38^, Karen H. Miga^3^, Matthew W. Mitchell^39^, Jean Monlong^3^, Jacquelyn Mountcastle^18^, Katherine M. Munson^5^, Moses Njagi Mwaniki^40^, Maria Nattestad^4^, Adam M. Novak^3^, Sergey Nurk^41^, Hugh E. Olsen^3^, Nathan D. Olson^38^, Benedict Paten^3^, Trevor Pesout^3^, Adam M. Phillippy^41^, David Porubsky^5^, Alice B. Popejoy^52^, Pjotr Prins^14^, Daniela Puiu^42^, Mikko Rautiainen^41^, Allison A Regier^2^, Arang Rhie^41^, Samuel Sacco^43^, Ashley D. Sanders^44^, Valerie A. Schneider^45^, Baergen I. Schultz^19^, Kishwar Shafin^4^, Jonas A. Sibbesen^46^, Jouni Sirén^3^, Michael W. Smith^19^, Heidi J. Sofia^19^, Ahmad N. Abou Tayoun^47,48^, Françoise Thibaud-Nissen^45^, Chad Tomlinson^2^, Francesca Floriana Tricomi^6^, Flavia Villani^14^, Mitchell R. Vollger^5,49^, Justin Wagner^38^, Brian Walenz^41^, Ting Wang^50^, Jonathan M. D. Wood^30^, Aleksey V. Zimin^42,51^, Justin M. Zook^38^

1 Division of Oncology, Department of Internal Medicine, Washington University School of Medicine, St. Louis, MO 63110, USA

2 McDonnell Genome Institute, Washington University School of Medicine, St. Louis, MO 63108, USA

3 UC Santa Cruz Genomics Institute, University of California, Santa Cruz, 1156 High St, Santa Cruz, CA, USA

4 Google LLC, 1600 Amphitheater Pkwy, Mountain View, CA 94043, USA

5 Department of Genome Sciences, University of Washington School of Medicine, Seattle, WA 98195, USA

6 European Molecular Biology Laboratory, European Bioinformatics Institute, Wellcome Genome Campus, Cambridge, CB10 1SD, UK

7 Department of Human Genetics, McGill University, Montreal, Québec H3A 0C7, Canada

8 Canadian Center for Computational Genomics, McGill University, Montreal, Québec H3A 0G1, Canada

9 Institute for the Advanced Study of Human Biology (WPI-ASHBi), Kyoto University, Kyoto 606-8501, Japan

10 Institute of Genetics and Biophysics, National Research Council, Naples 80111, Italy

11 University of Southern California, Quantitative and Computational Biology, 3551 Trousdale, Pkwy, Los Angeles, CA, USA

12 Department of Data Sciences, Dana-Farber Cancer Institute, Boston, MA 02215, USA

13 Department of Biomedical Informatics, Harvard Medical School, Boston, MA 02215, USA

14 Department of Genetics, Genomics and Informatics, University of Tennessee Health Science Center, Memphis, TN 38163, USA

15 School of Biological Sciences, Washington State University, Pullman WA 99163, USA

16 Institute for Medical Biometry and Bioinformatics, Medical Faculty, Heinrich Heine University Düsseldorf, Düsseldorf, Germany

17 Howard Hughes Medical Institute, Chevy Chase, MD 20815, USA

18 The Vertebrate Genome Laboratory, The Rockefeller University, New York, NY 10065, USA

19 National Institutes of Health (NIH)–National Human Genome Research Institute, Bethesda, MD, USA

20 Center for Computational and Genomic Medicine, The Children’s Hospital of Philadelphia, Philadelphia, PA 19104, USA.

21 Department of Biology, University of Copenhagen, Denmark

22 Department of Biomolecular Engineering, University of California, Santa Cruz, 1156 High St., Santa Cruz, CA 95064, USA

23 Dovetail Genomics, Scotts Valley, CA 95066, USA

24 Quantitative Life Sciences, McGill University, Montreal, Québec H3A 0C7, Canada

25 Genomics Research Centre, Human Technopole, Milan 20157, Italy

26 Department of Genetics, Yale University School of Medicine, New Haven, CT 06510, USA

27 Center for Genomic Health, Yale University School of Medicine, New Haven, CT 06510, USA

28 Quantitative Biology Center (QBiC), University of Tübingen, Tübingen 72076, Germany

29 Biomedical Data Science, Department of Computer Science, University of Tübingen, Tübingen 72076, Germany

30 Tree of Life, Wellcome Sanger Institute, Hinxton, Cambridge, CB10 1SA, UK

31 Northeastern University, Boston, MA 02115, USA

32 The Rockefeller University, New York, NY 10065, USA

33 Division of Oncology, Department of Medicine, Stanford University School of Medicine, Stanford, CA, 94305, USA

34 European Molecular Biology Laboratory, Genome Biology Unit, Meyerhofstr. 1, 69117 Heidelberg, Germany

35 Department of Medicine, Washington University School of Medicine, St. Louis, MO 63110, USA

36 Computer Sciences Department, Barcelona Supercomputing Center, Barcelona, Spain

37 Departament d’Arquitectura de Computadors i Sistemes Operatius, Universitat Autònoma de Barcelona, Barcelona, Spain

38 Material Measurement Laboratory, National Institute of Standards and Technology, Gaithersburg, MD 20877, USA

39 Coriell Institute for Medical Research, Camden, NJ 08103, USA

40 Department of Computer Science, University of Pisa, Pisa 56127, Italy

41 Genome Informatics Section, Computational and Statistical Genomics Branch, National Human

Genome Research Institute, National Institutes of Health, Bethesda, MD 20892, USA

42 Department of Biomedical Engineering, Johns Hopkins University, Baltimore 21218, MD, USA

43 Department of Ecology & Evolutionary Biology, University of California, Santa Cruz, 1156 High St, Santa Cruz, CA, USA

44 Berlin Institute for Medical Systems Biology, Max Delbrück Center for Molecular Medicine in the Helmholtz Association, Berlin, Germany

45 National Center for Biotechnology Information, National Library of Medicine, National Institutes of Health, Bethesda, MD 20894, USA

46 Center for Health Data Science, University of Copenhagen, Denmark

47 Al Jalila Genomics Center of Excellence, Al Jalila Children’s Specialty Hospital, Dubai, UAE

48 Center for Genomic Discovery, Mohammed Bin Rashid University of Medicine and Health Sciences, Dubai, UAE

49 Division of Medical Genetics, University of Washington School of Medicine, Seattle, WA 98195, USA

50 Department of Genetics, Washington University School of Medicine, St. Louis, MO 63110, USA

51 Center for Computational Biology, Johns Hopkins University, Baltimore, MD 21218, USA

52 Department of Public Health Sciences, University of California, Davis, One Shields Avenue, Medical Sciences 1C, Davis, CA 95616

## Notes

https://doi.org/10.5281/zenodo.6993789

https://github.com/pangenome/chromosome_communities

### Competing Interest Statement

The authors have declared no competing interest.

### Summary of Updates

We have extended the work to relate our observation of recombination between heterologous acrocentric chromosomes to the sites of recurrent Robertsonian translocations. We extend our study of this locus with a whole genome phylogenetic survey of the SST1 array that lies at its center. Our specific definition of the pseudo-homologous regions has improved due to use of the sex chromosomes as a positive control, leading to an increase in the power of our linkage disequilibrium analysis. We have extensively rewritten the manuscript to improve its readability by a general scientific audience. We added a new figure that summarizes our results and have generally improved our figure presentation. We include extended information on zenodo and supplementary figures and the PHR BEDs alongside the manuscript here.

## References

Ahuja, Jasvinder S., Catherine S. Harvey, David L. Wheeler, and Michael Lichten. 2021. “Repeated Strand Invasion and Extensive Branch Migration Are Hallmarks of Meiotic Recombination.” Molecular Cell 81 (20): 4258–70.e4.

Altemose, Nicolas, Nudrat Noor, Emmanuelle Bitoun, Afidalina Tumian, Michael Imbeault, J. Ross Chapman, A. Radu Aricescu, and Simon R. Myers. 2017. “A Map of Human PRDM9 Binding Provides Evidence for Novel Behaviors of PRDM9 and Other Zinc-Finger Proteins in Meiosis.” eLife 6 (October). https://doi.org/10.7554/eLife.28383.

Arenas, Miguel. 2013. “The Importance and Application of the Ancestral Recombination Graph.” Frontiers in Genetics 4 (October): 206.

Armstrong, Joel, Glenn Hickey, Mark Diekhans, Ian T. Fiddes, Adam M. Novak, Alden Deran, Qi Fang, et al. 2020. “Progressive Cactus Is a Multiple-Genome Aligner for the Thousand-Genome Era.” Nature 587 (7833): 246–51.

Bandyopadhyay, Ruma, Christopher McCaskill, Cami Knox-Du Bois, Yaolin Zhou, Sue Ann Berend, Emilia Bijlsma, and Lisa G. Shaffer. 2003. “Mosaicism in a Patient with Down Syndrome Reveals Post-Fertilization Formation of a Robertsonian Translocation and Isochromosome.” American Journal of Medical Genetics. Part A 116A (2): 159–63.

Bastian, Mathieu, Sebastien Heymann, and Mathieu Jacomy. 2009. “Gephi: An Open Source *Software for Exploring and Manipulating Networks*.” Proceedings of the International AAAI Conference on Web and Social Media 3 (1): 361–62.

Beichman, Annabel C., Tanya N. Phung, and Kirk E. Lohmueller. 2017. “Comparison of Single Genome and Allele Frequency Data Reveals Discordant Demographic Histories.” G3 7 (11): 3605–20.

Belbasi, Mahdi, Antonio Blanca, Robert S. Harris, David Koslicki, and Paul Medvedev. 2022. “The Minimizer Jaccard Estimator Is Biased and Inconsistent.” Bioinformatics 38 (Suppl 1): i169–76.

Berríos, S., and R. Fernández-Donoso. 1990. “Nuclear Architecture of Human Pachytene Spermatocytes: Quantitative Analysis of Associations between Nucleolar and XY Bivalents.” Human Genetics 86 (2): 103–16.

Berríos, Soledad, Raúl Fernández-Donoso, Juana Pincheira, Jesús Page, Marcia Manterola, and M. Cristina Cerda. 2004. “Number and Nuclear Localisation of Nucleoli in Mammalian Spermatocytes.” Genetica 121 (3): 219–28.

Bosch, Elena, Hafid Laayouni, Carlos Morcillo-Suarez, Ferran Casals, Andrés Moreno-Estrada, Anna Ferrer-Admetlla, Michelle Gardner, et al. 2009. “Decay of Linkage Disequilibrium within Genes across HGDP-CEPH Human Samples: Most Population Isolates Do Not Show Increased LD.” BMC Genomics 10 (July): 338.

Cheng, Edith Y., and Theresa Naluai-Cecchini. 2004. “FISHing for Acrocentric Associations *between Chromosomes 14 and 21 in Human Oogenesis*.” American Journal of Obstetrics and Gynecology 190 (6): 1781–85; discussion 1785–87.

Chen, Siwei, Laurent C. Francioli, Julia K. Goodrich, Ryan L. Collins, Masahiro Kanai, Qingbo Wang, Jessica Alföldi, et al. 2022. “A Genome-Wide Mutational Constraint Map Quantified from Variation in 76,156 Human Genomes.” bioRxiv. https://doi.org/10.1101/2022.03.20.485034.

Choo, K. H., B. Vissel, R. Brown, R. G. Filby, and E. Earle. 1988. “Homologous Alpha Satellite Sequences on Human Acrocentric Chromosomes with Selectivity for Chromosomes 13, 14 and 21: Implications for Recombination between Nonhomologues and Robertsonian Translocations.” Nucleic Acids Research 16 (4): 1273–84.

Cole, Francesca, Scott Keeney, and Maria Jasin. 2010. “Comprehensive, Fine-Scale Dissection of Homologous Recombination Outcomes at a Hot Spot in Mouse Meiosis.” Molecular Cell 39 (5): 700–710.

Cotter, Daniel J., Sarah M. Brotman, and Melissa A. Wilson Sayres. 2016. “Genetic Diversity on the Human X Chromosome Does Not Support a Strict Pseudoautosomal Boundary.” Genetics 203 (1): 485–92.

Darriba, Diego, Guillermo L. Taboada, Ramón Doallo, and David Posada. 2012. “jModelTest 2: More Models, New Heuristics and Parallel Computing.” Nature Methods 9 (8): 772.

Devilee, P., T. Cremer, P. Slagboom, E. Bakker, H. P. Scholl, H. D. Hager, A. F. Stevenson, C. J. Cornelisse, and P. L. Pearson. 1986. “Two Subsets of Human Alphoid Repetitive DNA Show Distinct Preferential Localization in the Pericentric Regions of Chromosomes 13, 18, and 21.” Cytogenetics and Cell Genetics 41 (4): 193–201.

Eizenga, Jordan M., Adam M. Novak, Jonas A. Sibbesen, Simon Heumos, Ali Ghaffaari, Glenn Hickey, Xian Chang, et al. 2020. “Pangenome Graphs.” Annual Review of Genomics and Human Genetics 21 (August): 139–62.

Epstein, N. D., S. Karlsson, S. O’Brien, W. Modi, A. Moulton, and A. W. Nienhuis. 1987. “A New *Moderately Repetitive DNA Sequence Family of Novel Organization*.” Nucleic Acids Research 15 (5): 2327–41.

Fischer, Christian, and Erik Garrison. 2022. Chfi/gfaestus: A Pangenome Graph Browser. https://doi.org/10.5281/zenodo.6954036.

Floutsakou, Ioanna, Saumya Agrawal, Thong T. Nguyen, Cathal Seoighe, Austen R. D. Ganley, and Brian McStay. 2013. “The Shared Genomic Architecture of Human Nucleolar Organizer Regions.” Genome Research 23 (12): 2003–12.

Garrison, Erik, and Andrea Guarracino. 2022. “Unbiased Pangenome Graphs.” Bioinformatics, November. https://doi.org/10.1093/bioinformatics/btac743.

Garrison, Erik, Andrea Guarracino, Simon Heumos, Joerg Hagmann, and Peter Sudmant. 2022. Pggb: The PanGenome Graph Builder. https://doi.org/10.5281/zenodo.6949381.

Garrison, Erik, Jouni Sirén, Adam M. Novak, Glenn Hickey, Jordan M. Eizenga, Eric T. Dawson, William Jones, et al. 2018. “Variation Graph Toolkit Improves Read Mapping by Representing Genetic Variation in the Reference.” Nature Biotechnology 36 (9): 875–79.

Gay, J., S. Myers, and G. McVean. 2007. “Estimating Meiotic Gene Conversion Rates from Population Genetic Data.” Genetics 177 (2): 881–94.

González, Beatriz, Maria Navarro-Jiménez, María José Alonso-De Gennaro, Sanne Marcia Jansen, Isabel Granada, Manuel Perucho, and Sergio Alonso. 2021. “Somatic Hypomethylation of Pericentromeric SST1 Repeats and Tetraploidization in Human Colorectal Cancer Cells.” Cancers 13 (21). https://doi.org/10.3390/cancers13215353.

Grant, Charles E., Timothy L. Bailey, and William Stafford Noble. 2011. “FIMO: Scanning for Occurrences of a given Motif.” Bioinformatics 27 (7): 1017–18.

Greig, G. M., P. E. Warburton, and H. F. Willard. 1993. “Organization and Evolution of an Alpha *Satellite DNA Subset Shared by Human Chromosomes 13 and 21*.” Journal of Molecular Evolution 37 (5): 464–75.

Guarracino, Andrea, Simon Heumos, Sven Nahnsen, Pjotr Prins, and Erik Garrison. 2022. “ODGI: Understanding Pangenome Graphs.” Bioinformatics, May. https://doi.org/10.1093/bioinformatics/btac308.

Guarracino, Andrea, Njagi Mwaniki, Santiago Marco-Sola, and Erik Garrison. 2021. Wfmash: A Pangenome-Scale Pairwise Aligner. https://doi.org/10.5281/zenodo.6949373.

Guissani, U., B. Facchinetti, G. Cassina, and O. Zuffardi. 1996. “Mitotic Recombination among Acrocentric Chromosomes’ Short Arms.” Annals of Human Genetics 60 (2): 91–97.

Hamerton, J. L., N. Canning, M. Ray, and S. Smith. 1975. “A Cytogenetic Survey of 14,069 Newborn Infants. I. Incidence of Chromosome Abnormalities.” Clinical Genetics 8 (4): 223–43.

Helena Mangs, A., and Brian J. Morris. 2007. “The Human Pseudoautosomal Region (PAR): Origin, Function and Future.” Current Genomics 8 (2): 129–36.

Henderson, A. S., D. Warburton, and K. C. Atwood. 1973. “Letter: Ribosomal DNA Connectives between Human Acrocentric Chromosomes.” Nature 245 (5420): 95–97.

Hickey, Glenn, David Heller, Jean Monlong, Jonas A. Sibbesen, Jouni Sirén, Jordan Eizenga, Eric T. Dawson, Erik Garrison, Adam M. Novak, and Benedict Paten. 2020. “Genotyping Structural Variants in Pangenome Graphs Using the vg Toolkit.” Genome Biology 21 (1): 35.

Holm, Preben Bach, and Søren Wilken Rasmussen. 1977. “Human Meiosis I. The Human Pachytene Karyotype Analyzed by Three Dimensional Reconstruction of the Synaptonemal Complex.” Carlsberg Research Communications 42 (4): 283.

Hoyt, Savannah J., Jessica M. Storer, Gabrielle A. Hartley, Patrick G. S. Grady, Ariel Gershman, Leonardo G. de Lima, Charles Limouse, et al. 2022. “From Telomere to Telomere: The Transcriptional and Epigenetic State of Human Repeat Elements.” Science 376 (6588): eabk3112.

Huttley, G. A., M. W. Smith, M. Carrington, and S. J. O’Brien. 1999. “A Scan for Linkage Disequilibrium across the Human Genome.” Genetics 152 (4): 1711–22. “Igraph Citation Info.” n.d. Accessed June 25, 2022. https://cran.r-project.org/web/packages/igraph/citation.html.

International Human Genome Sequencing Consortium. 2001. “Initial Sequencing and Analysis of the Human Genome.” Nature 409 (6822): 860–921.

Jarmuz-Szymczak, Malgorzata, Joanna Janiszewska, Krzysztof Szyfter, and Lisa G. Shaffer. 2014. “Narrowing the Localization of the Region Breakpoint in Most Frequent Robertsonian *Translocations*.” Chromosome Research: An International Journal on the Molecular, Supramolecular and Evolutionary Aspects of Chromosome Biology 22 (4): 517–32.

Jørgensen, A. L., C. J. Bostock, and A. L. Bak. 1987. “Homologous Subfamilies of Human *Alphoid Repetitive DNA on Different Nucleolus Organizing Chromosomes*.” Proceedings of the National Academy of Sciences of the United States of America 84 (4): 1075–79.

Jørgensen, A. L., S. Kølvraa, C. Jones, and A. L. Bak. 1988. “A Subfamily of Alphoid Repetitive DNA Shared by the NOR-Bearing Human Chromosomes 14 and 22.” Genomics 3 (2): 100–109.

Kaniecki, Kyle, Luisina De Tullio, and Eric C. Greene. 2018. “A Change of View: Homologous Recombination at Single-Molecule Resolution.” Nature Reviews. Genetics 19 (4): 191–207.

Kinene, T., J. Wainaina, S. Maina, and L. M. Boykin. 2016. “Rooting Trees, Methods for.” In Encyclopedia of Evolutionary Biology, edited by Richard M. Kliman, 489–93. Oxford: Academic Press.

Kobayashi, Takehiko. 2011. “Regulation of Ribosomal RNA Gene Copy Number and Its Role in Modulating Genome Integrity and Evolutionary Adaptability in Yeast.” Cellular and Molecular Life Sciences: CMLS 68 (8): 1395–1403.

Liao, Wen-Wei, Mobin Asri, Jana Ebler, Daniel Doerr, Marina Haukness, Glenn Hickey, Shuangjia Lu, et al. 2022. “A Draft Human Pangenome Reference.” bioRxiv. https://doi.org/10.1101/2022.07.09.499321.

Li, Heng, Xiaowen Feng, and Chong Chu. 2020. “The Design and Construction of Reference Pangenome Graphs with Minigraph.” Genome Biology 21 (1): 265.

Li, Na, and Matthew Stephens. 2003. “Modeling Linkage Disequilibrium and Identifying Recombination Hotspots Using Single-Nucleotide Polymorphism Data.” Genetics 165 (4): 2213–33.

Lindström, Mikael S., Deana Jurada, Sladana Bursac, Ines Orsolic, Jiri Bartek, and Sinisa Volarevic. 2018. “Nucleolus as an Emerging Hub in Maintenance of Genome Stability and Cancer Pathogenesis.” Oncogene 37 (18): 2351–66.

Logsdon, Glennis A., Mitchell R. Vollger, Pinghsun Hsieh, Yafei Mao, Mikhail A. Liskovykh, Sergey Koren, Sergey Nurk, et al. 2021. “The Structure, Function and Evolution of a Complete Human Chromosome 8.” Nature 593 (7857): 101–7.

Mack, H., and K. Swisshelm. 2013. “Robertsonian Translocations,” January, 301–5.

Marco-Sola, Santiago, Jordan M. Eizenga, Andrea Guarracino, Benedict Paten, Erik Garrison, and Miquel Moreto. 2022. “Optimal Gap-Affine Alignment in O(s) Space.” bioRxiv. https://doi.org/10.1101/2022.04.14.488380.

McClintock, Barbara. 1934. “The Relation of a Particular Chromosomal Element to the Development of the Nucleoli in Zea Mays.” Zeitschrift Für Zellforschung Und Mikroskopische Anatomie 21 (2): 294–326.

Nambiar, Mridula, and Gerald R. Smith. 2016. “Repression of Harmful Meiotic Recombination in Centromeric Regions.” Seminars in Cell & Developmental Biology 54 (June): 188–97.

Nishiyama, Rie, Lixin Qi, Koji Tsumagari, Karen Weissbecker, Louis Dubeau, Martin Champagne, Suresh Sikka, Hisaki Nagai, and Melanie Ehrlich. 2005. “A DNA Repeat, NBL2, Is Hypermethylated in Some Cancers but Hypomethylated in Others.” Cancer Biology & Therapy 4 (4): 440–48.

Nurk, Sergey, Sergey Koren, Arang Rhie, Mikko Rautiainen, Andrey V. Bzikadze, Alla Mikheenko, Mitchell R. Vollger, et al. 2022. “The Complete Sequence of a Human Genome.” Science 376 (6588): 44–53.

Paigen, Kenneth, and Petko M. Petkov. 2018. “PRDM9 and Its Role in Genetic Recombination.” Trends in Genetics: TIG 34 (4): 291–300.

Paten, Benedict, Jordan M. Eizenga, Yohei M. Rosen, Adam M. Novak, Erik Garrison, and Glenn Hickey. 2018. “Superbubbles, Ultrabubbles, and Cacti.” Journal of Computational Biology: A Journal of Computational Molecular Cell Biology 25 (7): 649–63.

Paten, Benedict, Adam M. Novak, Jordan M. Eizenga, and Erik Garrison. 2017. “Genome Graphs and the Evolution of Genome Inference.” Genome Research 27 (5): 665–76.

Peng, Zhen, Weichen Zhou, Wenqing Fu, Renqian Du, Li Jin, and Feng Zhang. 2015. “Correlation between Frequency of Non-Allelic Homologous Recombination and Homology Properties: Evidence from Homology-Mediated CNV Mutations in the Human Genome.” Human Molecular Genetics 24 (5): 1225–33.

Purcell, Shaun, Benjamin Neale, Kathe Todd-Brown, Lori Thomas, Manuel A. R. Ferreira, David Bender, Julian Maller, et al. 2007. “PLINK: A Tool Set for Whole-Genome Association and Population-Based Linkage Analyses.” American Journal of Human Genetics 81 (3): 559–75.

Quinlan, Aaron R., and Ira M. Hall. 2010. “BEDTools: A Flexible Suite of Utilities for Comparing Genomic Features.” Bioinformatics 26 (6): 841–42.

Rautiainen, Mikko, and Tobias Marschall. 2020. “GraphAligner: Rapid and Versatile Sequence-to-Graph Alignment.” Genome Biology 21 (1): 253.

Rautiainen, Mikko, Sergey Nurk, Brian P. Walenz, Glennis A. Logsdon, David Porubsky, Arang Rhie, Evan E. Eichler, Adam M. Phillippy, and Sergey Koren. 2022. “Verkko: Telomere-to-Telomere Assembly of Diploid Chromosomes.” bioRxiv. https://doi.org/10.1101/2022.06.24.497523.

Rhie, Arang, Sergey Nurk, Monika Cechova, Savannah J. Hoyt, Dylan J. Taylor, Nicolas Altemose, Paul W. Hook, et al. 2022. “The Complete Sequence of a Human Y Chromosome.” bioRxiv. https://doi.org/10.1101/2022.12.01.518724.

Roberts, Paul A. 1965. “Difference in the Behaviour of Eu-and Hetero-Chromatin: Crossing-Over.” Nature 205 (4972): 725–26.

Ross, Mark T., Darren V. Grafham, Alison J. Coffey, Steven Scherer, Kirsten McLay, Donna Muzny, Matthias Platzer, et al. 2005. “The DNA Sequence of the Human X Chromosome.” Nature 434 (7031): 325–37.

Samuelsson, Johanna K., Gabrijela Dumbovic, Cristian Polo, Cristina Moreta, Andreu Alibés, Tatiana Ruiz-Larroya, Pepita Giménez-Bonafé, Sergio Alonso, Sonia-V Forcales, and Perucho Manuel. 2017. “Helicase Lymphoid-Specific Enzyme Contributes to the Maintenance of Methylation of SST1 Pericentromeric Repeats That Are Frequently Demethylated in Colon Cancer and Associate with Genomic Damage.” Epigenomes 1 (1). https://doi.org/10.3390/epigenomes1010002.

Sirén, Jouni, Jean Monlong, Xian Chang, Adam M. Novak, Jordan M. Eizenga, Charles Markello, Jonas A. Sibbesen, et al. 2021. “Pangenomics Enables Genotyping of Known Structural Variants in 5202 Diverse Genomes.” Science 374 (6574): abg8871.

Sluis, Marjolein van, Michael Ó. Gailín, Joseph G. W. McCarter, Hazel Mangan, Alice Grob, and Brian McStay. 2019. “Human NORs, Comprising rDNA Arrays and Functionally Conserved Distal Elements, Are Located within Dynamic Chromosomal Regions.” Genes & Development 33 (23-24): 1688–1701.

Spinner, N. B. 2013. “Chromosome Banding.” In Brenner’s Encyclopedia of Genetics (Second Edition), edited by Stanley Maloy and Kelly Hughes, 546–48. San Diego: Academic Press.

Sullivan, B. A., L. S. Jenkins, E. M. Karson, J. Leana-Cox, and S. Schwartz. 1996. “Evidence for Structural Heterogeneity from Molecular Cytogenetic Analysis of Dicentric Robertsonian Translocations.” American Journal of Human Genetics 59 (1): 167–75.

Taliun, Daniel, Daniel N. Harris, Michael D. Kessler, Jedidiah Carlson, Zachary A. Szpiech, Raul Torres, Sarah A. Gagliano Taliun, et al. 2021. “Sequencing of 53,831 Diverse Genomes from the NHLBI TOPMed Program.” Nature 590 (7845): 290–99.

Traag, V. A., L. Waltman, and N. J. van Eck. 2019. “From Louvain to Leiden: Guaranteeing Well-Connected Communities.” Scientific Reports 9 (1): 5233.

Tremblay, Deanna C., Graham Alexander Jr, Shawn Moseley, and Brian P. Chadwick. 2010. “Expression, Tandem Repeat Copy Number Variation and Stability of Four Macrosatellite Arrays in the Human Genome.” BMC Genomics 11 (November): 632.

Veerappa, Avinash M., Prakash Padakannaya, and Nallur B. Ramachandra. 2013. “Copy Number Variation-Based Polymorphism in a New Pseudoautosomal Region 3 (PAR3) of a *Human X-Chromosome-Transposed Region (XTR) in the Y Chromosome*.” Functional & Integrative Genomics 13 (3): 285–93.

Zickler, Denise, and Nancy Kleckner. 2015. “Recombination, Pairing, and Synapsis of Homologs *during Meiosis*.” Cold Spring Harbor Perspectives in Biology. https://doi.org/10.1101/cshperspect.a016626.

