## Supplementary Figures and captions for "Recombination between heterologous human acrocentric chromosomes"

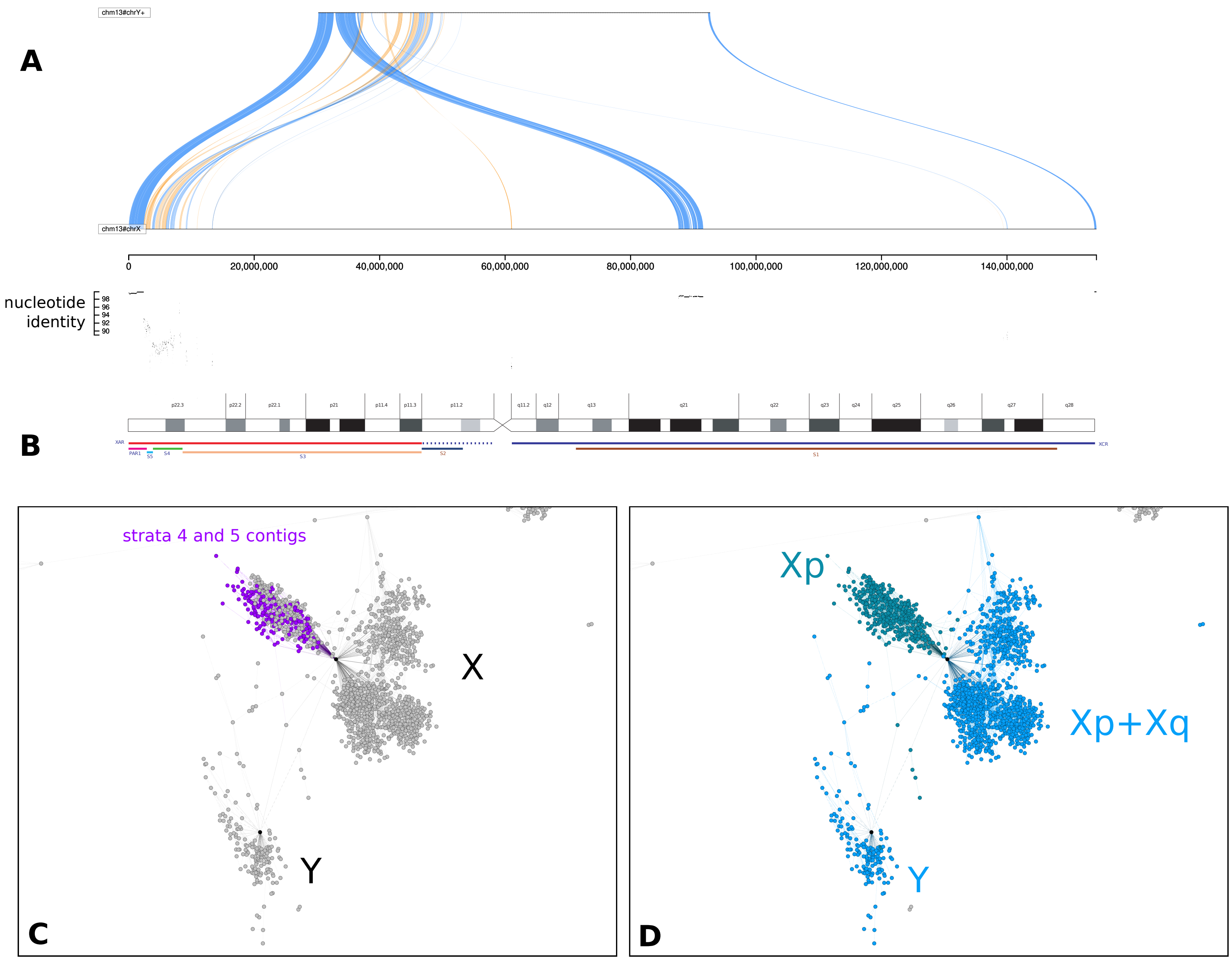


Supplementary Figure 1: (A) Visualization with Saffire (https://mrvollger.github.io/SafFire/) of the alignment between T2T-CHM13 X and Y reveals that strata 5 and 4 feature low identity ( 90%), numerous inversions, and some rearrangements; (B) X chromosome ideogram according to (Ross et al. 2005). On the bottom, its evolutionary domains: the X-added region (XAR), the X-conserved region (XCR; dotted region in proximal Xp does not appear to be part of the XCR), the pseudoautosomal region PAR1, and evolutionary strata S5–S1. (C) The reduced all-to-all mapping graph of HPRCy1 versus itself, with contigs represented as nodes and mappings as edges, rendered in Gephi (Bastian, Heymann, and Jacomy 2009). In red contigs covering the evolutionary strata 5 and 4 on chromosome X; (D) Coloring the reduced homology mapping graph in C with community assignments. Panels C and D use the same layout as Figure 1 but focus only on the X and Y region of the visualization.

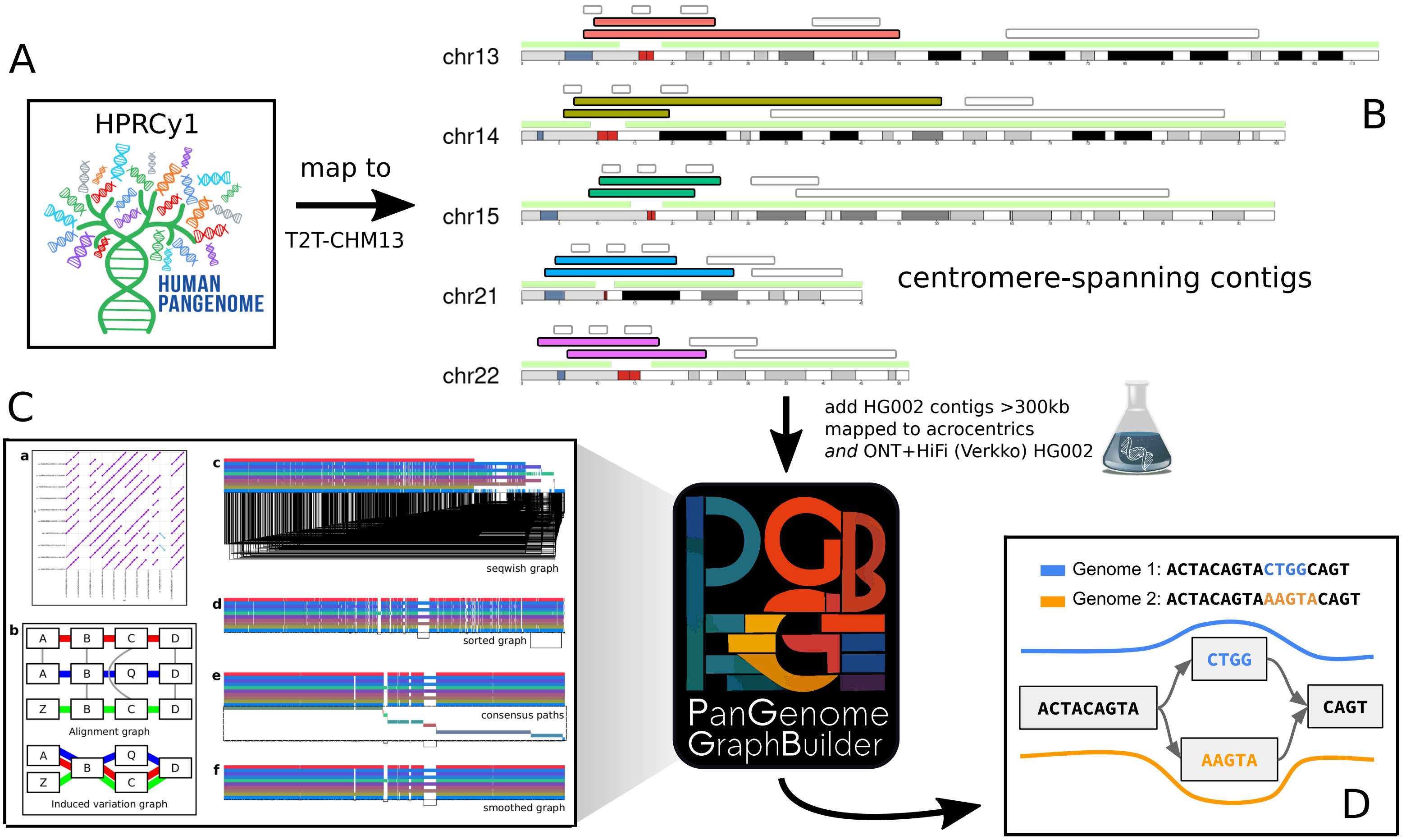


Supplementary Figure 2: An overview of our approach to build a PVG for HPRCy1 contigs that can be anchored to a specific acrocentric q-arm. (A) As input, we take the entire HPRCy1 and map it to T2T-CHM13. (B) This yields mappings to acrocentric chromosomes, which we filter to select contigs that map across the centromeres (red cytobands) between non-centromeric regions (over-labeled green). We include two HG002 assemblies based on standard HiFi (from HPRCy1) and on both HiFi and ONT data (from Verkko). (C) We then apply PGGB to build a PVG from the HPRCy1-acro collection. PGGB first obtains an all-to-all alignment of the input (C.a.), which is converted to a variation graph with SEQWISH (Garrison and Guarracino 2022) (C.b.), then normalized with sorting and multiple sequence alignment steps in SMOOTHXG (C.c-f). (D) The resulting PVG expresses genomes as paths, or walks, through a common sequence graph. This model thus contains all input sequences and their relative alignments to all others—in the example we see a CTGG/AAGTA block substitution between genomes 1 and 2.

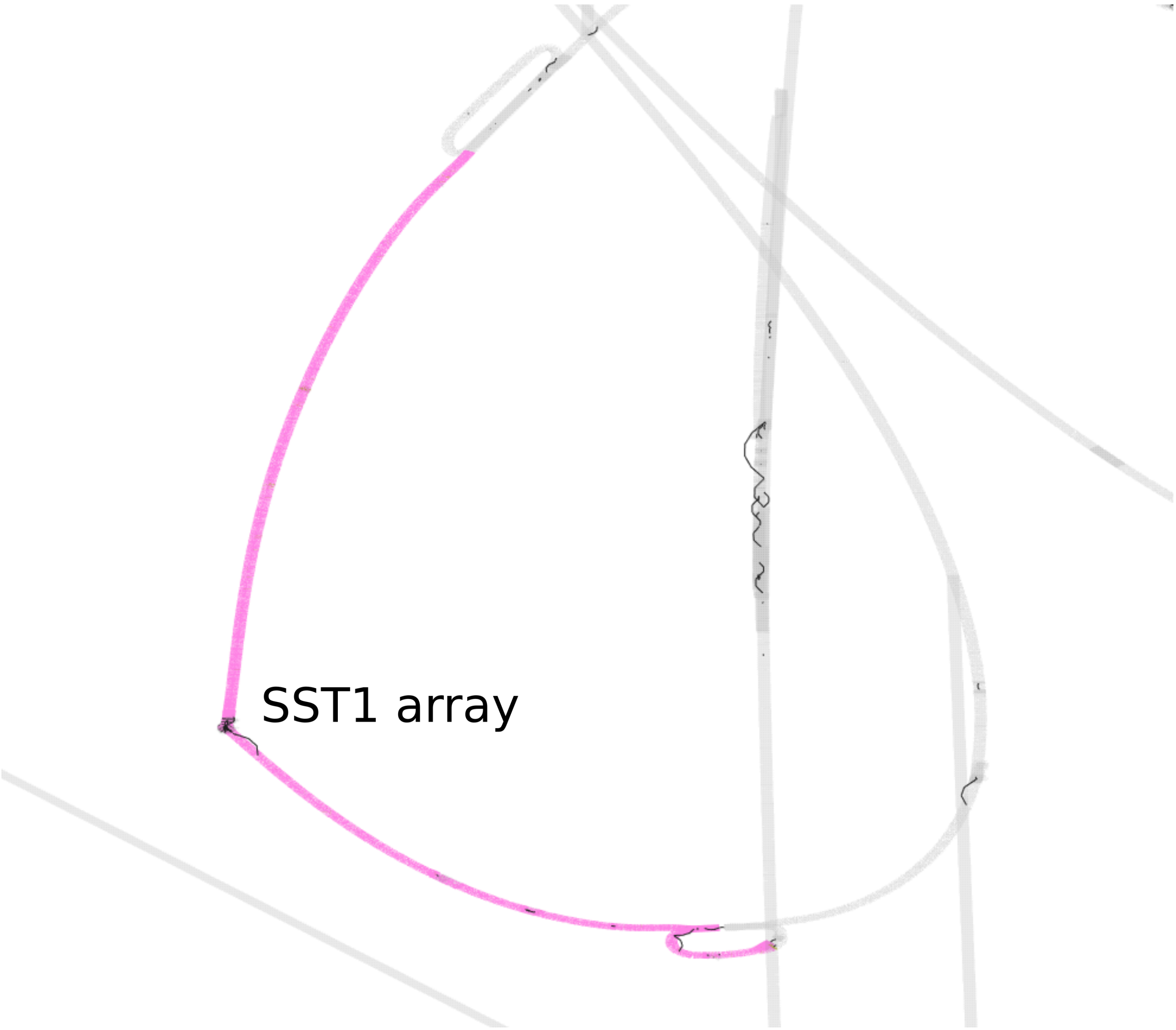


Supplementary Figure 3: Regions of a PVG built from centromere-spanning HPRCy1 contigs plus T2T-CHM13 and GRCh38 references. We apply GFAESTUS to visualize the 2D layout generated by ODGI in the PGGB pipeline. We focus on the segmentally duplicated core centered in the SST1 array (labeled).

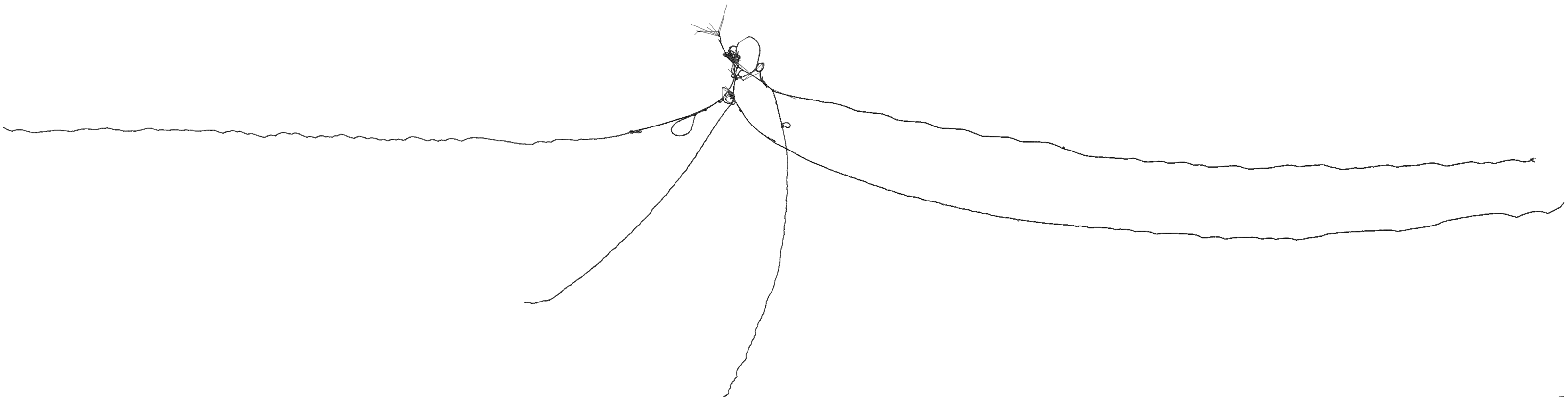

Supplementary Figure 4: A PVG built from centromere-spanning HPRCy1 contigs, without embedding the T2T-CHM13 and GRCh38 references. We apply ODGI to visualize the 2D layout generated in the PGGB pipeline. This renders sequences and chains of small variants as linear structures, while repeats caused by segmental duplications, inversions, and other structural variants tend to form loops and tangles. The acrocentric q-arms are almost completely separated, while the p-arms unite in a structure adjacent to the rDNA array.

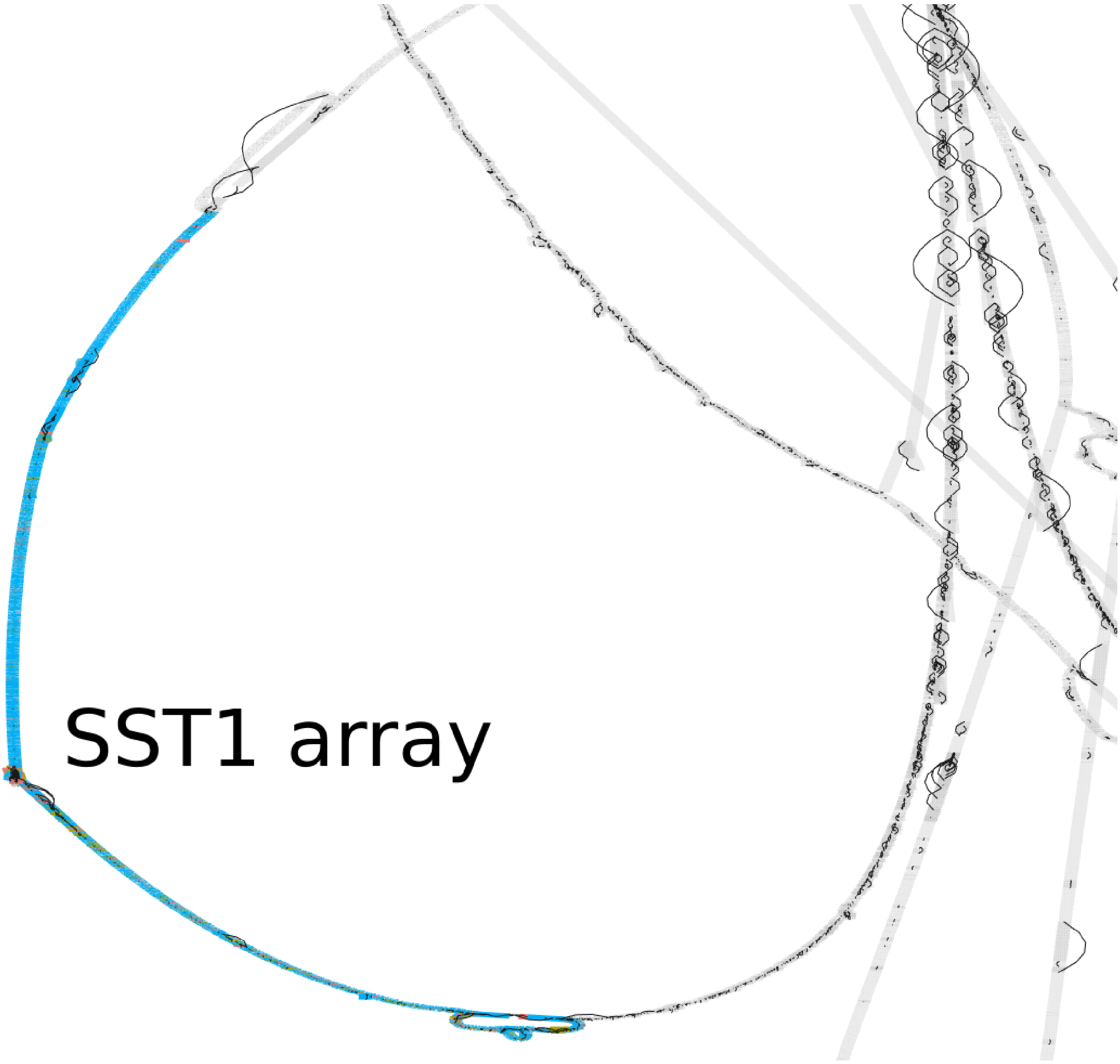


Supplementary Figure 5: Regions of a PVG built from centromere-spanning HPRCy1 contigs without the T2T-CHM13 and GRCh38 references. We apply GFAESTUS to visualize the 2D layout generated by ODGI in the PGGB pipeline. We focus on the segmentally duplicated core centered in the SST1 array (labeled).

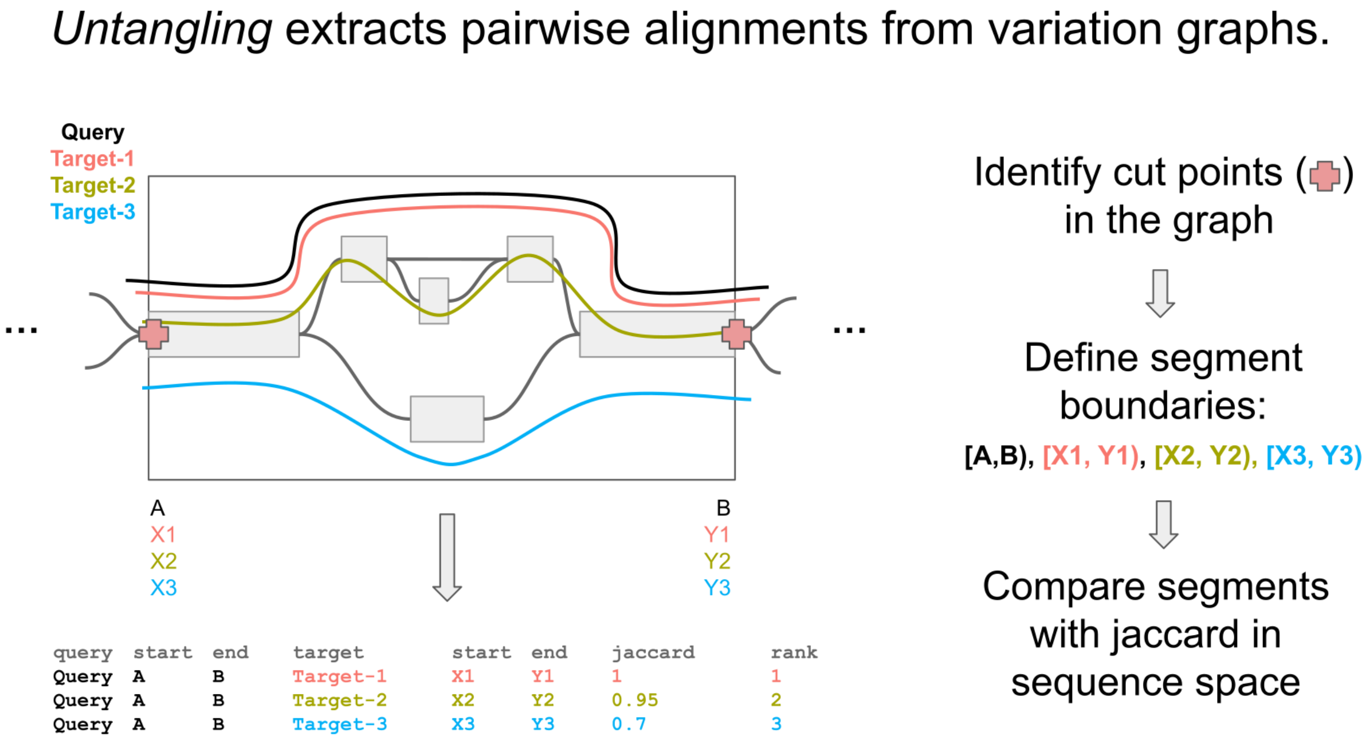


Supplementary Figure 6: Scheme of the graph untangling. We applied ODGI UNTANGLE to obtain a mapping from segments of all PVG paths onto T2T-CHM13. The segmentation cuts the graph into regular-sized regions whose boundaries occur at structural variant breakpoints. For each query subpath through a graph segment, we use a Jaccard metric over the sequence space of the subpaths to find the best-matching reference segment.

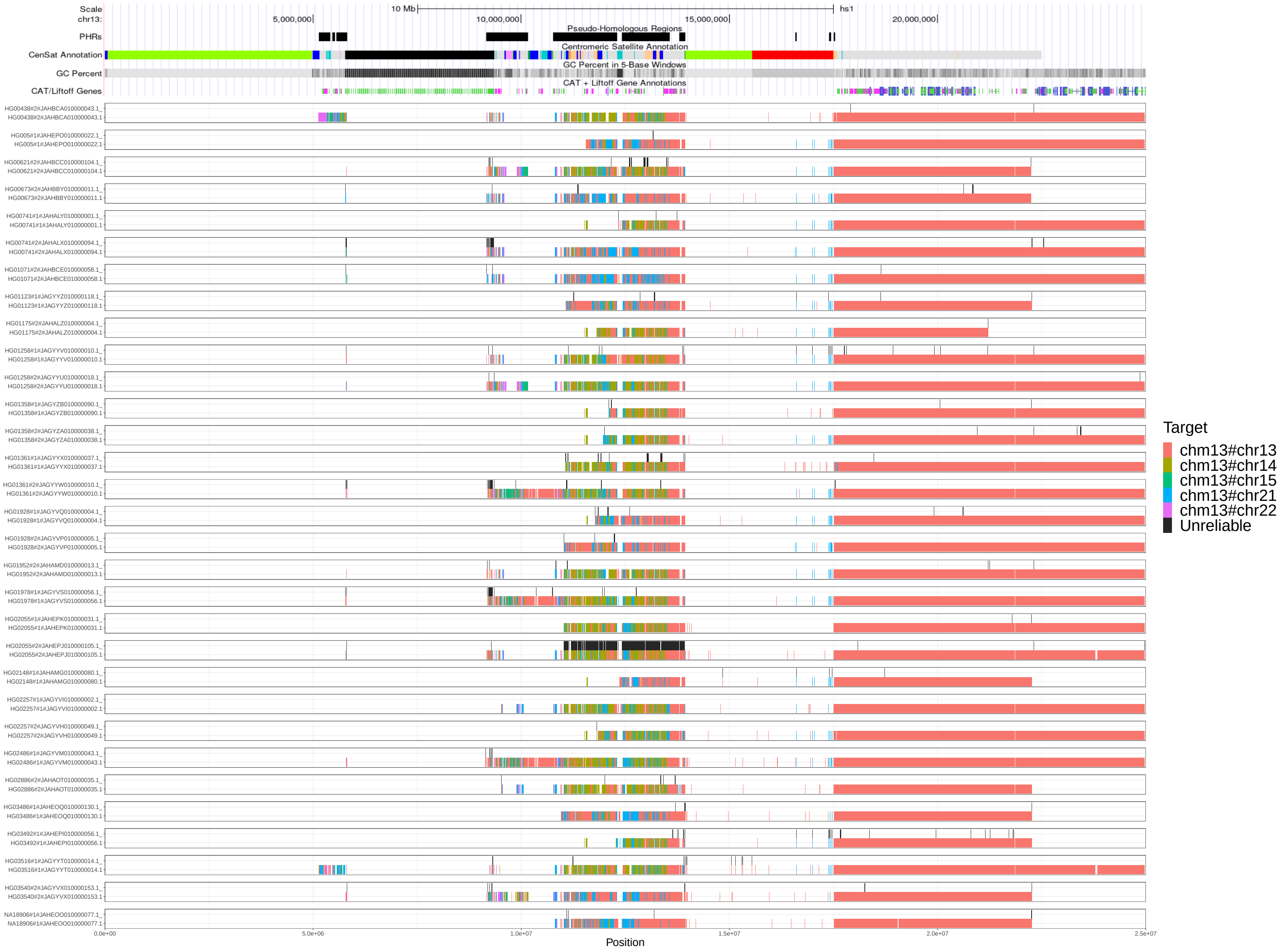


Supplementary Figure 7: Untangling of HPRCy1-acro's sequences belonging to chromosome 13 versus T2T-CHM13. We display all mappings above 90% estimated pairwise identity, without removing those covering regions classified as unreliable. Above each contig, we display the unreliable regions in black.

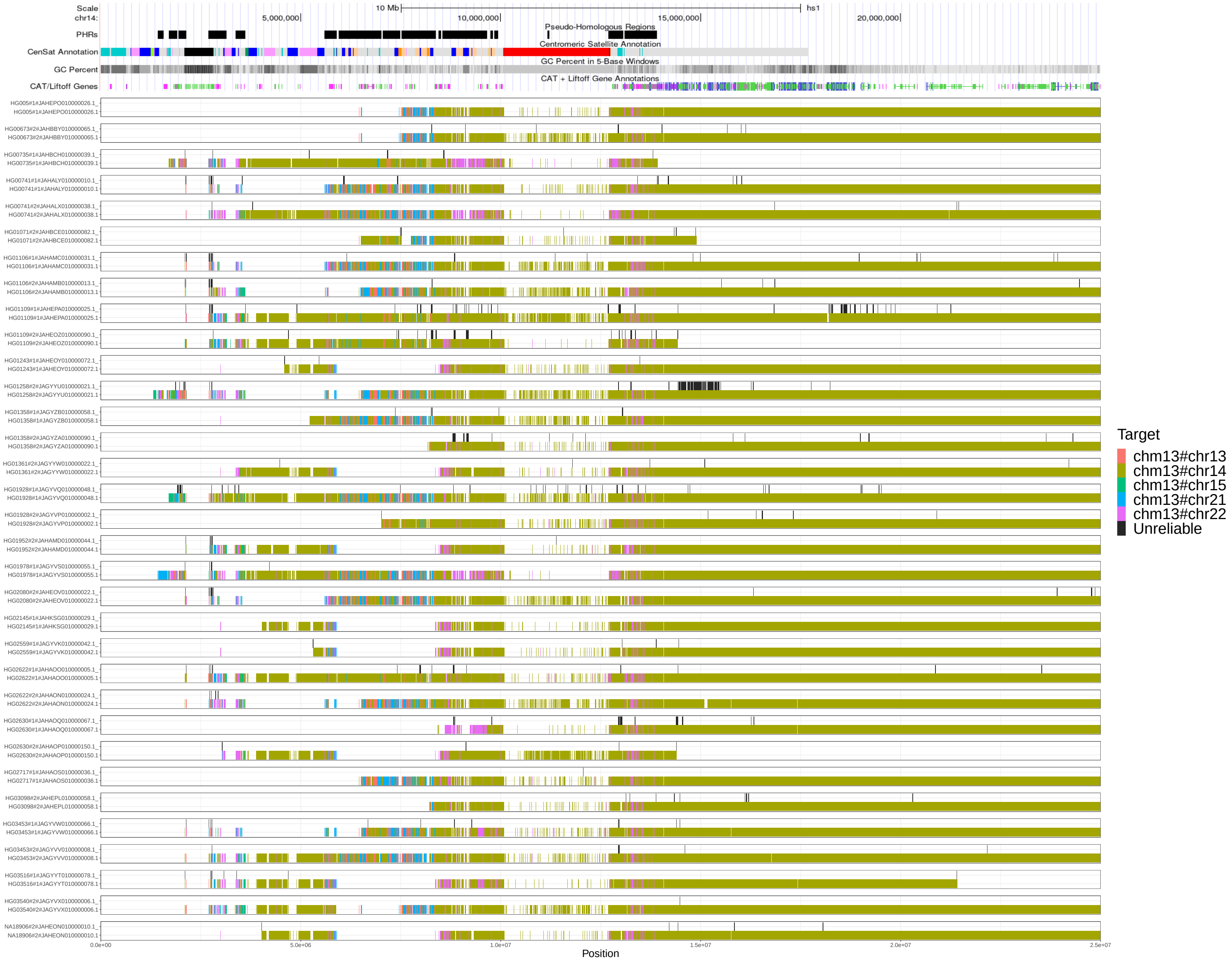


Supplementary Figure 8: Untangling of HPRCy1-acro's sequences belonging to chromosome 14 versus T2T-CHM13. We display all mappings above 90% estimated pairwise identity, without removing those covering regions classified as unreliable. Above each contig, we display the unreliable regions in black.

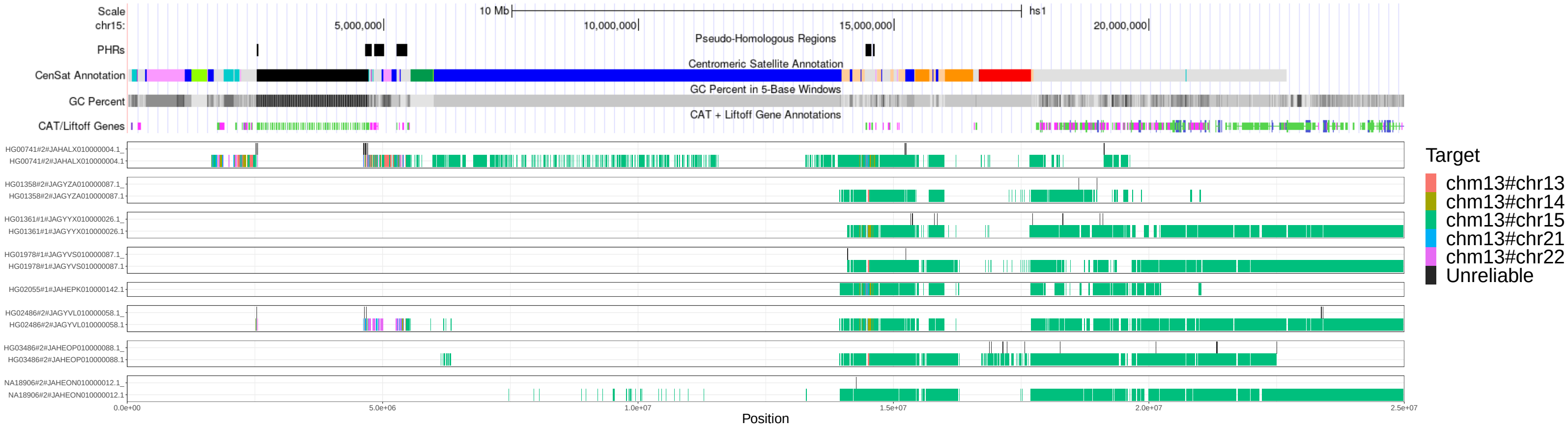


Supplementary Figure 9: Untangling of HPRCy1-acro's sequences belonging to chromosome 15 versus T2T-CHM13. We display all mappings above 90% estimated pairwise identity, without removing those covering regions classified as unreliable. Above each contig, we display the unreliable regions in black.

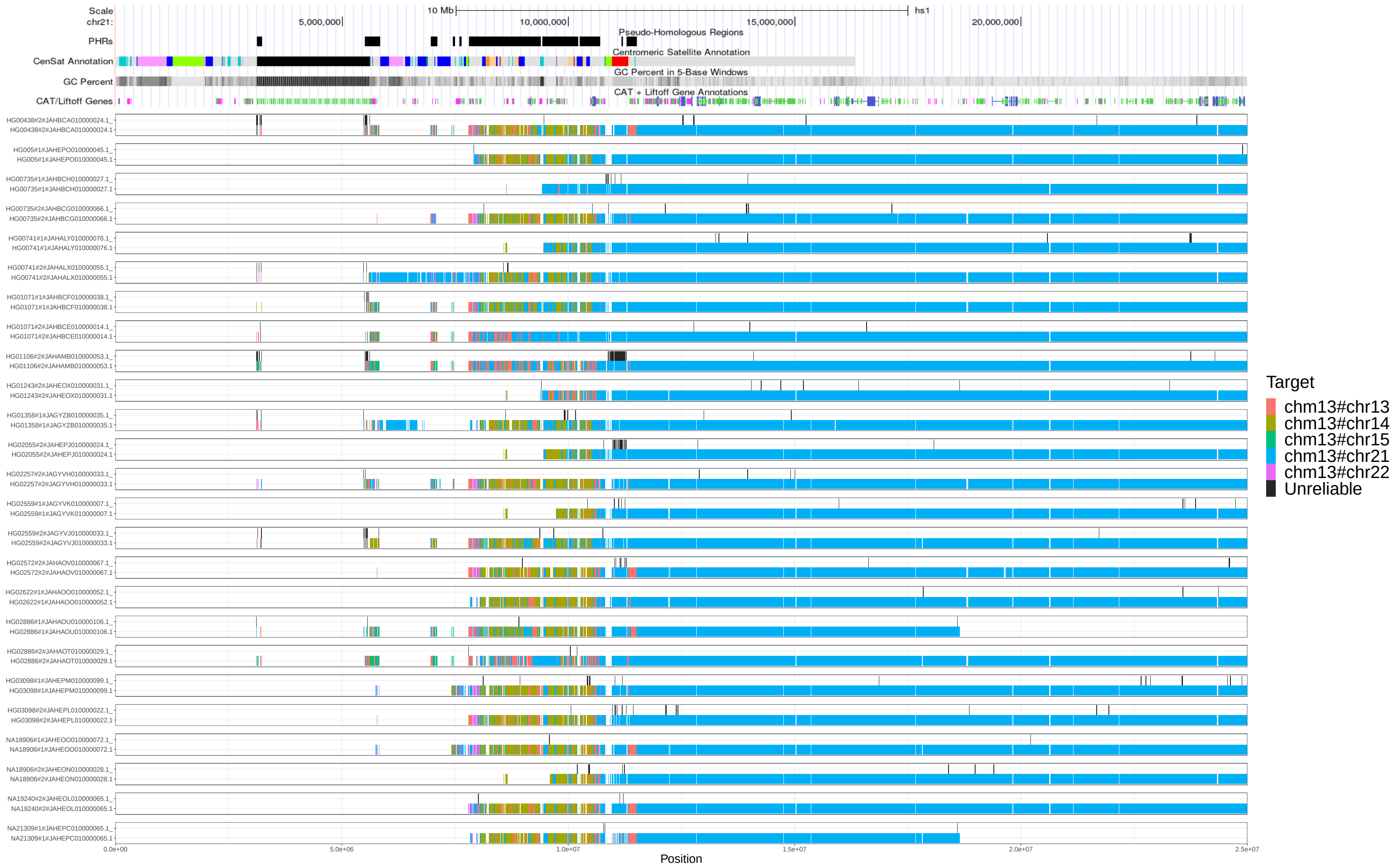


Supplementary Figure 10: Untangling of HPRCy1-acro's sequences belonging to chromosome 21 versus T2T-CHM13. We display all mappings above 90% estimated pairwise identity, without removing those covering regions classified as unreliable. Above each contig, we display the unreliable regions in black.

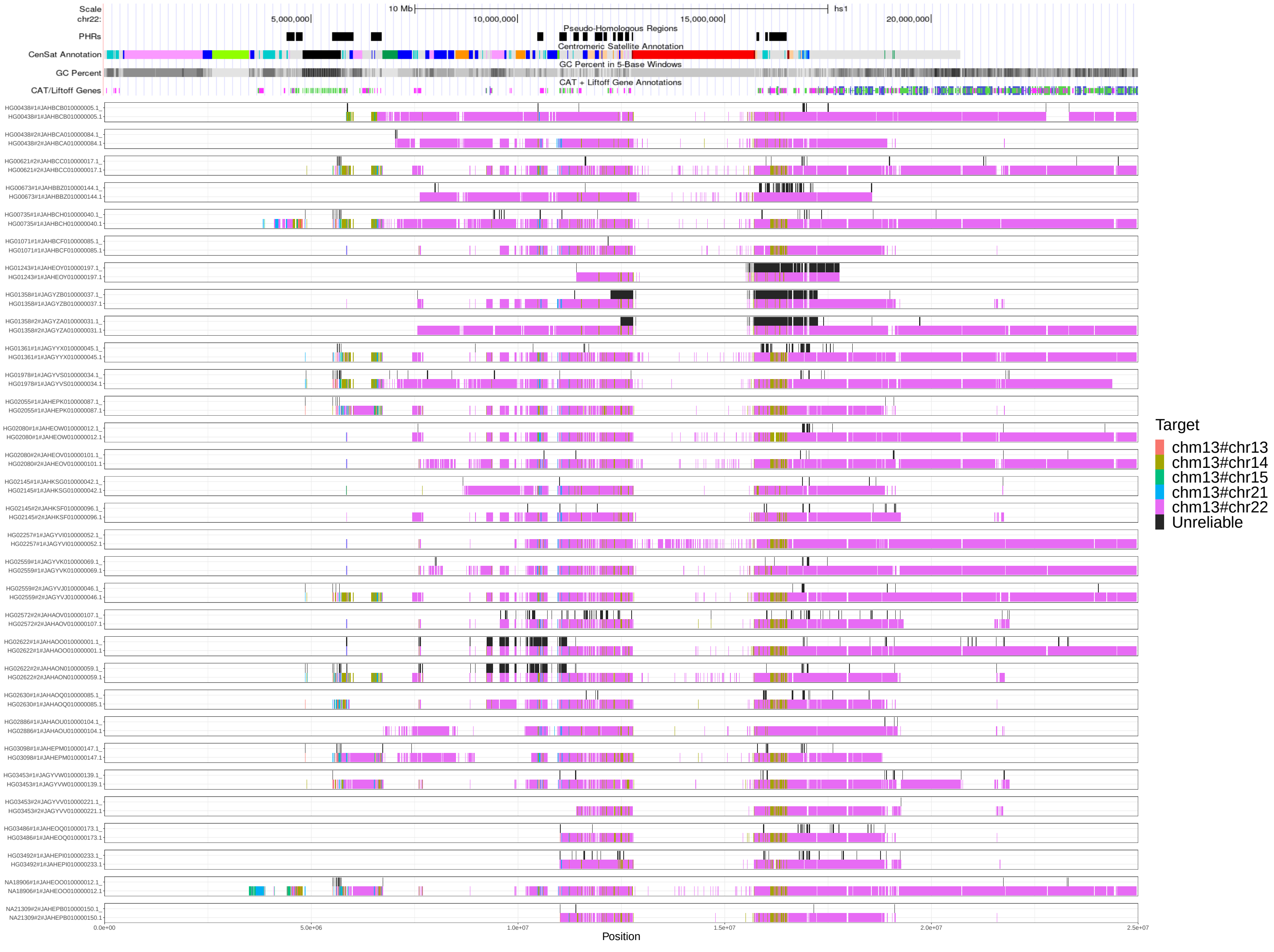


Supplementary Figure 11: Untangling of HPRCy1-acro's sequences belonging to chromosome 22 versus T2T-CHM13. We display all mappings above 90% estimated pairwise identity, without removing those covering regions classified as unreliable. Above each contig, we display the unreliable regions in black.

| | |
|---|---|
| Active αSat HOR (hor ... L) | red |
| Inactive αSat HOR (hor) | orange |
| Divergent αSat HOR (dhor) | dark red |
| Monomeric αSat (mon) | peach/yellow |
| Classical Human Satellite 1A (hsat1A) | light green |
| Classical Human Satellite 1B (hsat1B) | dark green |
| Classical Human Satellite 2 (hsat2) | light blue |
| Classical Human Satellite 3 (hsat3) | blue |
| Beta Satellite (bsat) | pink |
| Gamma Satellite (gsat) | purple |
| Other centromeric satellites (censat) | teal |
| Centromeric transition regions (ct) | grey |

Supplementary Figure 12: Centromeric Satellite Annotation (CenSat Annotation) track legend.

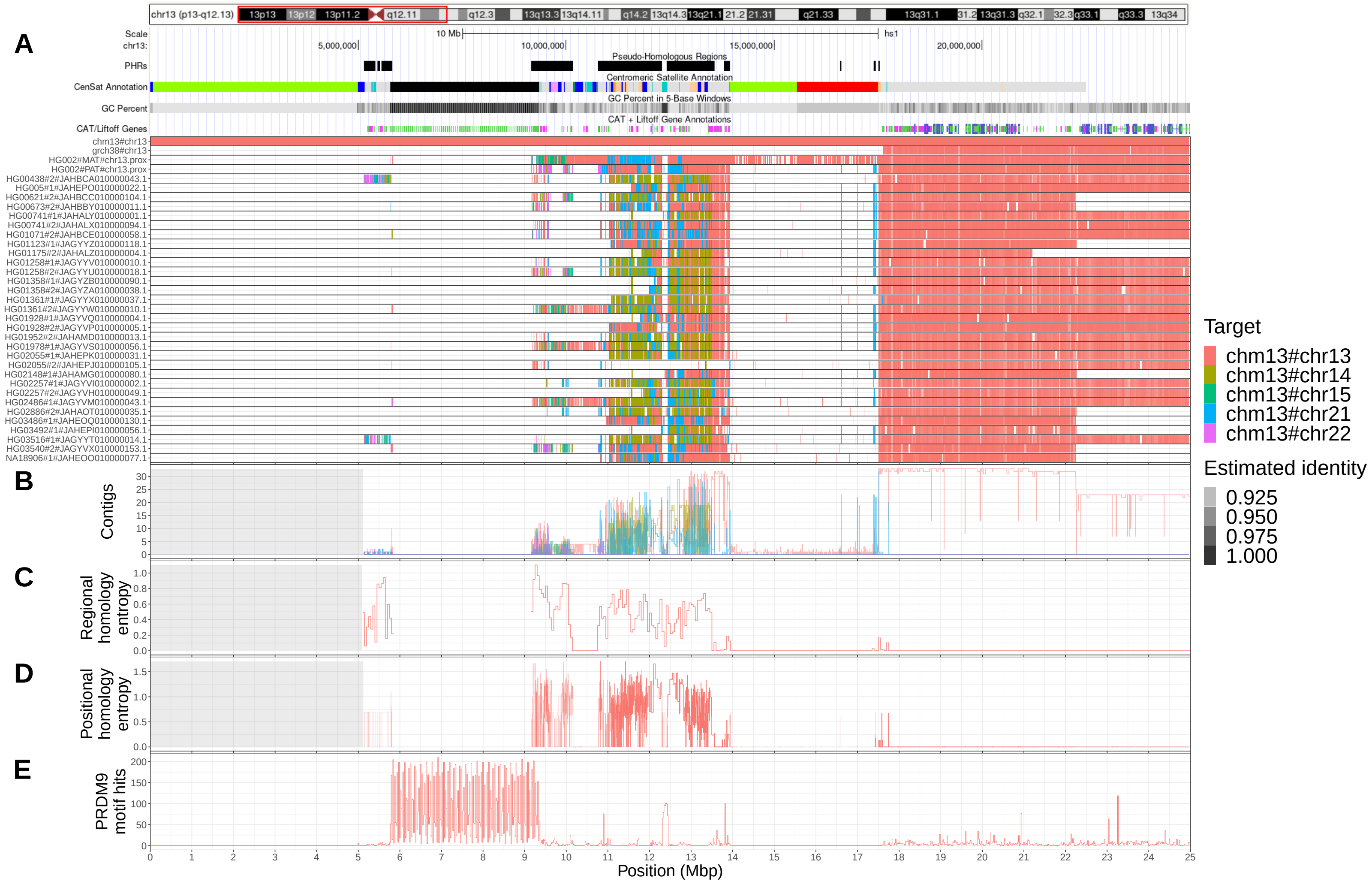

Supplementary Figure 13: (A) We focus on the first 25 Mbp of chromosome 13 shown here as a red box over T2T-CHM13 cytobands. Pseudo-homologous regions (PHRs), where diverse sets of acrocentric chromosomes recombine, are highlighted relative to T2T-CHM13 genome annotations for repeats, GC percentage, and genes. Above, we indicate regions of interest described in the main text: rDNA, SST1 array, centromere, and q-arm. Below, we show T2T-CHM13-relative homology mosaics for each chromosome 13 matched contig from HPRCy1-acro, with the most-similar reference chromosome at each region shown using the given colors (Target). (B) Aggregated untangle results in the SAACs. For each acrocentric chromosome, we show the count of its HPRCy1 q-arm-anchored contigs mapping itself and all other acrocentrics (Contigs), (C) as well as the regional (50kbp) untangle entropy metric (Regional homology entropy) computed over the contigs' T2T-CHM13-relative untanglings. (D) By considering the multiple untangling of each HPRCy1-acro contig, we develop a point-wise metric that captures diversity in T2T-CHM13-relative homology patterns (Positional homology entropy), leading to our definition of the PHRs. (E) The patterns of homology mosaicism suggest ongoing recombination exchange in the SAACs. A scan over T2T-CHM13 reveals that the rDNA units are enriched for PRDM9 binding motifs, and thus may host frequent double stranded breaks during meiosis. In (B-D) a gray background indicates regions with missing data due to the lack of non-T2T-CHM13 contigs. We provide the Centromeric Satellite Annotation (CenSat Annotation) track legend in Supplementary Figure 12.

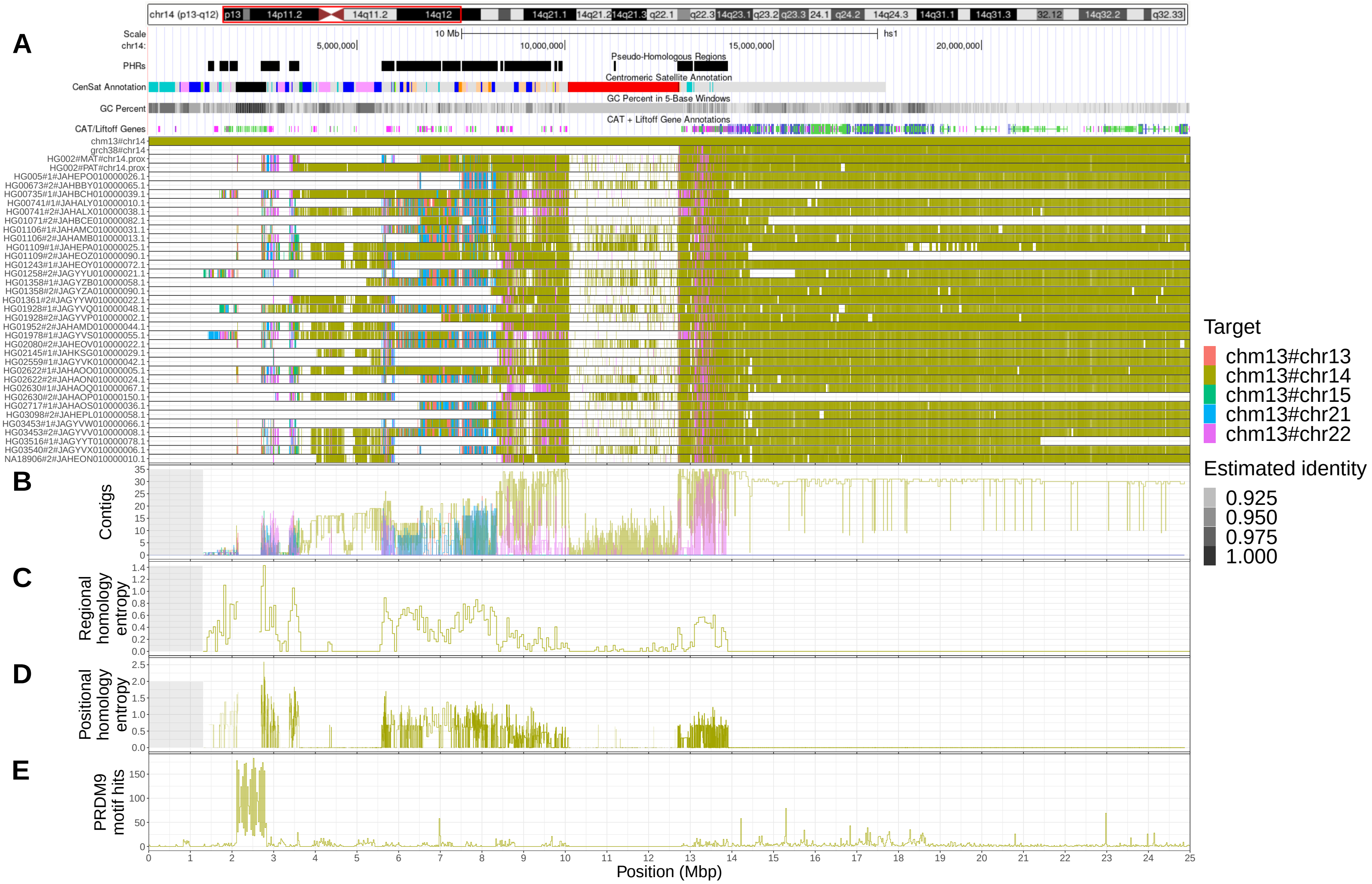


Supplementary Figure 14: (A) We focus on the first 25 Mbp of chromosome 14 shown here as a red box over T2T-CHM13 cytobands. Pseudo-homologous regions (PHRs), where diverse sets of acrocentric chromosomes recombine, are highlighted relative to T2T-CHM13 genome annotations for repeats, GC percentage, and genes. Above, we indicate regions of interest described in the main text: rDNA, SST1 array, centromere, and q-arm. Below, we show T2T-CHM13-relative homology mosaics for each chromosome 13 matched contig from HPRCy1-acro, with the most-similar reference chromosome at each region shown using the given colors (Target). (B) Aggregated untangle results in the SAACs. For each acrocentric chromosome, we show the count of its HPRCy1 q-arm-anchored contigs mapping itself and all other acrocentrics (Contigs), (C) as well as the regional (50kbp) untangle entropy metric (Regional homology entropy) computed over the contigs' T2T-CHM13-relative untanglings. (D) By considering the multiple untangling of each HPRCy1-acro contig, we develop a point-wise metric that captures diversity in T2T-CHM13-relative homology patterns (Positional homology entropy), leading to our definition of the PHRs. (E) The patterns of homology mosaicism suggest ongoing recombination exchange in the SAACs. A scan over T2T-CHM13 reveals that the rDNA units are enriched for PRDM9 binding motifs, and thus may host frequent double stranded breaks during meiosis. In (B-D) a gray background indicates regions with missing data due to the lack of non-T2T-CHM13 contigs. We provide the Centromeric Satellite Annotation (CenSat Annotation) track legend in Supplementary Figure 12.

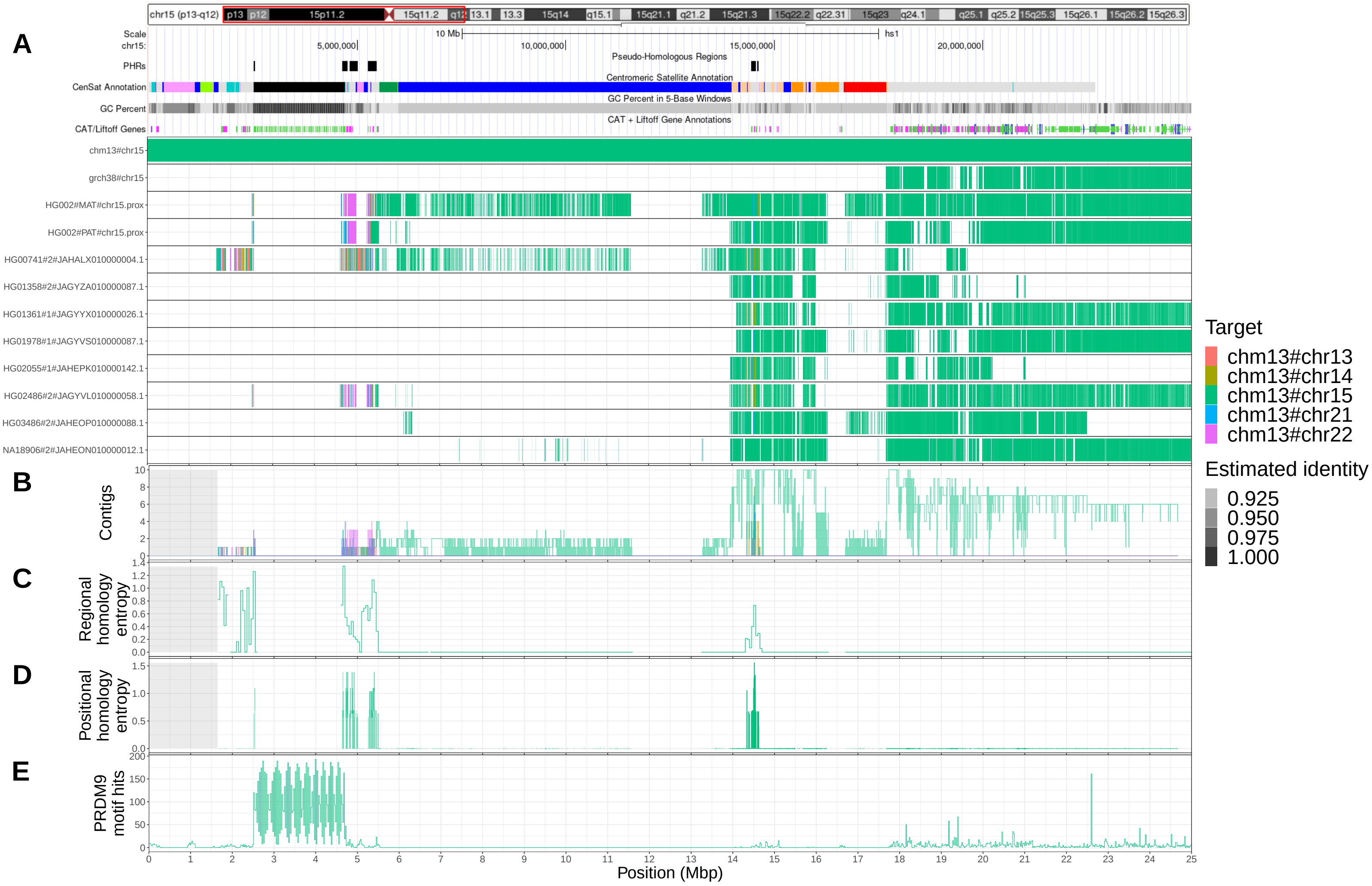


Supplementary Figure 15: (A) We focus on the first 25 Mbp of chromosome 15 shown here as a red box over T2T-CHM13 cytobands. Pseudo-homologous regions (PHRs), where diverse sets of acrocentric chromosomes recombine, are highlighted relative to T2T-CHM13 genome annotations for repeats, GC percentage, and genes. Above, we indicate regions of interest described in the main text: rDNA, SST1 array, centromere, and q-arm. Below, we show T2T-CHM13-relative homology mosaics for each chromosome 13 matched contig from HPRCy1-acro, with the most-similar reference chromosome at each region shown using the given colors (Target). (B) Aggregated untangle results in the SAACs. For each acrocentric chromosome, we show the count of its HPRCy1 q-arm-anchored contigs mapping itself and all other acrocentrics (Contigs), (C) as well as the regional (50kbp) untangle entropy metric (Regional homology entropy) computed over the contigs' T2T-CHM13-relative untanglings. (D) By considering the multiple untangling of each HPRCy1-acro contig, we develop a point-wise metric that captures diversity in T2T-CHM13-relative homology patterns (Positional homology entropy), leading to our definition of the PHRs. (E) The patterns of homology mosaicism suggest ongoing recombination exchange in the SAACs. A scan over T2T-CHM13 reveals that the rDNA units are enriched for PRDM9 binding motifs, and thus may host frequent double stranded breaks during meiosis. In (B-D) a gray background indicates regions with missing data due to the lack of non-T2T-CHM13 contigs. We provide the Centromeric Satellite Annotation (CenSat Annotation) track legend in Supplementary Figure 12.

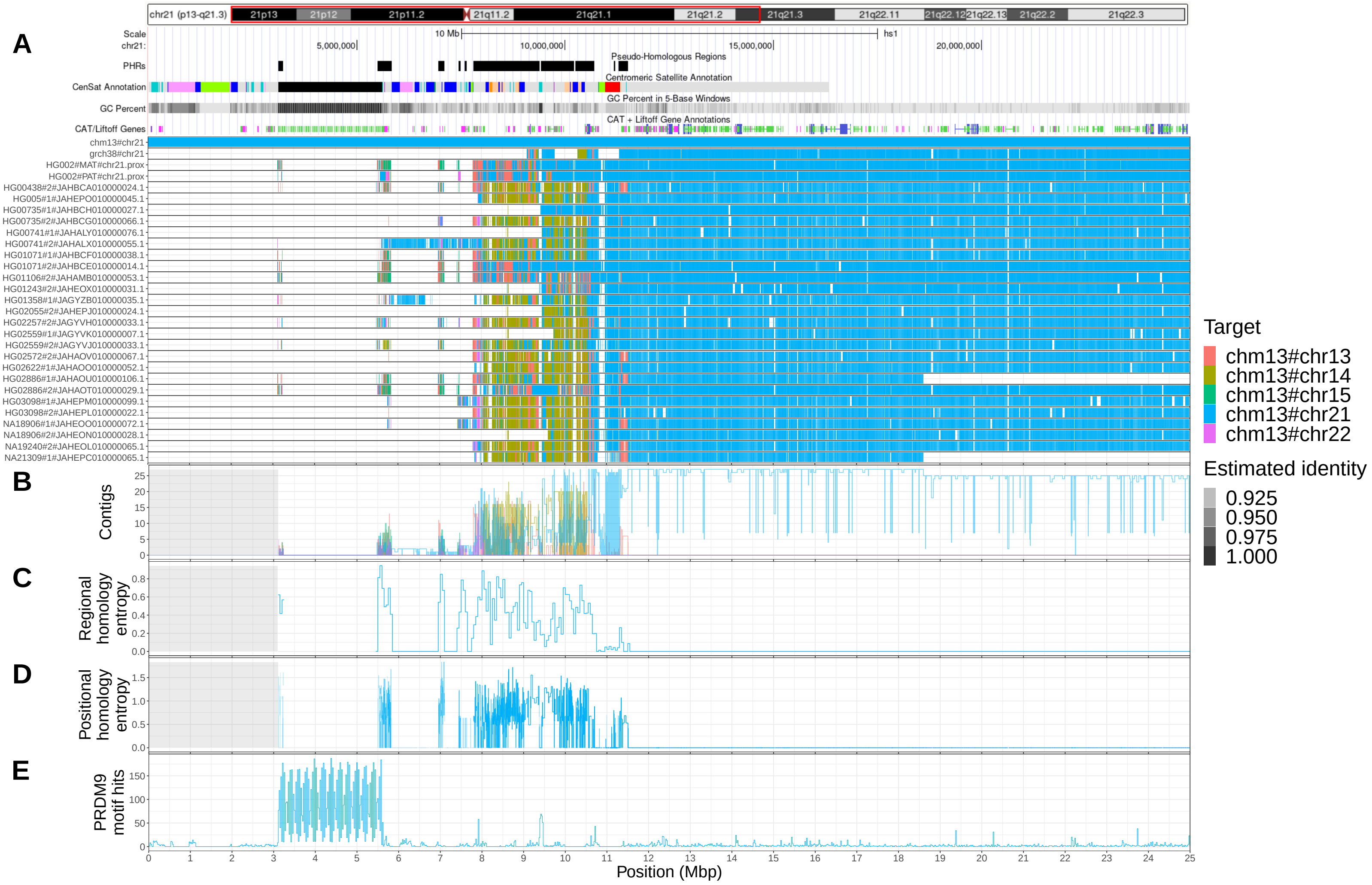


Supplementary Figure 16: (A) We focus on the first 25 Mbp of chromosome 21 shown here as a red box over T2T-CHM13 cytobands. Pseudo-homologous regions (PHRs), where diverse sets of acrocentric chromosomes recombine, are highlighted relative to T2T-CHM13 genome annotations for repeats, GC percentage, and genes. Above, we indicate regions of interest described in the main text: rDNA, SST1 array, centromere, and q-arm. Below, we show T2T-CHM13-relative homology mosaics for each chromosome 13 matched contig from HPRCy1-acro, with the most-similar reference chromosome at each region shown using the given colors (Target). (B) Aggregated untangle results in the SAACs. For each acrocentric chromosome, we show the count of its HPRCy1 q-arm-anchored contigs mapping itself and all other acrocentrics (Contigs), (C) as well as the regional (50kbp) untangle entropy metric (Regional homology entropy) computed over the contigs' T2T-CHM13-relative untanglings. (D) By considering the multiple untangling of each HPRCy1-acro contig, we develop a point-wise metric that captures diversity in T2T-CHM13-relative homology patterns (Positional homology entropy), leading to our definition of the PHRs. (E) The patterns of homology mosaicism suggest ongoing recombination exchange in the SAACs. A scan over T2T-CHM13 reveals that the rDNA units are enriched for PRDM9 binding motifs, and thus may host frequent double stranded breaks during meiosis. In (B-D) a gray background indicates regions with missing data due to the lack of non-T2T-CHM13 contigs. We provide the Centromeric Satellite Annotation (CenSat Annotation) track legend in Supplementary Figure 12.

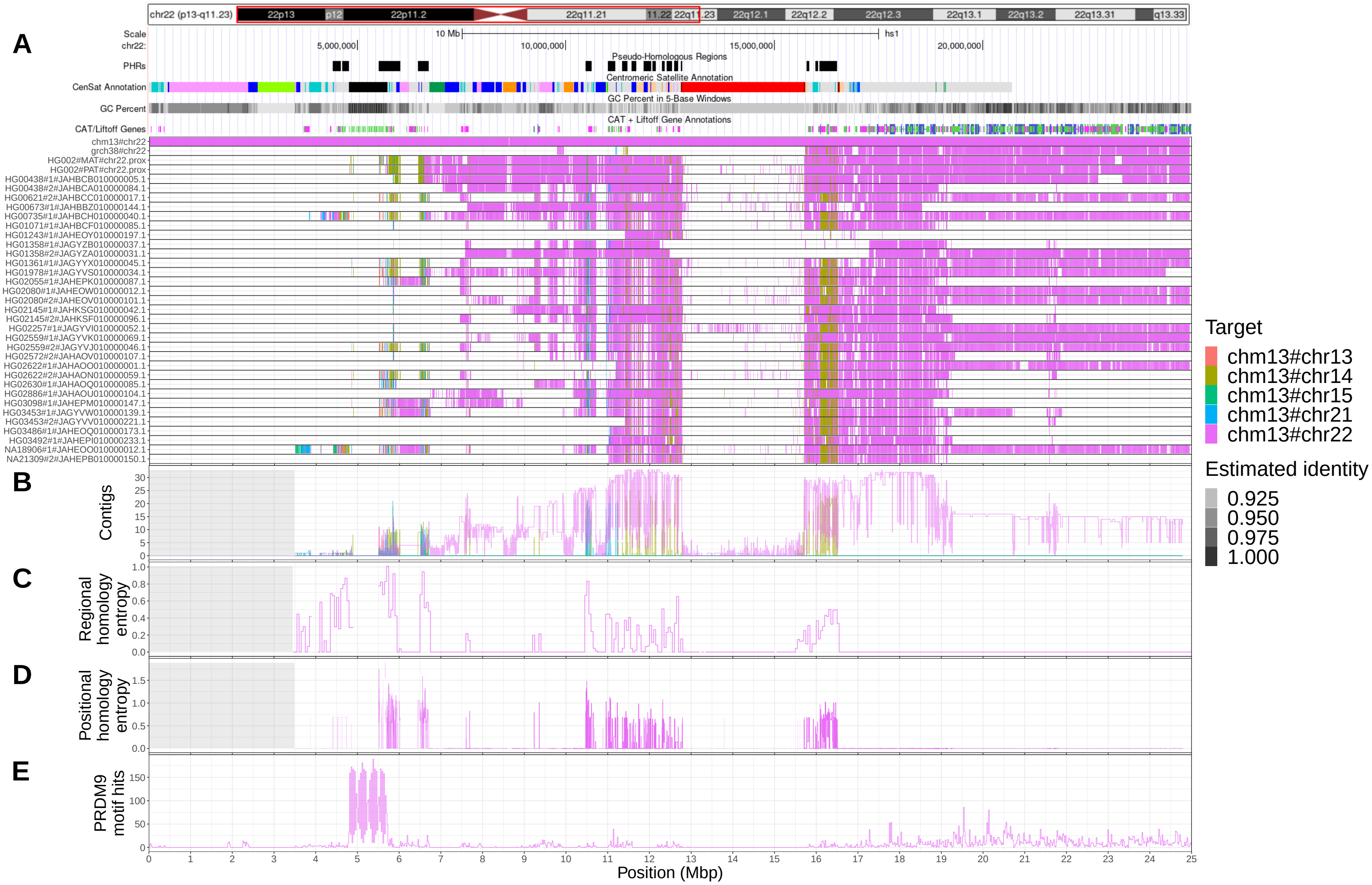


Supplementary Figure 17: (A) We focus on the first 25 Mbp of chromosome 22 shown here as a red box over T2T-CHM13 cytobands. Pseudo-homologous regions (PHRs), where diverse sets of acrocentric chromosomes recombine, are highlighted relative to T2T-CHM13 genome annotations for repeats, GC percentage, and genes. Above, we indicate regions of interest described in the main text: rDNA, SST1 array, centromere, and q-arm. Below, we show T2T-CHM13-relative homology mosaics for each chromosome 13 matched contig from HPRCy1-acro, with the most-similar reference chromosome at each region shown using the given colors (Target). (B) Aggregated untangle results in the SAACs. For each acrocentric chromosome, we show the count of its HPRCy1 q-arm-anchored contigs mapping itself and all other acrocentrics (Contigs), (C) as well as the regional (50kbp) untangle entropy metric (Regional homology entropy) computed over the contigs' T2T-CHM13-relative untanglings. (D) By considering the multiple untangling of each HPRCy1-acro contig, we develop a point-wise metric that captures diversity in T2T-CHM13-relative homology patterns (Positional homology entropy), leading to our definition of the PHRs. (E) The patterns of homology mosaicism suggest ongoing recombination exchange in the SAACs. A scan over T2T-CHM13 reveals that the rDNA units are enriched for PRDM9 binding motifs, and thus may host frequent double stranded breaks during meiosis. In (B-D) a gray background indicates regions with missing data due to the lack of non-T2T-CHM13 contigs. We provide the Centromeric Satellite Annotation (CenSat Annotation) track legend in Supplementary Figure 12.

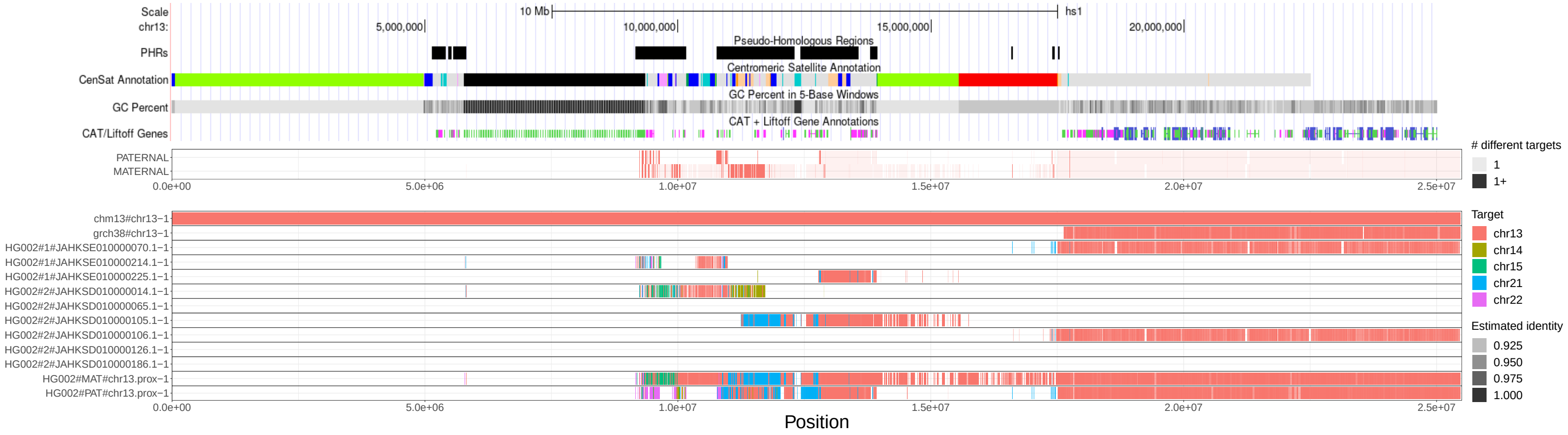


Supplementary Figure 18: For each base position of T2T-CHM13 chromosome 13, we compared the untangling best-hit of the HG002-HPRCy1 contigs with the best-hit supported by HG002-Verkko's contigs. We considered only best-hits with an estimated identity of at least 90% that cover regions labeled as reliable. We reported the number of different targets for each reference position. On the bottom, the untangling of HG002's contigs from HG002-Verkko assembly as those seen in HG002-HPRCy1 assemblies, for chromosome 13 versus T2T-CHM13. Transparency shows the estimated identity of the mappings. We display all mappings above 90% estimated pairwise identity. Checkerboard patterns observed in several regions of the SAACs correspond to contexts that may permit ongoing recombination.

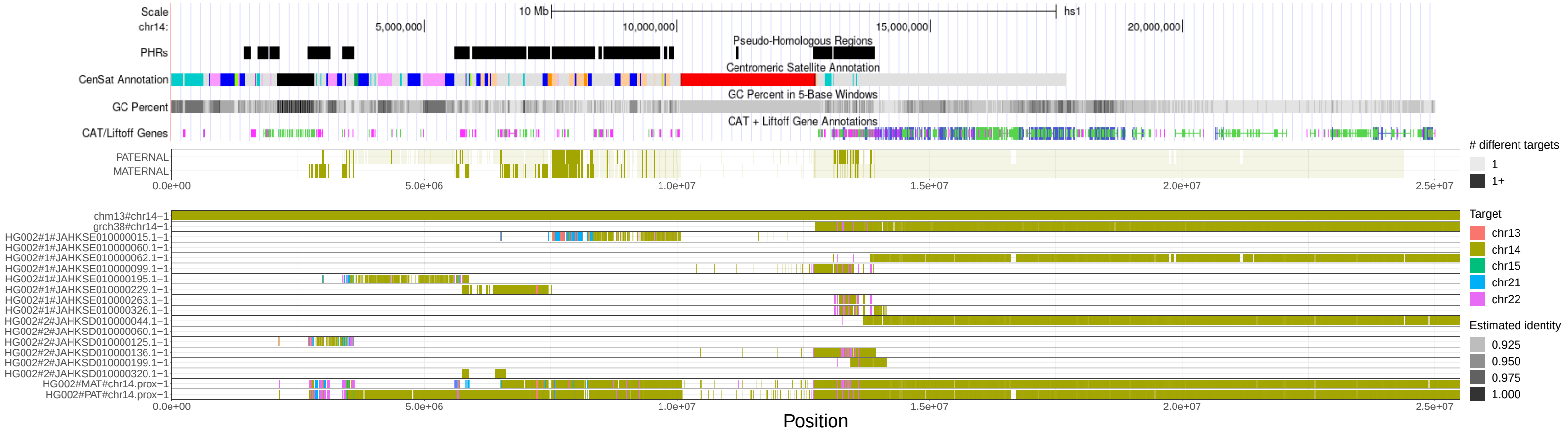


Supplementary Figure 19: For each base position of T2T-CHM13 chromosome 14, we compared the untangling best-hit of the HG002-HPRCy1 contigs with the best-hit supported by HG002-Verkko's contigs. We considered only best-hits with an estimated identity of at least 90% that cover regions labeled as reliable. We reported the number of different targets for each reference position. On the bottom, the untangling of HG002's contigs from HG002-Verkko assembly as those seen in HG002-HPRCy1 assemblies, for chromosome 14 versus T2T-CHM13. Transparency shows the estimated identity of the mappings. We display all mappings above 90% estimated pairwise identity. Checkerboard patterns observed in several regions of the SAACs correspond to contexts that may permit ongoing recombination.

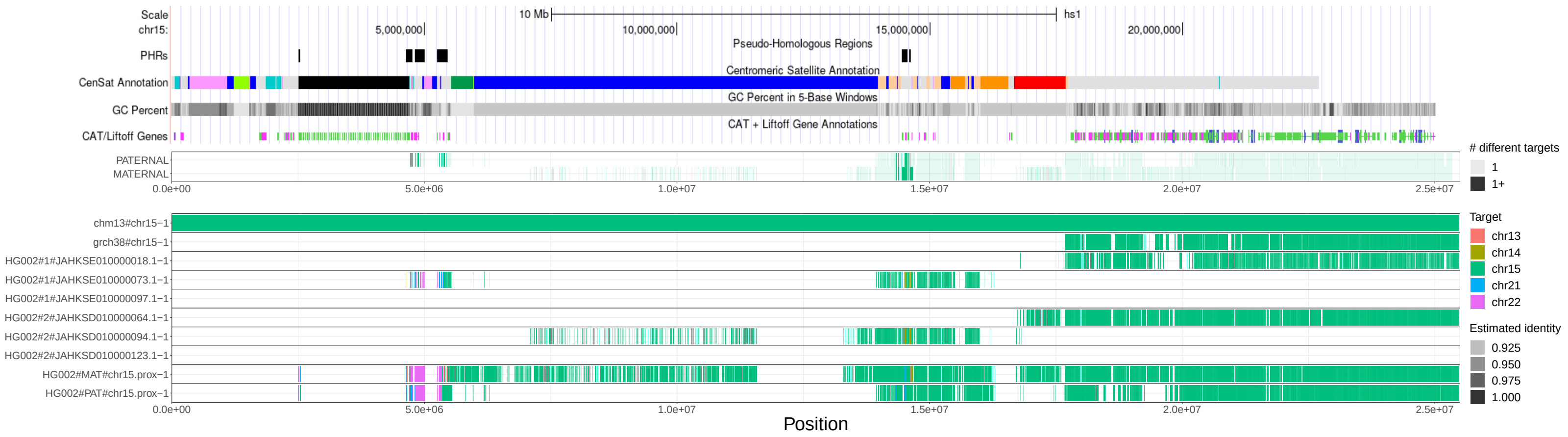


Supplementary Figure 20: For each base position of T2T-CHM13 chromosome 15, we compared the untangling best-hit of the HG002-HPRCy1 contigs with the best-hit supported by HG002-Verkko's contigs. We considered only best-hits with an estimated identity of at least 90% that cover regions labeled as reliable. We reported the number of different targets for each reference position. On the bottom, the untangling of HG002's contigs from HG002-Verkko assembly as those seen in HG002-HPRCy1 assemblies, for chromosome 15 versus T2T-CHM13. Transparency shows the estimated identity of the mappings. We display all mappings above 90% estimated pairwise identity. Checkerboard patterns observed in several regions of the SAACs correspond to contexts that may permit ongoing recombination.

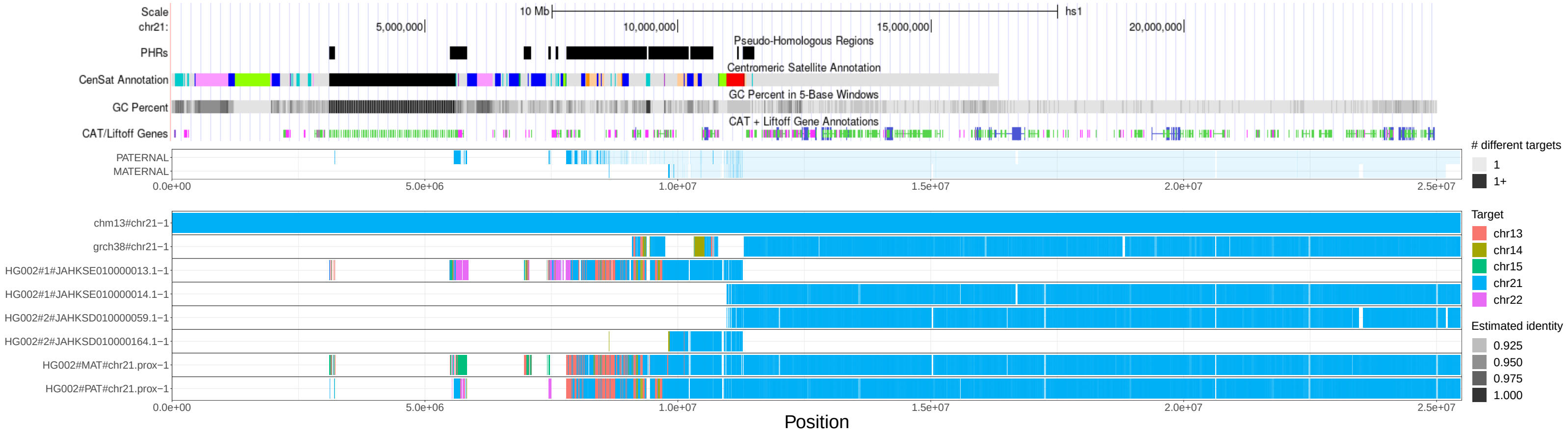


Supplementary Figure 21: For each base position of T2T-CHM13 chromosome 21, we compared the untangling best-hit of the HG002-HPRCy1 contigs with the best-hit supported by HG002-Verkko's contigs. We considered only best-hits with an estimated identity of at least 90% that cover regions labeled as reliable. We reported the number of different targets for each reference position. On the bottom, the untangling of HG002's contigs from HG002-Verkko assembly as those seen in HG002-HPRCy1 assemblies, for chromosome 21 versus T2T-CHM13. Transparency shows the estimated identity of the mappings. We display all mappings above 90% estimated pairwise identity. Checkerboard patterns observed in several regions of the SAACs correspond to contexts that may permit ongoing recombination.

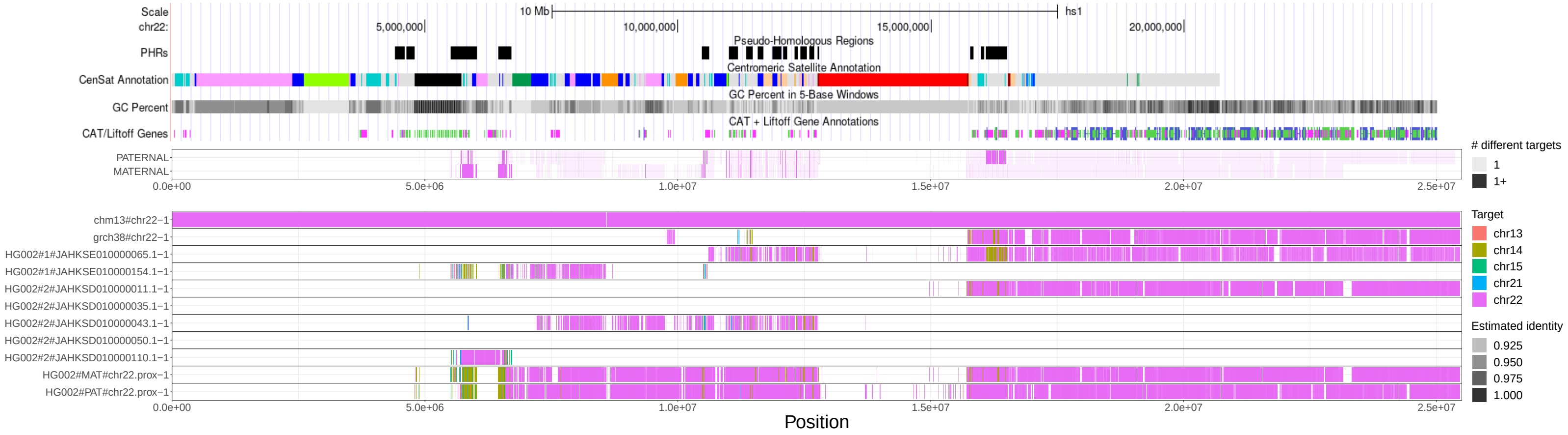


Supplementary Figure 22: For each base position of T2T-CHM13 chromosome 22, we compared the untangling best-hit of the HG002-HPRCy1 contigs with the best-hit supported by HG002-Verkko's contigs. We considered only best-hits with an estimated identity of at least 90% that cover regions labeled as reliable. We reported the number of different targets for each reference position. On the bottom, the untangling of HG002's contigs from HG002-Verkko assembly as those seen in HG002-HPRCy1 assemblies, for chromosome 22 versus T2T-CHM13. Transparency shows the estimated identity of the mappings. We display all mappings above 90% estimated pairwise identity. Checkerboard patterns observed in several regions of the SAACs correspond to contexts that may permit ongoing recombination.

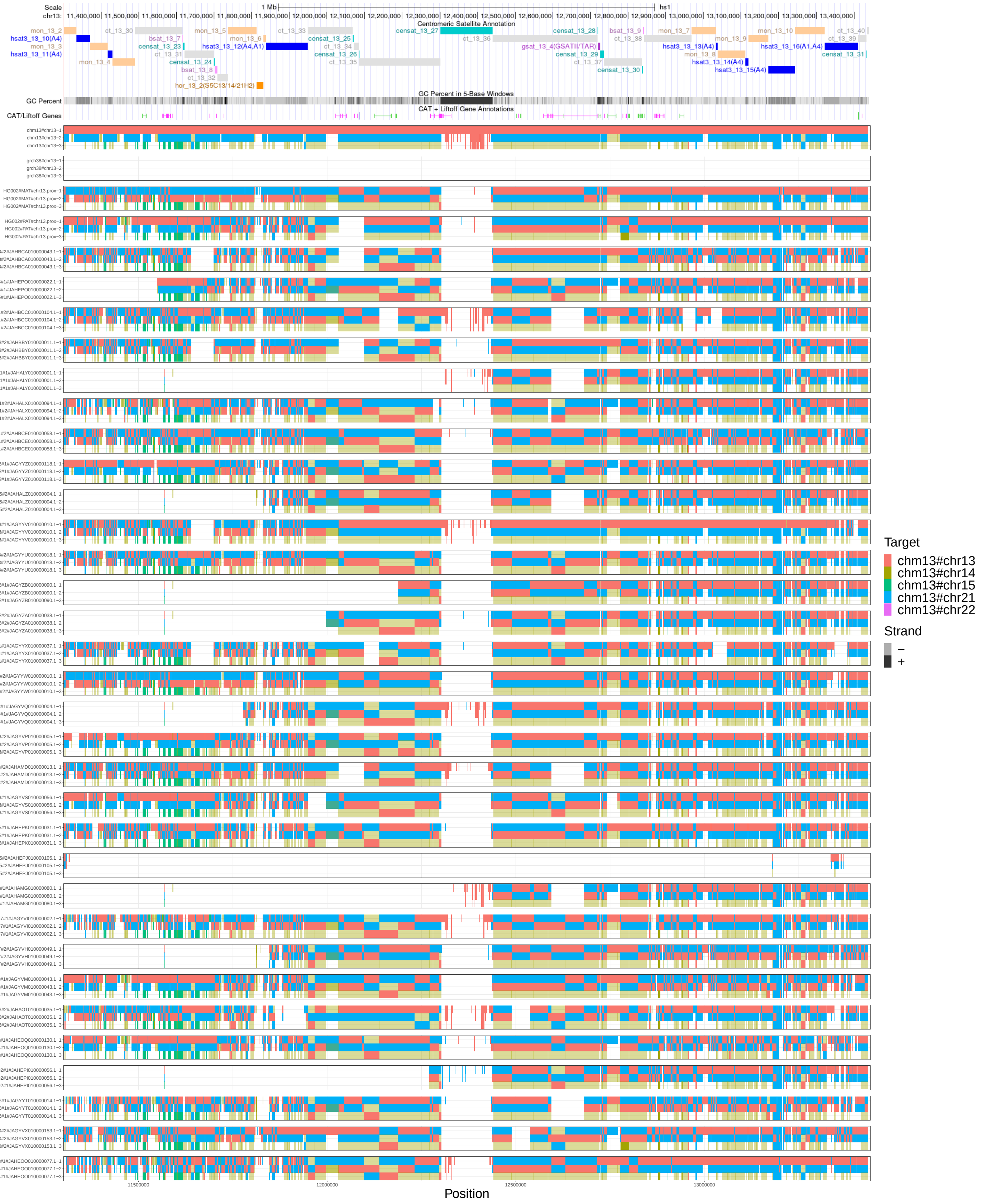


Supplementary Figure 23: Multiple untangling of T2T-CHM13, GRCh38, HG002-Verkko haplotypes, and HPRCy1-acro contigs versus T2T-CHM13. Chromosome 13 results are represented, in the chr13:11,301,367-13,440,010 region (censat_13_27 coordinates +- 1Mbp). Transparency shows the different orientations of the mappings. We display all mappings above 90% estimated pairwise identity. To analyze simultaneous hits to all acrocentrics, each grouping shows the first 3 best alternative mappings. The figure shows that chromosome 13's contigs map in forward orientation on T2T-CHM13 chromosome 13 and 21 (orange and cyan rectangles), while their mappings are inverted on chromosomes 14 (transparent gold rectangles).

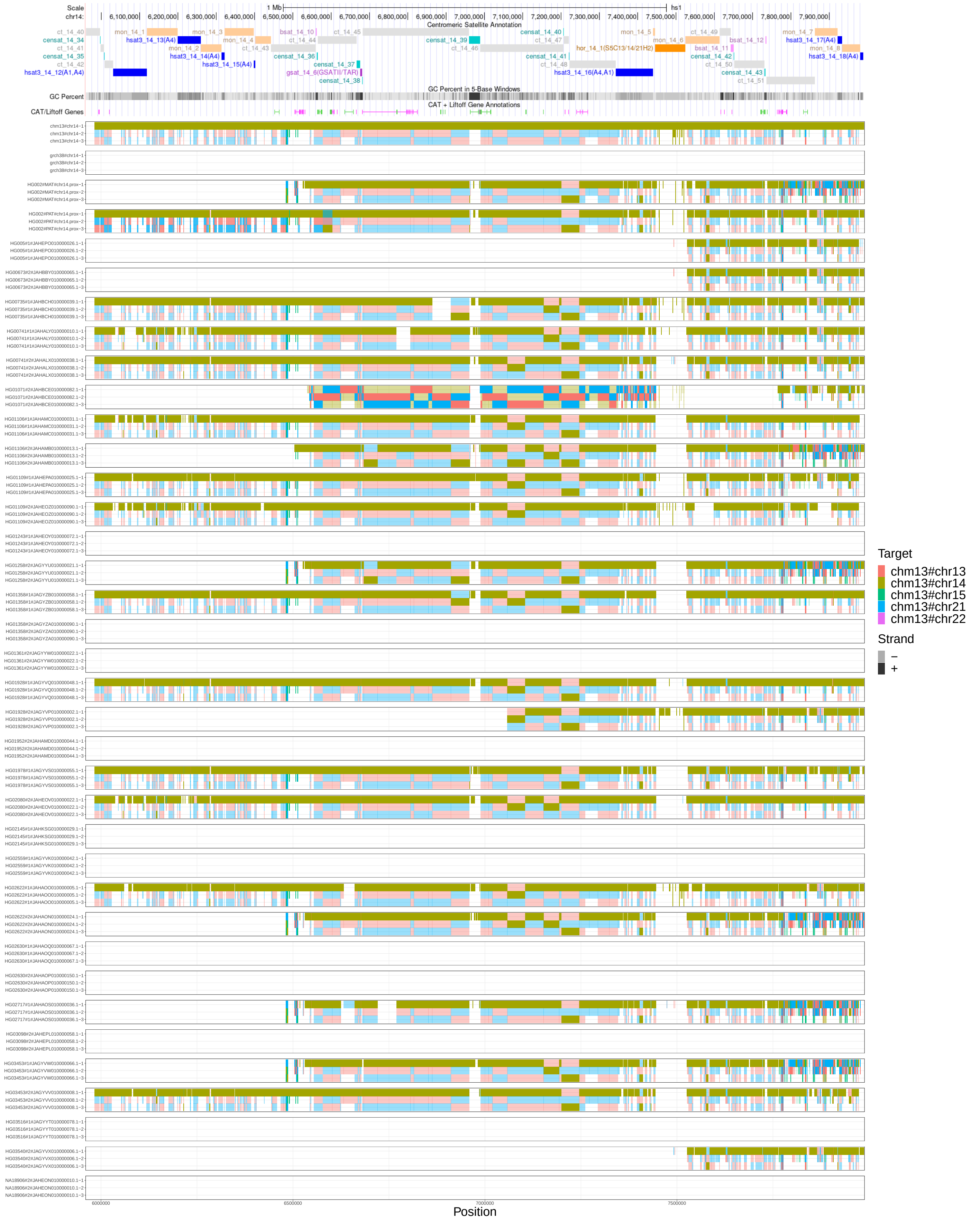


Supplementary Figure 24: Multiple untangling of T2T-CHM13, GRCh38, HG002-Verkko haplotypes, and HPRCy1-acro contigs versus T2T-CHM13. Chromosome 14 results are represented, in the chr14:5,960,008-7,988,409 region (censat_14_39 coordinates +- 1Mbp). Transparency shows the different orientations of the mappings. We display all mappings above 90% estimated pairwise identity. To analyze simultaneous hits to all acrocentrics, each grouping shows the first 3 best alternative mappings. The figure shows that chromosome 14's contigs map in forward orientation on T2T-CHM13 chromosome 14 (gold rectangles), while their mappings are inverted on chromosomes 13 and 21 (transparent orange and cyan rectangles), with the sole exception of HG01071#2#JAHBCE010000082.1, where the trend is reversed.

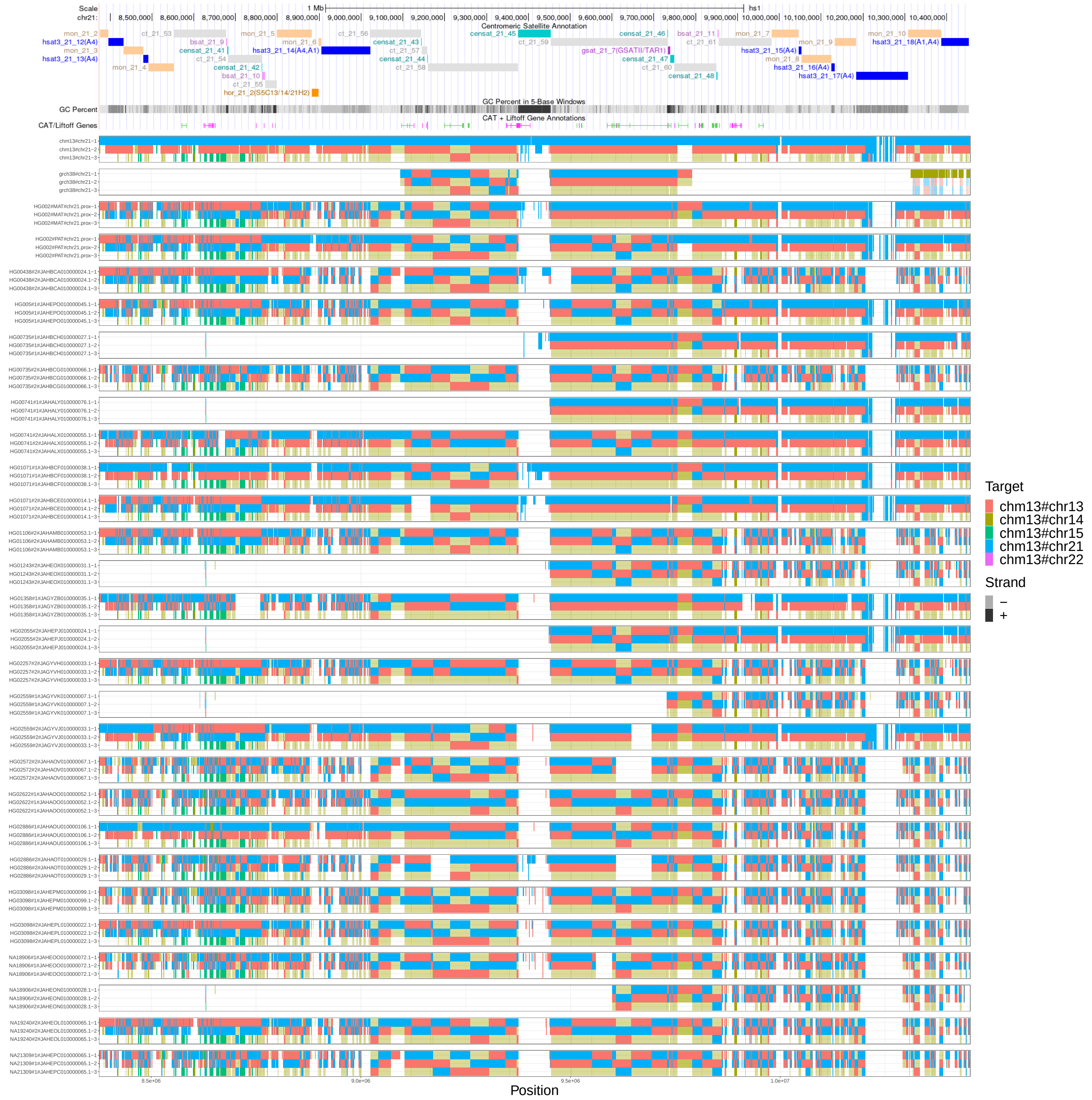


Supplementary Figure 25: Multiple untangling of T2T-CHM13, GRCh38, HG002-Verkko haplotypes, and HPRCy1-acro contigs versus T2T-CHM13. Chromosome 21 results are represented, in the chr21:8,375,567-10,453,313 region (censat_21_45 coordinates +- 1Mbp). Transparency shows the different orientations of the mappings. We display all mappings above 90% estimated pairwise identity. To analyze simultaneous hits to all acrocentrics, each grouping shows the first 3 best alternative mappings. The figure shows that chromosome 21's contigs map in forward orientation on T2T-CHM13 chromosome 21 and 13 (cyan and orange rectangles), while their mappings are inverted on chromosomes 14 (transparent gold rectangles).

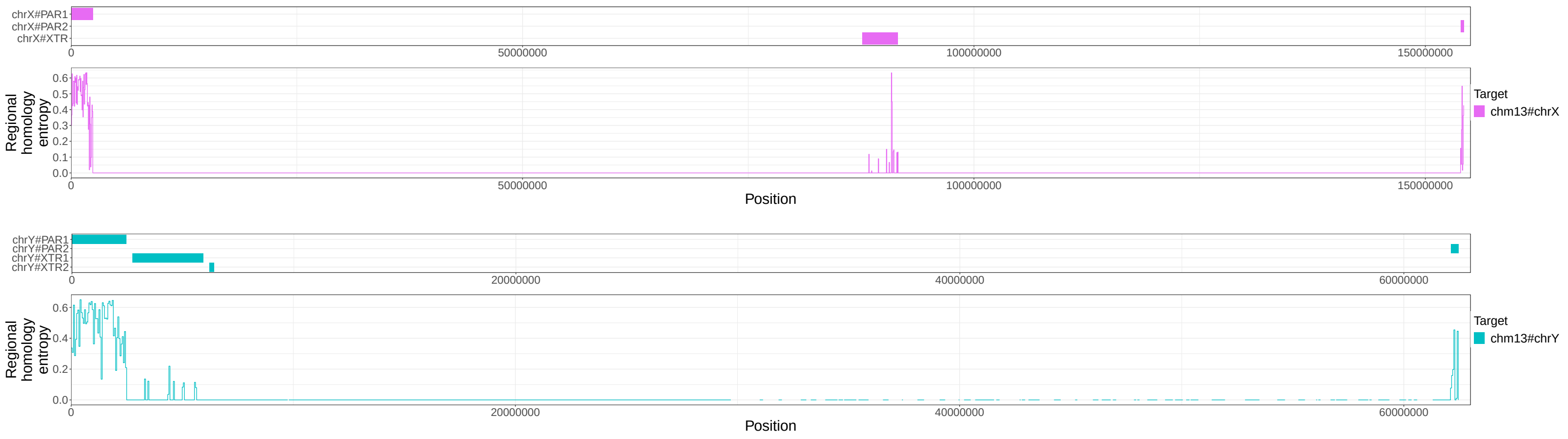


Supplementary Figure 26: For HPRCy1 contigs from chromosome X and Y, we show the tracks for the pseudoautosomal regions (PARs) and the X-transposed region (XTRs) with respect to T2T-CHM13 (on the top) as well as the untangle entropy metric (Regional homology metric, on the bottom) computed over the contigs' T2T-CHM13-relative untanglings.

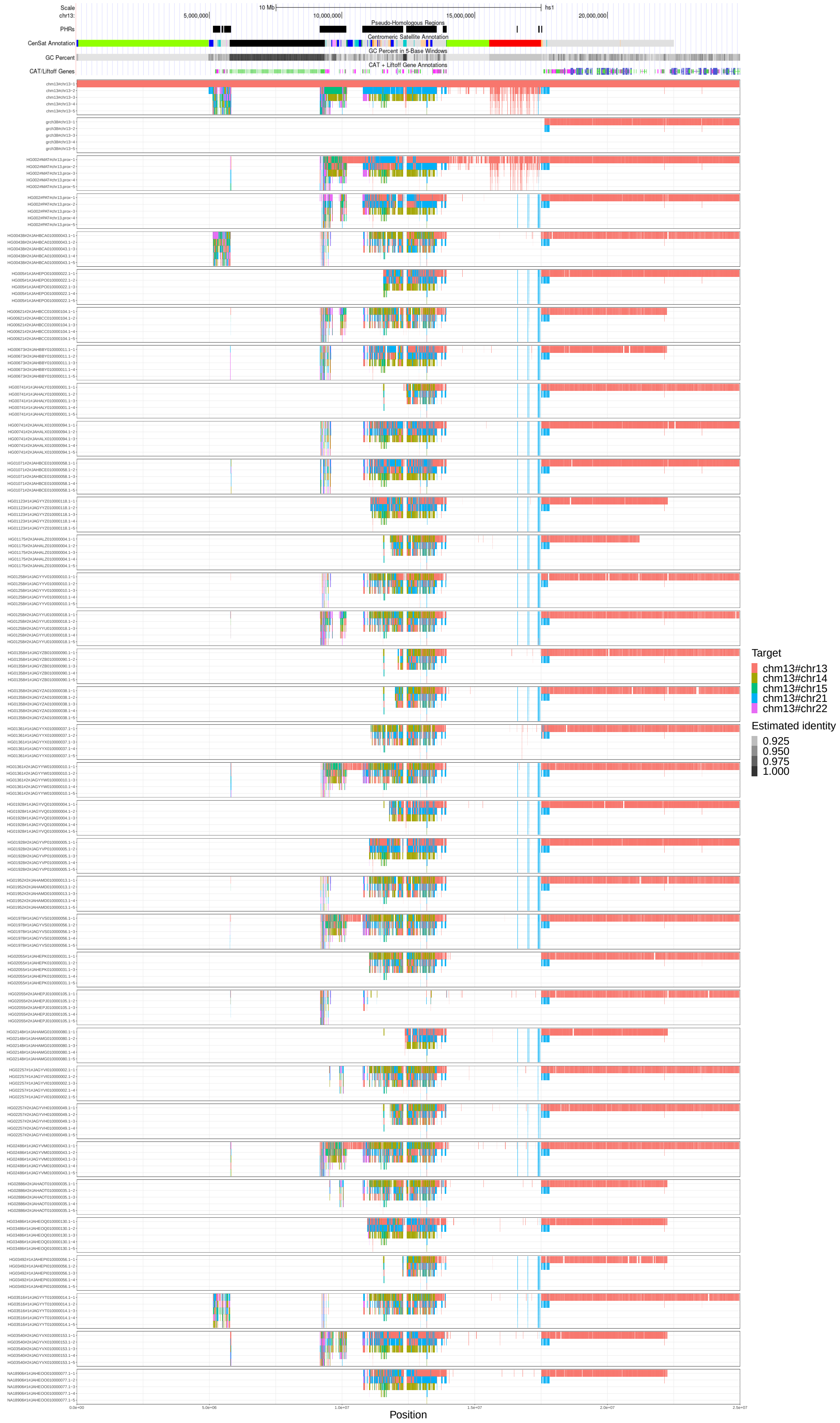


Supplementary Figure 27: Multiple untangling of HPRCy1-acro's sequences belonging to chromosome 13 versus T2T-CHM13. Transparency shows the estimated identity of the mappings. We display all mappings above 90% estimated pairwise identity that cover regions labeled as reliable. To allow the display of simultaneous hits to all acrocentrics, each grouping shows the first 5 best alternative mappings. Checkerboard patterns observed in several regions of the SAACs correspond to contexts that may permit ongoing recombination.

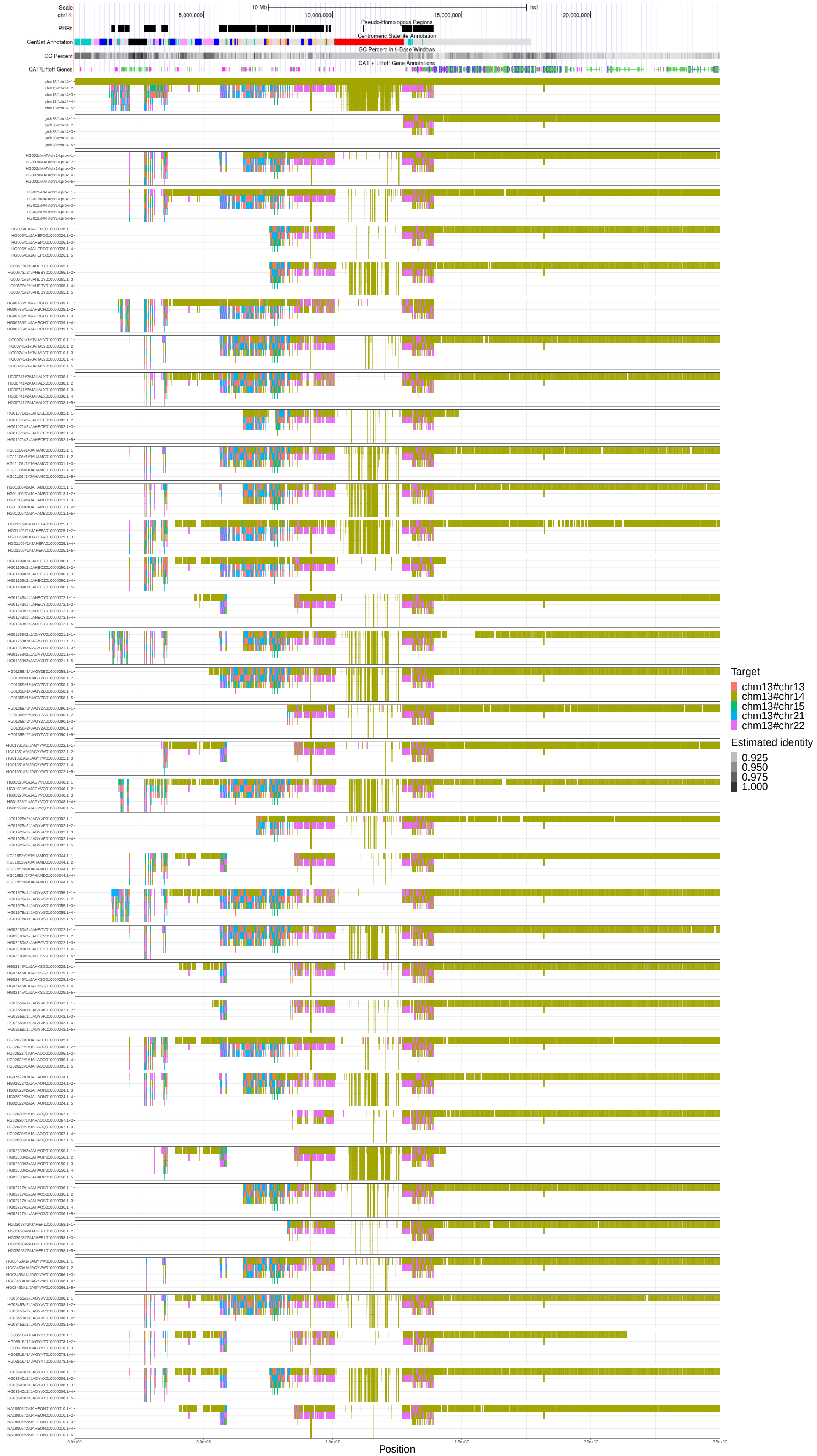


Supplementary Figure 28: Multiple untangling of HPRCy1-acro's sequences belonging to chromosome 14 versus T2T-CHM13. Transparency shows the estimated identity of the mappings. We display all mappings above 90% estimated pairwise identity that cover regions labeled as reliable. To allow the display of simultaneous hits to all acrocentrics, each grouping shows the first 5 best alternative mappings. Checkerboard patterns observed in several regions of the SAACs correspond to contexts that may permit ongoing recombination.

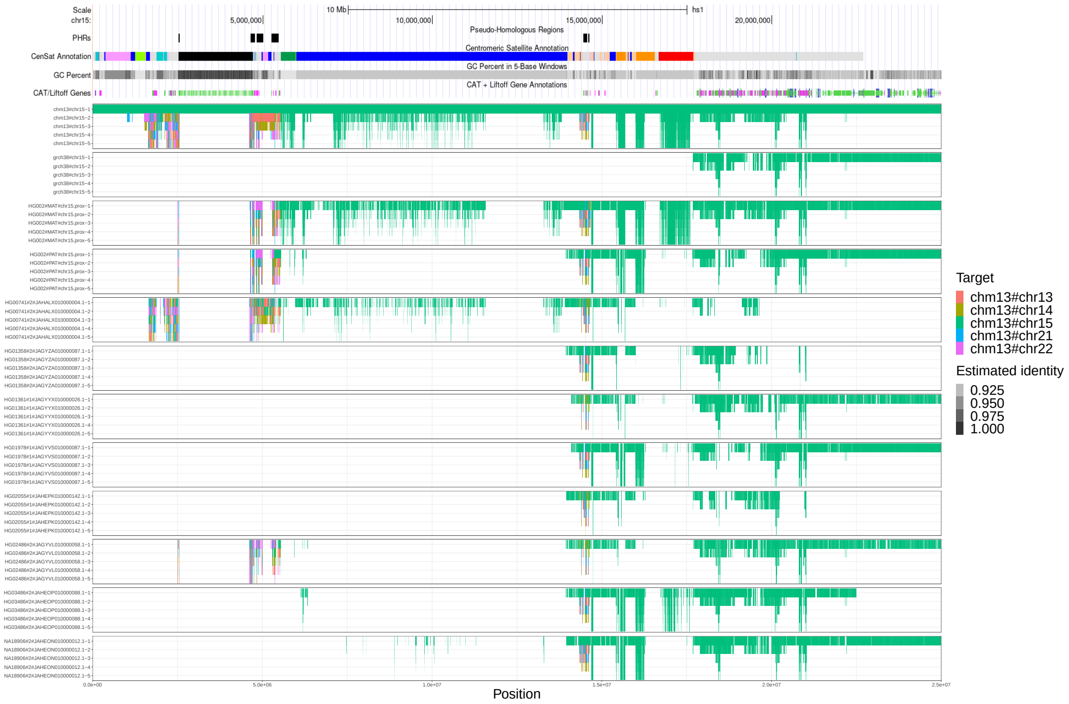


Supplementary Figure 29: Multiple untangling of HPRCy1-acro's sequences belonging to chromosome 15 versus T2T-CHM13. Transparency shows the estimated identity of the mappings. We display all mappings above 90% estimated pairwise identity that cover regions labeled as reliable. To allow the display of simultaneous hits to all acrocentrics, each grouping shows the first 5 best alternative mappings. Checkerboard patterns observed in several regions of the SAACs correspond to contexts that may permit ongoing recombination.

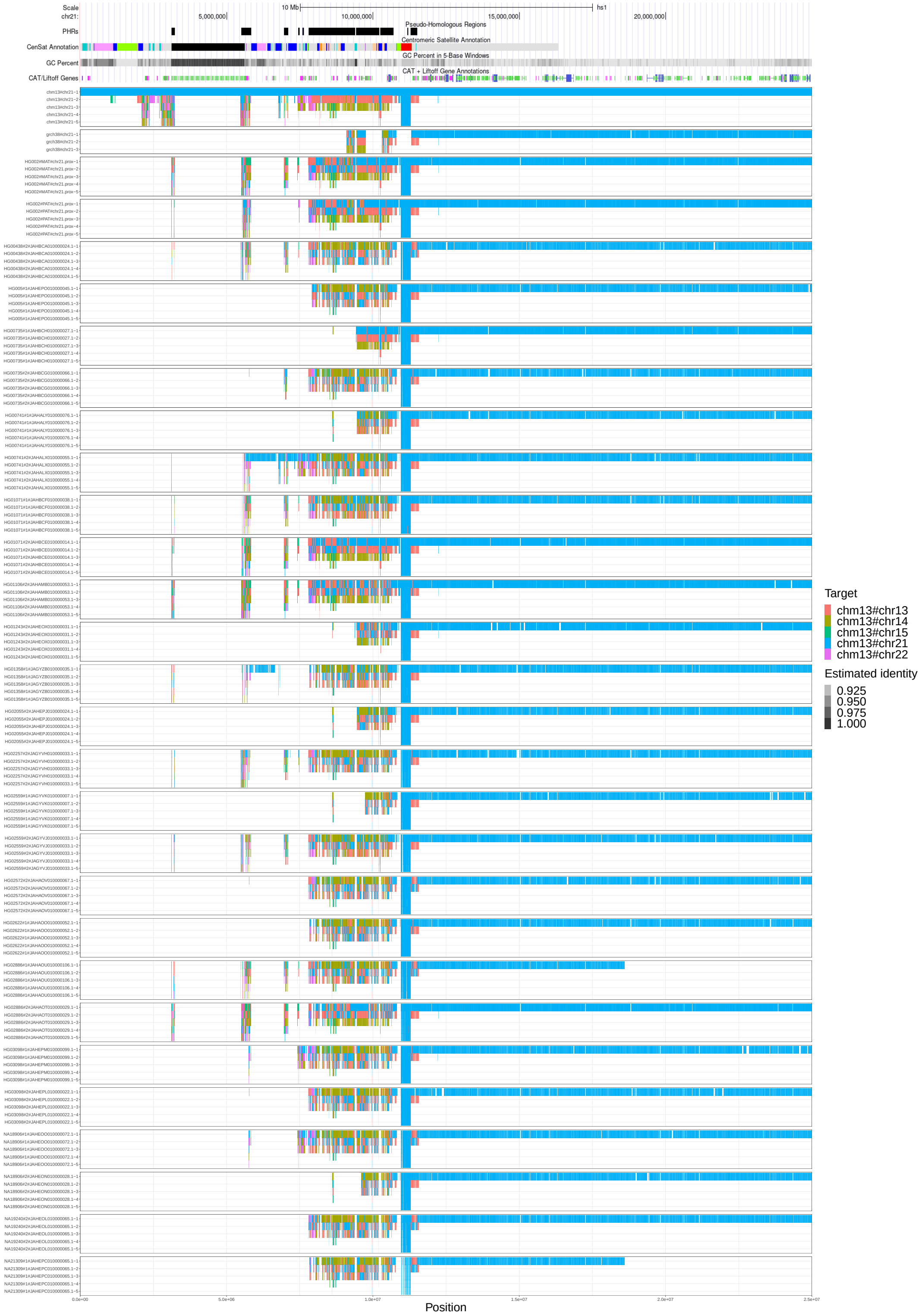


Supplementary Figure 30: Multiple untangling of HPRCy1-acro's sequences belonging to chromosome 21 versus T2T-CHM13. Transparency shows the estimated identity of the mappings. We display all mappings above 90% estimated pairwise identity that cover regions labeled as reliable. To allow the display of simultaneous hits to all acrocentrics, each grouping shows the first 5 best alternative mappings. Checkerboard patterns observed in several regions of the SAACs correspond to contexts that may permit ongoing recombination.

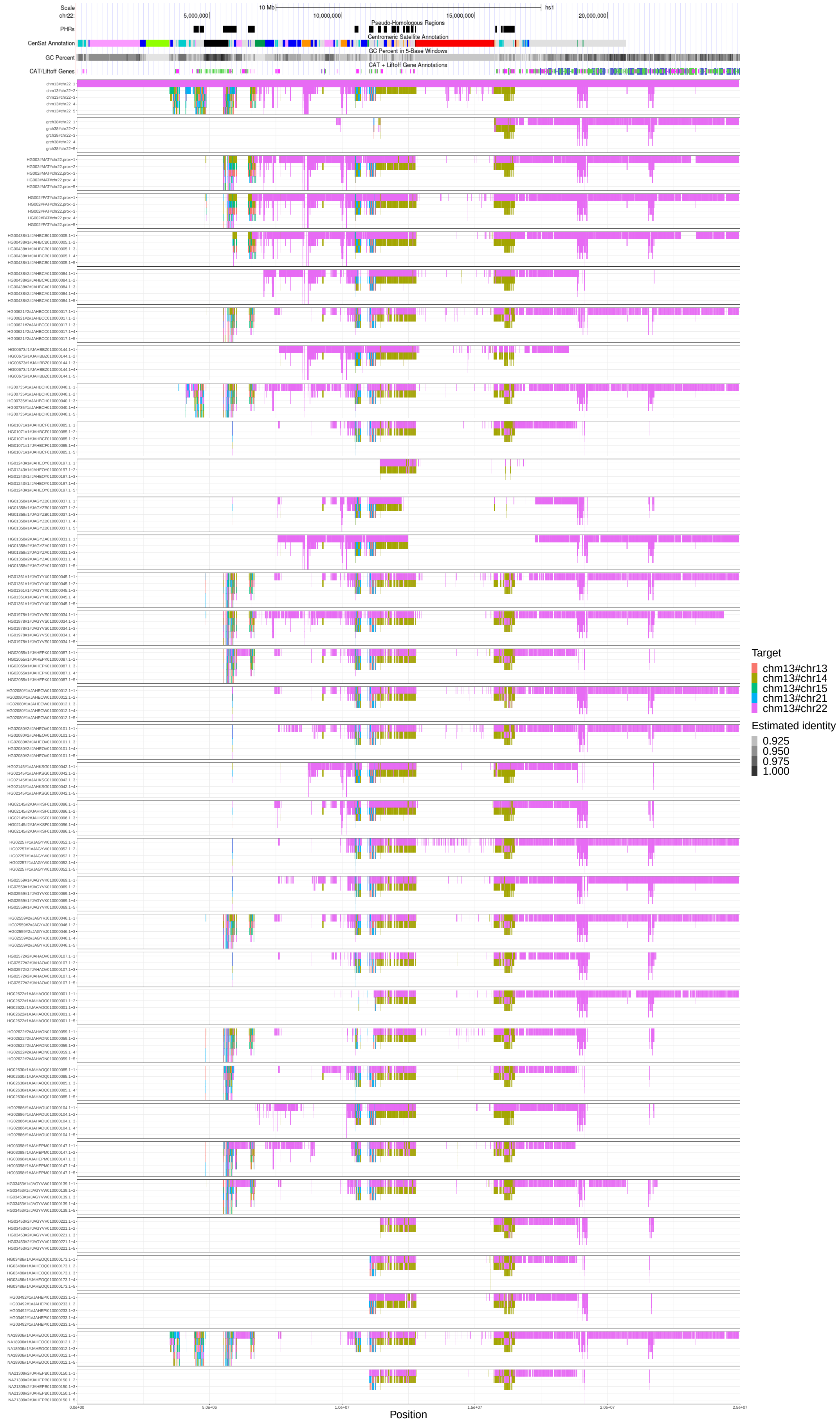


Supplementary Figure 31: Multiple untangling of HPRCy1-acro's sequences belonging to chromosome 22 versus T2T-CHM13. Transparency shows the estimated identity of the mappings. We display all mappings above 90% estimated pairwise identity that cover regions labeled as reliable. To allow the display of simultaneous hits to all acrocentrics, each grouping shows the first 5 best alternative mappings. Checkerboard patterns observed in several regions of the SAACs correspond to contexts that may permit ongoing recombination.

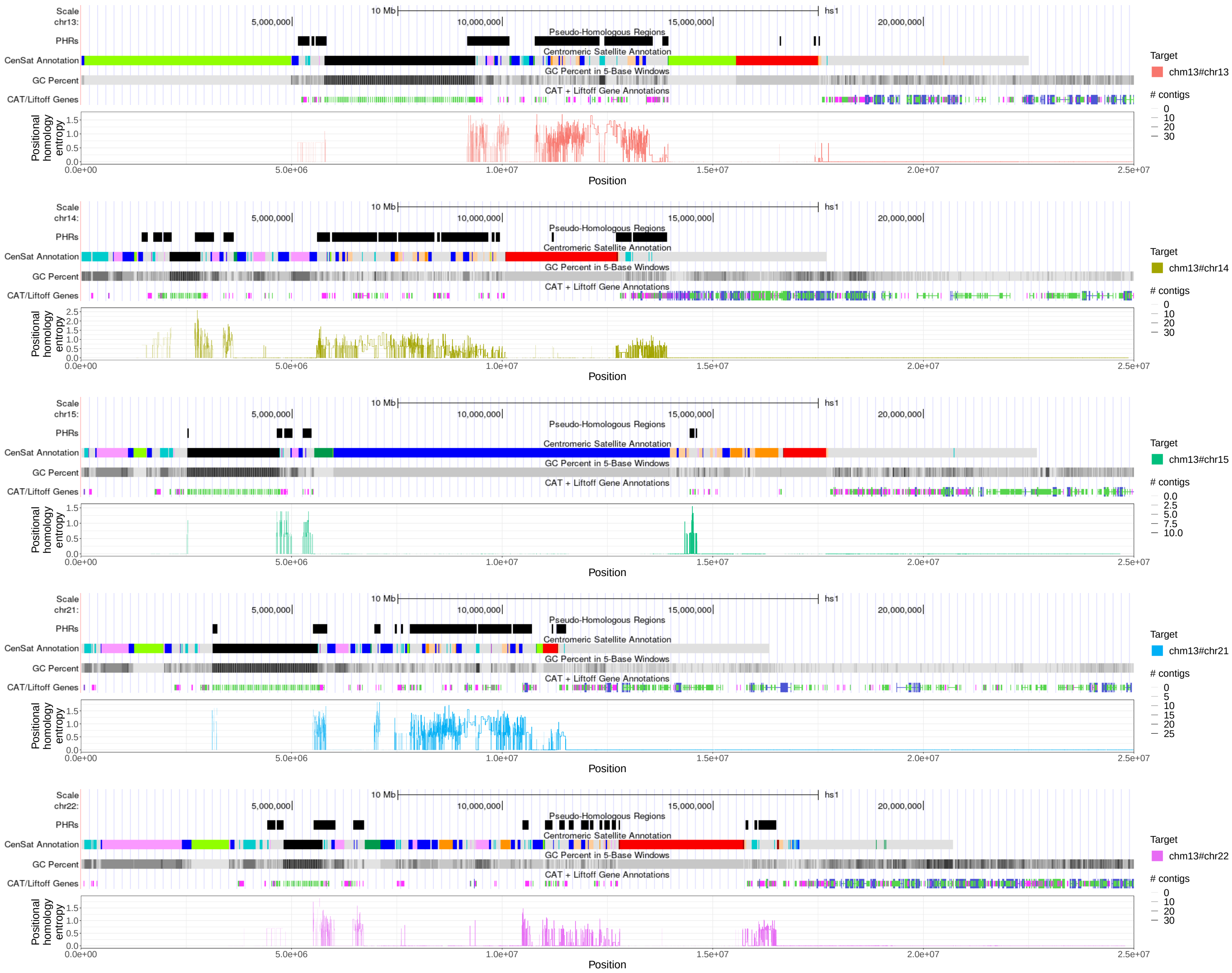


Supplementary Figure 32: Multi-hit untangling diversity entropy for HPRCy1-acro's sequences belonging to chromosome 13, 14, 15, 21, 22 versus T2T-CHM13. We considered all mappings above 90% estimated pairwise identity that cover regions labeled as reliable.

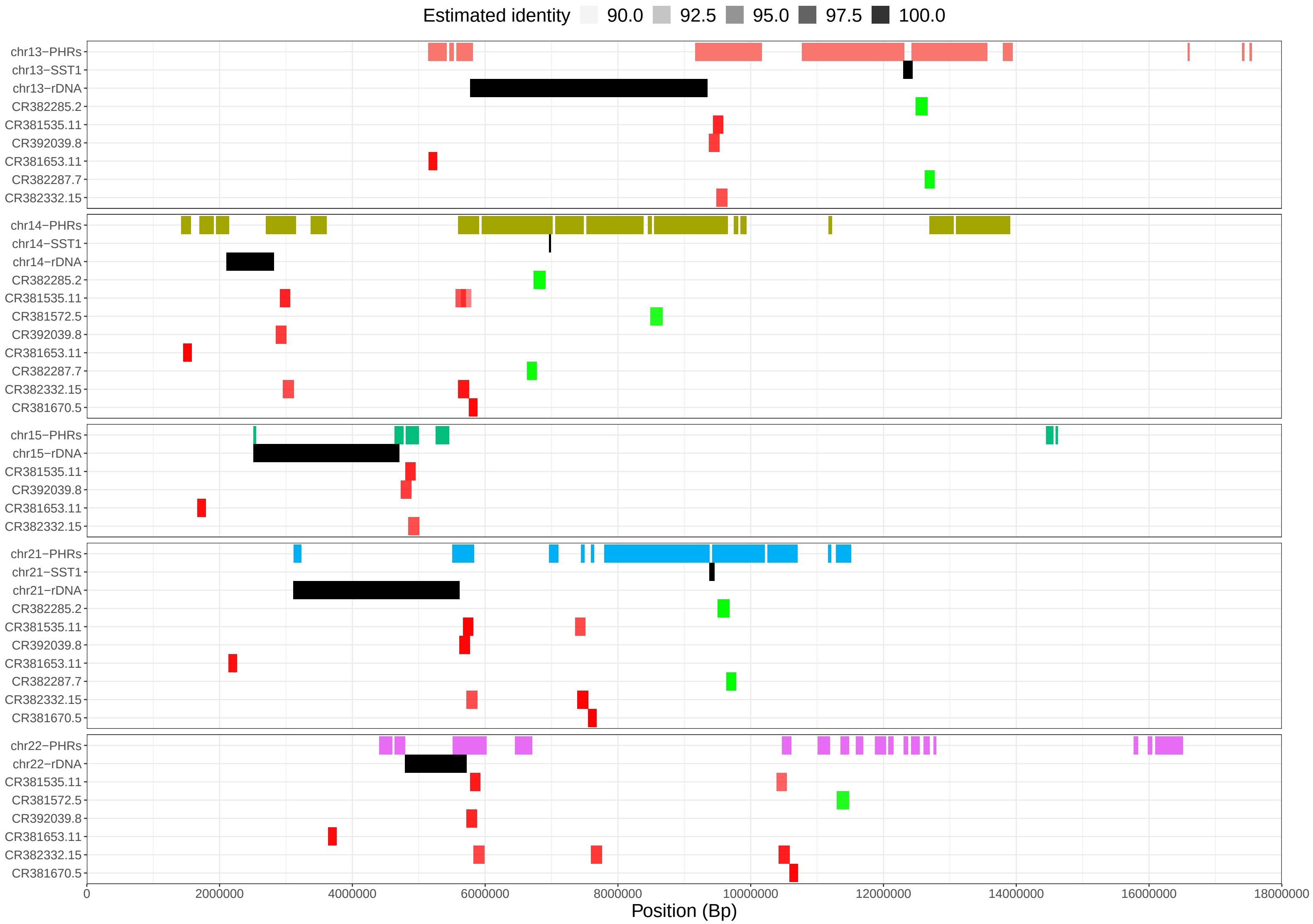


Supplementary Figure 33: For each T2T-CHM13 acrocentric chromosome, we show the tracks for the pseudo-homologous regions (PHRs), the SST1 array, the rDNA arrays, and for the regions where the bacterial artificial chromosome (BAC) clones from (Jarmuz-Szymczak et al. 2014) map on those chromosomes. Most of the mappings cover the PHRs. The mappings with higher estimated identity (¿ 99%) are on chr21 and chr14. We colored BAC clones' mappings according to (Jarmuz-Szymczak et al. 2014).

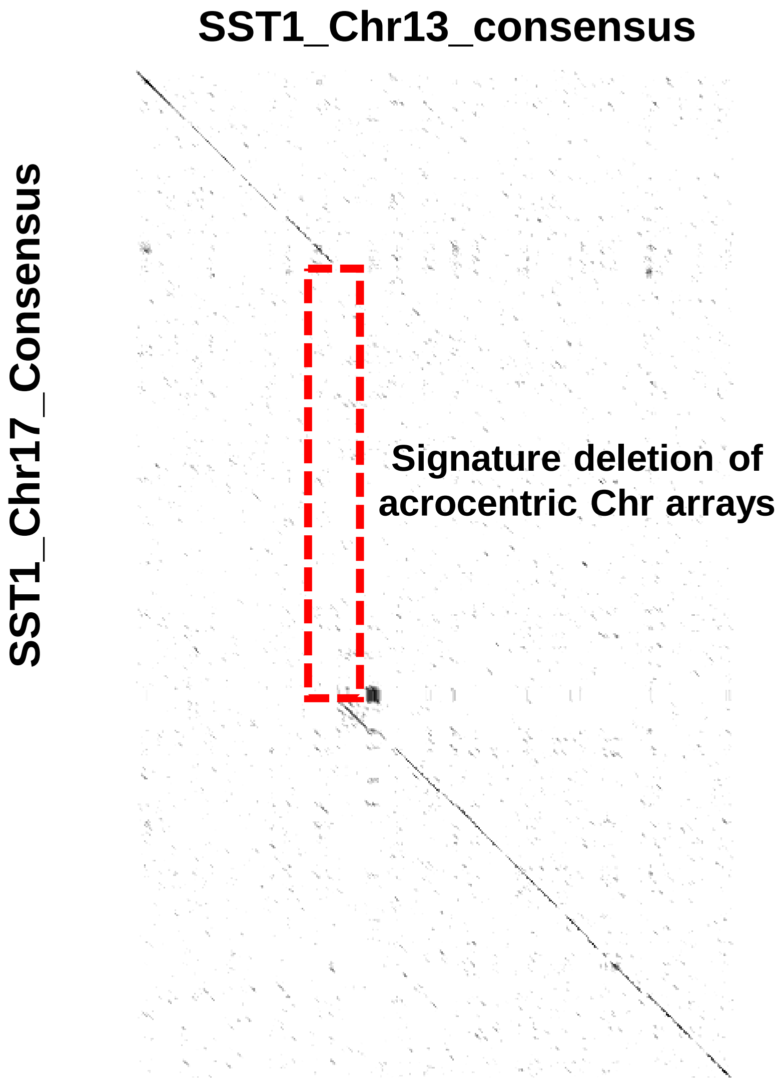


Supplementary Figure 34: Comparison of chr13 and chr17 SST1 consensus sequences by alignment dotplot indicate the unique deletion shared exclusively by the acrocentric chromosomes (dotted red rectangle).

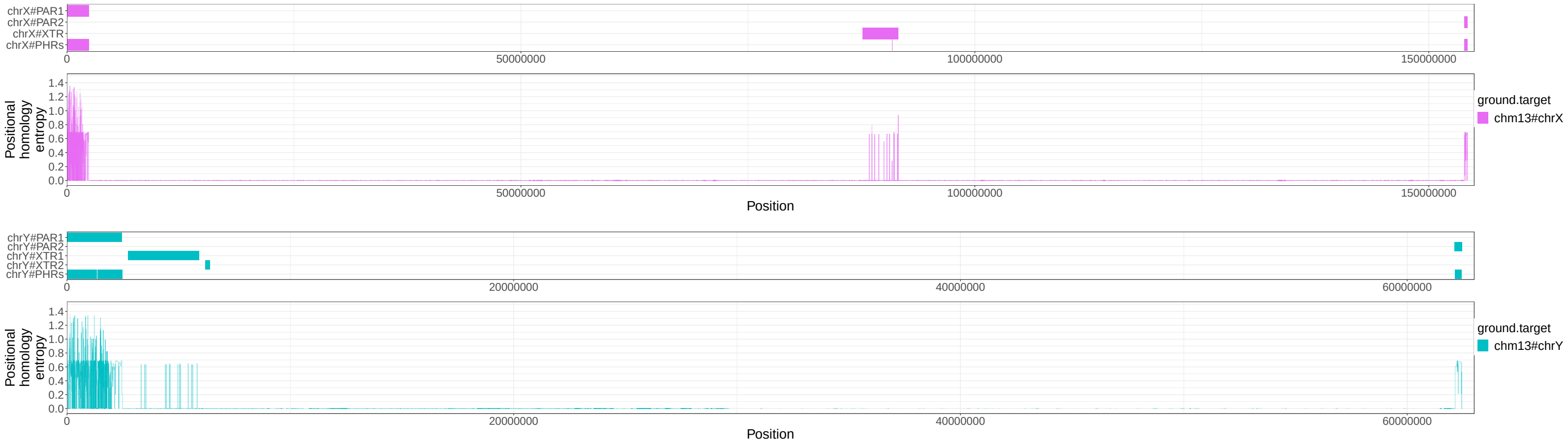


Supplementary Figure 35: For HPRCy1 contigs from chromosome X and Y, we show the tracks for the pseudoautosomal regions (PARs) and the X-transposed region (XTRs) with respect to T2T-CHM13 as well as the multi untangle entropy metric (Positional homology entropy, on the bottom) computed over the contigs' T2T-CHM13-relative untanglings.

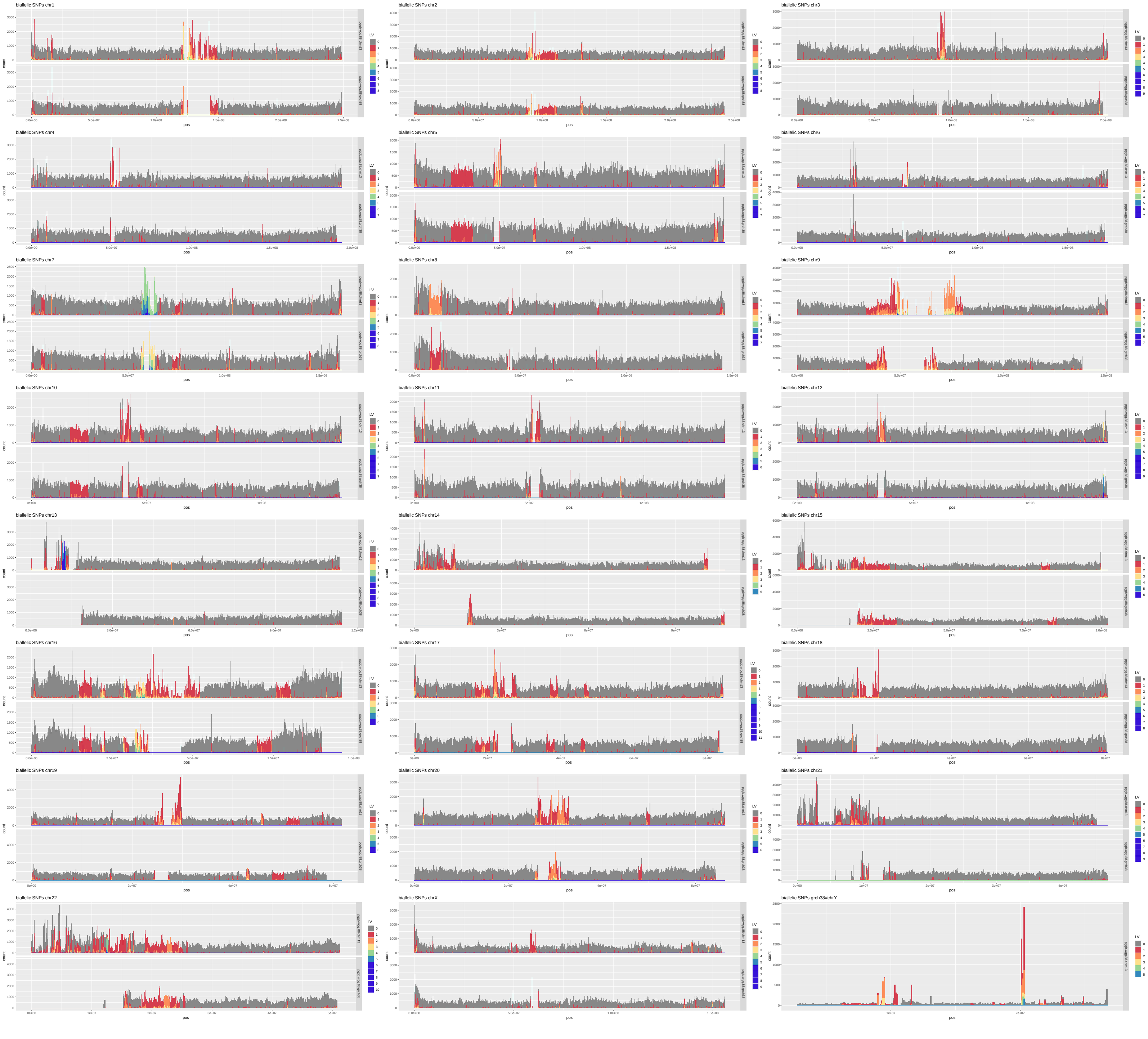

Supplementary Figure 36: Biallelic SNP density in the full HPRCy1 draft pangenome. Colors indicate the count of SNPs in each bin at given variant nesting levels (Paten et al. 2018), with gray showing the base level and a red to violet spectrum for increasing level.

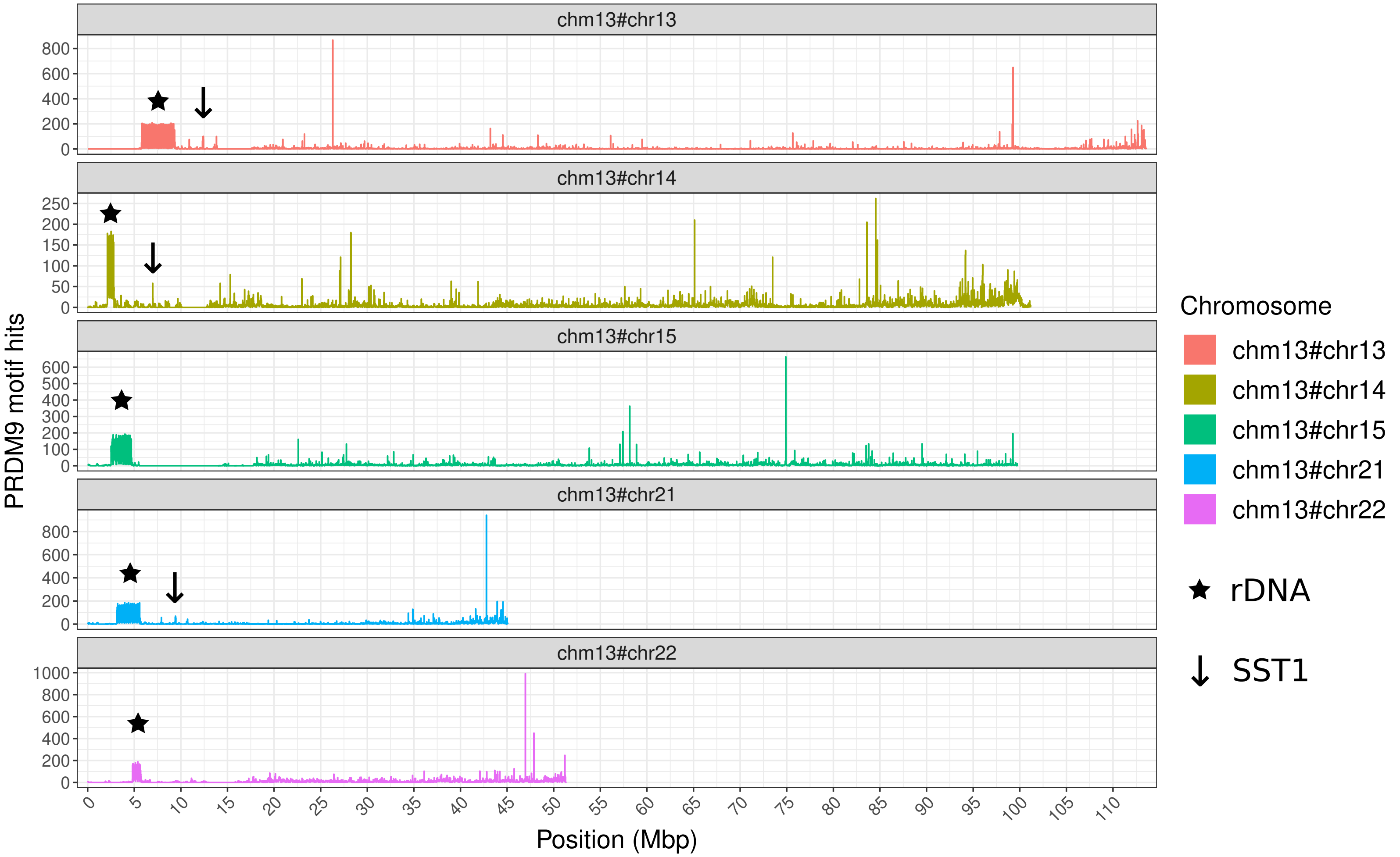


Supplementary Figure 37: For each T2T-CHM13 acrocentric chromosome, we show the number of human PRDM9 binding motif hits present in windows 20 kbps long.

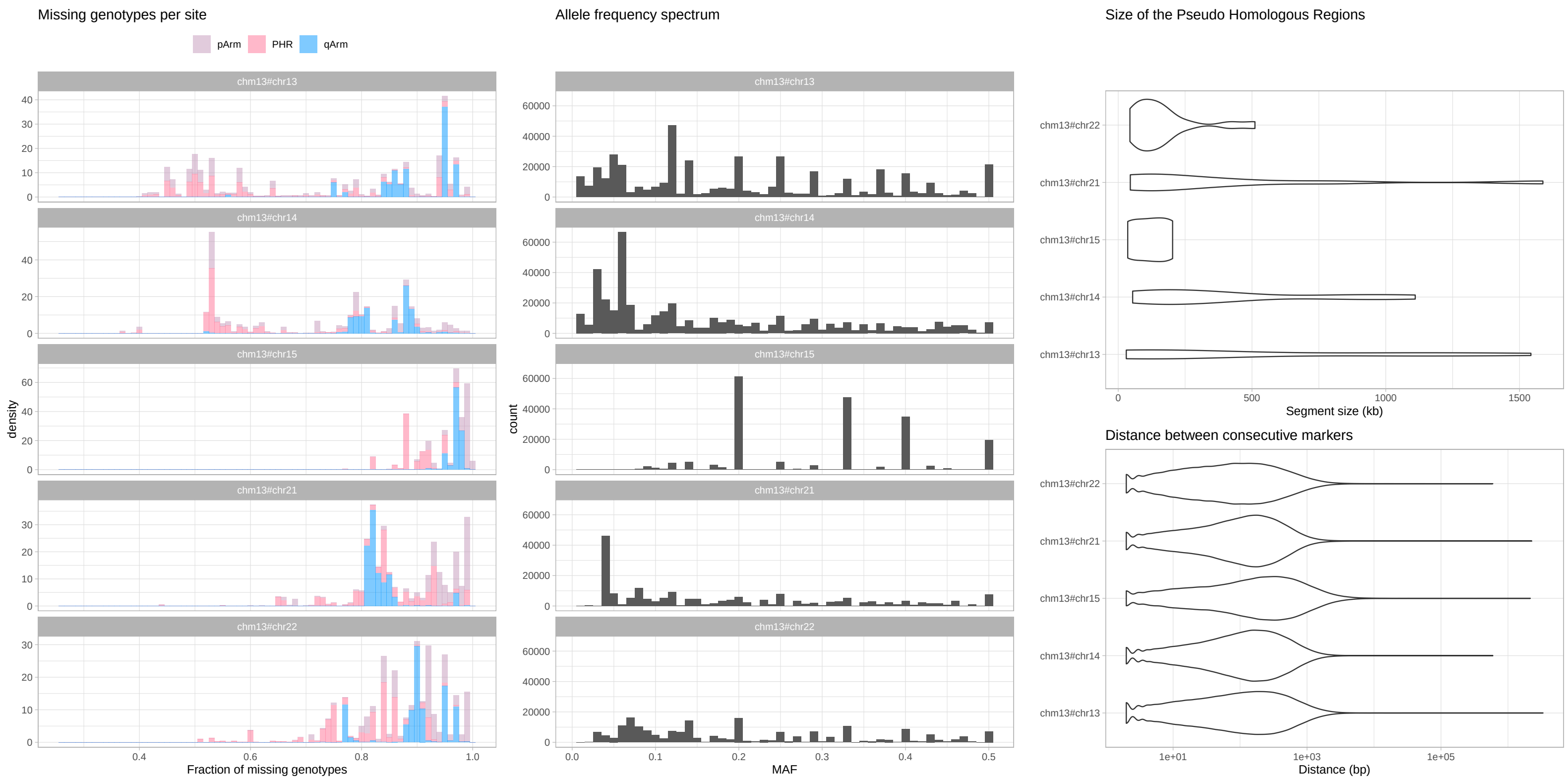


Supplementary Figure 38: Left) Histograms showing missing genotypes per site for variants called with respect to the T2T-CHM13 acrocentrics. Middle) Histograms showing the allele frequency spectrum for variants called with respect to the T2T-CHM13 acrocentrics. MAF = Minor Allele Frequency. Right-Top) Segment size distributions of the pseudo-homologous regions with respect to the T2T-CHM13 acrocentrics. Right-Bottom) Distributions of the distance between consecutive markers with respect to the T2T-CHM13 acrocentrics.

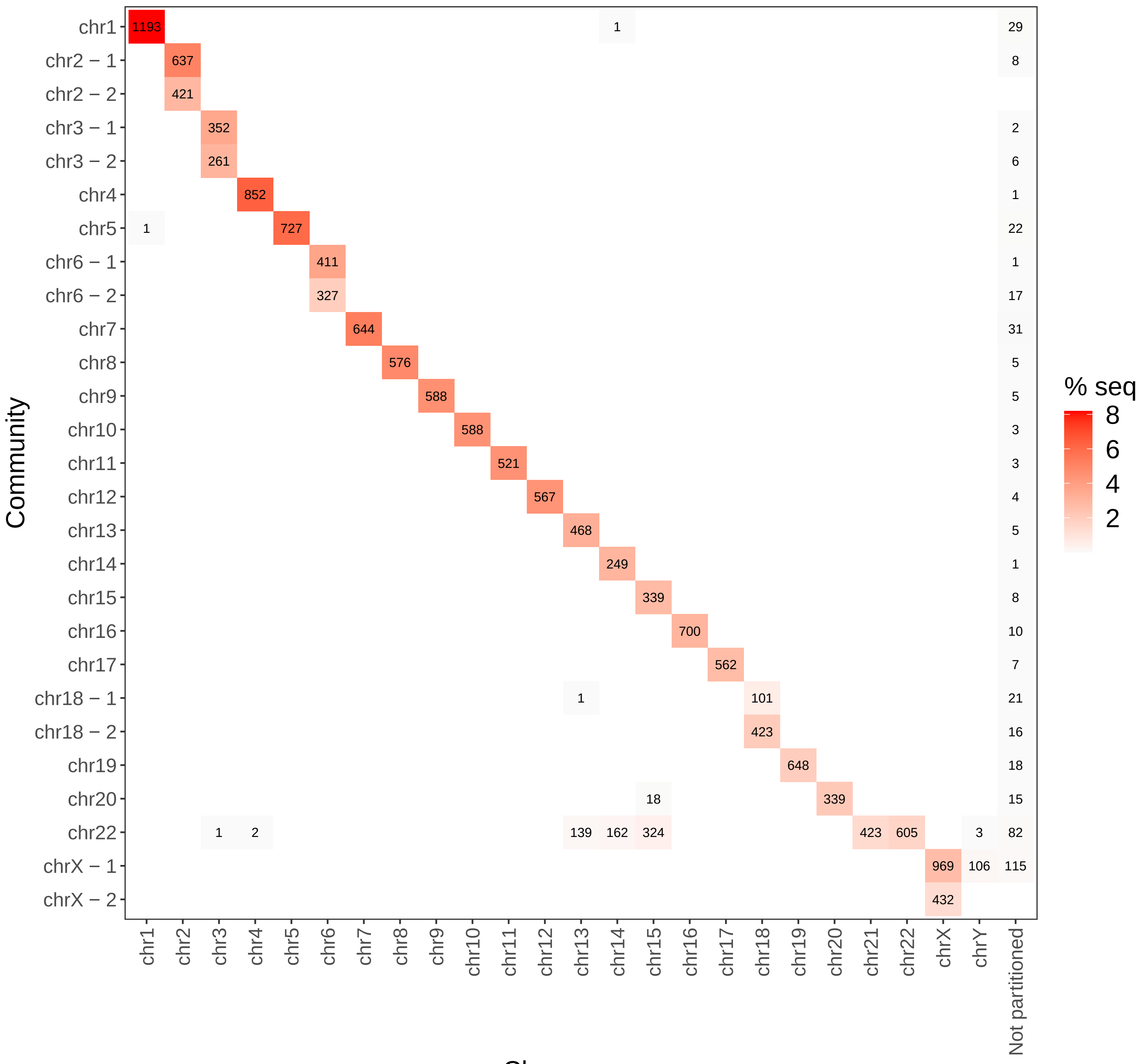


Supplementary Figure 39: Results of community assignment on the mapping graph when using chains of 10 kbp seeds of 95% average nucleotide identity. On the x-axis the chromosome to which contigs belong based on competitive mapping to T2T-CHM13 and GRCh38, while the y-axis indicates the community, which is named by the chromosome partition that contributes the largest number of contigs to it.
