## Supplementary Note 1 for "Recombination between heterologous human acrocentric chromosomes"

### 1 Physical coincidence of chromosomes on a manifold

We consider the situation of chromosomes in the space of the nucleus versus their coincidence near the nucleolar complex. We model how much more frequently should chromosomes occur at the same place if they are constrained to be near a 2D manifold versus freely moving in 3D. This provides intuition for the degree to which chromosomes that are maintained physically close to the nucleolus can physically interact with each other.

### 2 Problem formulation

Consider two types of random motion of identical objects approximated by a sphere of radius $a$ – inside a sphere of radius $R$ and on the surface of a sphere with radius $r < R$. When two or more objects contact each other they merge. Our goal is to compare probabilities of merging events in 2D and 3D cases.

First estimate a probabilbity $P_1$ to observe a single object at a specific location inside the sphere as ratio of the volumes $P_1 = (a/R)^3$. The probability to find two objects simultaneously at the same point is $P_2 = P_1^2 = (a/R)^6$.

For the 2D motion the estimate $p_1 = (a/r)^2$ and thus $p_2 = p_1^2 = (a/r)^4$. Then the desired nondimensional ratio $\rho$ reads

$$\rho = P_2/p_2 = \frac{a^6}{R^6} \cdot \frac{r^4}{a^4} = \left(\frac{ar^2}{R^3}\right)^2.$$

Choose the value of $r$ as a length scale and introduce two nondimensional quantitites $k = r/a > 1$ and $m = R/r > 1$. Then we obtain

$$\rho = \left(\frac{ar^2}{R^3}\right)^2 = \frac{1}{k^2 m^6} \ll 1.$$

When motion velocities in 2D $v_2$ and 3D $v_3$ differ significantly one has to introduce a correction. As the probability to visit a specific point is linearly proportional to the velocity we have a correction factor $(v_3/v_2)^2 = \nu^2$ and the ratio reads

$$\rho = \frac{\nu^2}{k^2 m^6}.$$
